# Profiling of Nascent Lariat Intermediates Reveals Key Genetic Determinants of the Timing of Human Co-transcriptional Splicing

**DOI:** 10.1101/2021.10.18.464728

**Authors:** Yi Zeng, Huilin Zeng, Benjamin J Fair, Aiswarya Krishnamohan, Yichen Hou, Johnathon M Hall, Alexander J Ruthenburg, Yang I Li, Jonathan P Staley

## Abstract

As splicing is intimately coupled with transcription, understanding splicing mechanisms requires an understanding of splicing timing, which is currently limited. Here, we developed CoLa-seq (<u>co</u>-transcriptional <u>la</u>riat <u>seq</u>uencing), a genomic assay that reports splicing timing relative to transcription through analysis of nascent lariat intermediates. In human cells, we mapped 165,282 branch points and characterized splicing timing for over 70,000 introns. Splicing timing varies dramatically across introns, with regulated introns splicing later than constitutive introns. Machine learning-based modeling revealed genetic elements predictive of splicing timing, notably the polypyrimidine tract, intron length, and regional GC content, which illustrate the significance of the broader genomic context of an intron and the impact of co-transcriptional splicing. The importance of the splicing factor U2AF in early splicing rationalizes surprising observations that most introns can splice independent of exon definition. Together, these findings establish a critical framework for investigating the mechanisms and regulation of co-transcriptional splicing.

**Highlights:**

1. CoLa-seq enables cell-type specific, genome-wide branch point annotation with unprecedented efficiency.
2. CoLa-seq captures co-transcriptional splicing for tens of thousands of introns and reveals splicing timing varies dramatically across introns.
3. Modeling uncovers key genetic determinants of splicing timing, most notably regional GC content, intron length, and the polypyrimidine tract, the binding site for U2AF2.
4. Early splicing precedes transcription of a downstream 5’ SS and in some cases accessibility of the upstream 3’ SS, precluding exon definition.

## Introduction

As an essential step of gene expression, splicing converts pre-mRNAs to mRNAs by removing introns to ligate flanking exons. Splicing is catalyzed by the spliceosome, a large and dynamic ribonucleoprotein complex that recognizes conserved sequence elements in the intron at the 5’ splice site (5’ SS), the branch point (BP), and the 3’ splice site (3’ SS) (Kastner et al., 2019). The spliceosome catalyzes two sequential transesterification reactions. In the first step, referred to as lariat formation, the 2’ hydroxyl of a conserved BP adenosine attacks the 5’ SS phosphate, forming a free 5’ exon and a branched lariat intermediate, characterized by a 2’-5’ phosphodiester linkage between the BP and the 5’ SS. In the second step, referred to as exon ligation, the 3’ hydroxyl of the free 5’ exon attacks the 3’ SS phosphate, ligating the exons and excising the lariat intron. The fidelity of these two steps is critical to human health (Manning and Cooper, 2017). While splicing *in vitro* occurs in the absence of transcription, splicing *in vivo* is intimately coupled with transcription both physically and functionally (Herzel et al., 2017). However, defining co-transcriptional splicing mechanisms remains a challenge in part due to limited insight into the timing of splicing relative to transcription.

In the earliest steps of spliceosome assembly, the 5’ SS is recognized by the U1 small ribonucleoprotein particle (snRNP), the branch point is recognized by SF1, and the poly-pyrimidine tract (PPT) and 3’ splice site AG dinucleotide are recognized by the U2AF2 and U2AF1 components of the heterodimeric U2AF, respectively. U2AF then enables the U2 snRNP to replace SF1 at the BP (Chen and Manley, 2009). Due to generally limited information encoded at the 5’ and 3’ ends of the intron, early intron recognition is thought to require cooperative binding of U1 snRNP and U2AF to an intron (Hollander et al., 2016), in one of two ways. First, for short introns, most common in lower eukaryotes, U1 and U2AF cooperate across an intron in a process called intron definition. Second, for longer introns, separated by short exons, as is most common in higher eukaryotes, U1 snRNP and U2AF are thought to instead cooperate across an exon in a process called exon definition; in this case, U1 snRNP cooperates from the 5’ SS that is downstream of the intron to be spliced. This cross-exon interaction can be stabilized by additional splicing factors, such as SR proteins, which commonly bind to exonic sequences (Wu and Maniatis, 1993). Because long introns and short exons predominate in human genes, exon definition is thought to be the major mode of spliceosome assembly in humans. Indeed, mutations of splice sites often lead to exon skipping rather than intron retention (Berget, 1995; Hollander et al., 2016). However, recent genomic studies have documented splicing before transcription of the downstream 5’ splice site (Drexler et al., 2020; Reimer et al., 2021; Sousa-Luís et al., 2021); it remains unclear how prevalent such splicing is in higher eukaryotes. Further, the importance of intron and exon length relative to other genetic features in splicing timing remains to be defined.

Though BP selection can impact splice site selection, and mutations altering BP selection induce human disease (Darman et al., 2015), BP usage, in comparison to 5’ and 3’ splice site usage, is poorly defined in humans and many other organisms, because of its degenerate sequence motif and the low abundance of lariat species required to map BP location. To annotate BPs and consequently to enable studies of BP usage and its impact on splice site selection, recent large-scale studies have mined RNA-seq datasets for the incidentally-captured, inverted reads that reveal juxtaposition of the 5’ SS with the BP. However, these reads are extremely rare, found in only ∼1 in a million reads (Taggart et al., 2012). Although the low-abundance of these reads can be partially mitigated by mining thousands of RNA-seq libraries across different tissues and cell types (Pineda and Bradley, 2018; Taggart et al., 2017), such an approach precludes studies of differential BP usage between different conditions, cell types, and tissue types. Alternatively, lariat RNAs have been enriched by degrading linear RNAs using the 3’-5’ exonuclease RNase R; stabilizing lariat RNA with the knockdown of Dbr1, which debranches excised lariat introns for degradation; isolating spliceosomes; and targeting introns for capture (Briese et al., 2019; Chen et al., 2018b; Mercer et al., 2015; Qin et al., 2016; Wan et al., 2021). However, these methods are biased toward subpopulations of lariat RNAs, difficult to implement generally, or restrictive in that they uncouple the BP from the 3’ SS.

Studies have indicated that the majority of splicing (>75%) occurs while RNA polymerase II (RNAP II) synthesizes RNA (Beyer and Osheim, 1988; Herzel et al., 2017; Kessler et al., 1993; Tilgner et al., 2012). Thus, during co-transcriptional splicing, at any given moment, only a subset of splice sites and regulatory elements of a nascent transcript are available to the spliceosome and *trans*-acting splicing factors. As a result, splicing outcome, including splice site selection, critically depends on splicing timing, which we define here as reflecting the interplay between transcription speed and splicing speed (Braunschweig et al., 2013; Herzel et al., 2017). For example, slower transcription has been implicated in favoring earlier splicing timing, with respect to RNAP II position, as well as the upstream of two competing 3’ SSs, whereas faster transcription has been implicated in favoring later splicing timing and a downstream, stronger 3’ SS (Hollander et al., 2016; de la Mata et al., 2003). Further, faster splicing favors in-order splicing in which introns are spliced in the order transcribed, whereas slower splicing permits out-of-order splicing, which has been observed frequently (Kessler et al., 1993; Kim et al., 2017); notably, out-of-order splicing can impact the outcome of alternative splicing by changing the proximity of regulatory RNA elements in downstream regions (Nasim et al., 1990; Takahara et al., 2002). Therefore, defining the timing of co-transcriptional splicing and the *cis*-regulatory sequences and *trans*-acting factors that govern timing is paramount to define how splice sites are selected and regulated during co-transcriptional splicing.

Recent genome-wide studies implementing distinct methodologies in a number of organisms have begun to reveal the timing of co-transcriptional splicing (Wachutka et al., 2019; Drexler et al., 2020; Wan et al., 2021; Carrillo Oesterreich et al., 2016; Herzel et al., 2018; Sousa-Luís et al., 2021; Reimer et al., 2021). For example, some studies in mammals and lower eukaryotes have indicated that splicing can be fast, occurring immediately after transcription of a 3’ SS (Carrillo Oesterreich et al., 2016; Drexler et al., 2020; Herzel et al., 2018; Reimer et al., 2021; Sousa-Luís et al., 2021). Other studies also in mammals, though, imply slower timings for splicing (Drexler et al., 2020; Sousa-Luís et al., 2021; Wachutka et al., 2019; Wan et al., 2021). These differences could reflect a stochasticity of splicing timing or the impact of genetic factors on splicing timing. These differences could also reflect methodological shortcomings, which include mRNA contamination during chromatin isolation, limited depth of long-read sequencing, limited sampling for live imaging, as well as the preferential enrichment of U-rich transcripts or the perturbation of splicing kinetics when metabolically labeling nascent RNA (Testa et al., 1999). Further, due to limited sequencing depth, current methods have only provided an analysis of global splicing timing and have not yet yielded insights into features that predict splicing timing for individual introns.

In this study, we present a novel transcriptome-scale approach, called CoLa-seq (<u>co</u>-transcriptional <u>la</u>riat <u>seq</u>uencing), to study the dynamics and regulation of human co-transcriptional splicing at single nucleotide resolution. CoLa-seq captures excised lariat introns (ELIs), which encode BP usage and 3’ SS usage, and nascent lariat intermediates (NLIs), which encode BP usage and RNAP II position at the time of splicing during transcription. CoLa-seq enabled mapping of 165,282 BPs with a thousand-fold increase in efficiency compared to previous methods and revealed coupling between BP choice and 3’ SS selection. We found that splicing timing, measured as the distance from the 3’ SS to RNAP II at the time of lariat formation, varies over five orders of magnitude. Intriguingly, we found that a majority of introns can splice independent of exon definition, that some introns can even splice before the 3’ splice site emerges from RNAP II, and that alternatively spliced introns splice later than constitutive introns. Most importantly, we discovered multiple features that predict splicing timing, most notably U2AF binding, intron length, and regional GC content. Overall, CoLa-seq establishes an approach and a framework to investigate the modulation of BP usage and the regulation of co-transcriptional splicing.

## Results

### A strategy to capture co-transcriptional lariat intermediates and excised lariat introns

To investigate the timing of lariat formation, relative to the position of elongating RNAP II, as well as to define BP usage, genome-wide in humans, we developed CoLa-seq to capture and sequence the tails of NLIs (**Fig. 1A, B**). In reporting the timing of co-transcriptional lariat formation, which we hereafter refer to as splicing timing, NLIs provide several key advantages over nascent linear transcripts (unspliced and spliced transcripts). First, unlike nascent linear transcripts, NLIs indicate inherently that splicing is in progress. Specifically, whereas the 3’ end of the NLI tail reports on the position of RNAP II, the 5’ end of the tail reveals that the RNA is branched and at the intermediate stage of splicing between the first and second chemical step. Second, NLIs offer a unique opportunity to efficiently define BP usage, with implications for 3’ SS selection. Third, the unique chemical nature of the BP allows for the enzymatic enrichment of NLIs, relative to linear transcripts, and therefore the potential to capture co-transcriptional lariat formation broadly in humans, where the genome encodes >375,000 introns (Piovesan et al., 2016).

**Figure 1.**
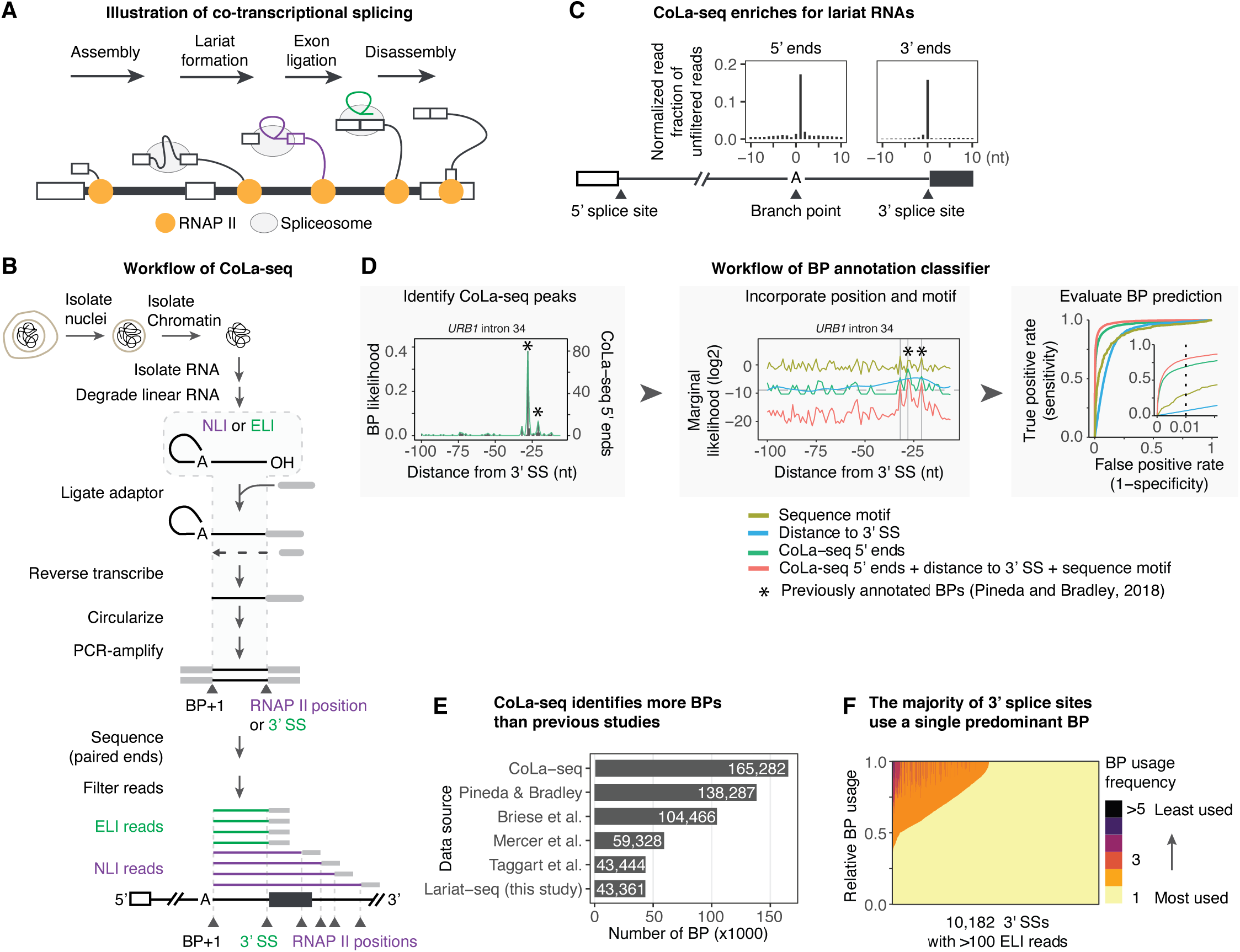
Workflow of CoLa-seq and efficient genome-wide BP mapping. **A.** Illustration of co-transcriptional splicing. The nascent lariat intermediate is colored in purple; the excised lariat intron is colored in green. **B.** Workflow of CoLa-seq. NLI, nascent lariat intermediate; ELI, excised lariat intron; BP, branch point; 3’ SS, 3’ splice site; RNAP II, RNA polymerase II. **C.** Enrichment for lariat tails. Bar plots show alignment of all 5’ ends of reads, relative to BPs, or all 3’ ends, relative to 3’ splice sites, in a 20 nt window. The left meta-plot illustrates that the 5’ ends of reads peaked 1 nt downstream of annotated BPs; the right meta-plot illustrates that the 3’ ends of reads peaked at the last nucleotide of introns. Uniquely mapped, deduplicated reads, before filtering for ELI or NLI reads, are shown, after local normalization. Only constitutive 3’ SSs were used. **D.** Workflow for BP annotation. (**Left panel**) A 100-nt region upstream of a 3’ SS in *URB1* is shown (chr21:32320645−32320740), with previously annotated BPs marked by asterisks (Pineda and Bradley, 2018), as an example illustrating the conversion of the 5’ end of a CoLa-seq read to a likelihood that a 5’ end corresponds to a BP. CoLa-seq reads were shifted 5’ by one nucleotide. (**Middle panel**) The same region of *URB1* is shown as an example illustrating BP likelihoods based on CoLa-seq 5’ ends, similarity to the BP sequence motif (Pineda and Bradley, 2018), and distance to the 3’ SS – alone or as a composite of all three. Composite likelihood peaks greater than a threshold (dotted line) are classified as BPs. In this example, three BPs are identified, as marked by three vertical lines, including two previously identified BPs marked by asterisks. (**Right panel**) The receiver operating characteristic (ROC) curve quantifies the ability of the likelihood functions to identify a set of gold-standard BPs identified in both a previous study (Pineda and Bradley, 2018) and by lariat-seq (**Table S1**). (Inset) the threshold for BP peak calling (vertical dotted line) was determined by considering the sensitivity and specificity of the composite likelihood function. **E.** The bar plot illustrates the number of BPs identified by CoLa-seq, lariat-seq, and previous studies. **F.** The relative usage of CoLa-seq-annotated BPs for each 3’ SS with >100 filtered, ELI and/or NLI reads are shown as a stacked bar graph. Over half of these 3’ SS couple to only one annotated BP. More rarely, 3’ SSs couple to two or more BPs at approximately equal frequency.

To capture NLI tails, we first extracted chromatin-associated RNAs via subcellular fractionation (Pandya-Jones and Black, 2009; Werner and Ruthenburg, 2015; Wuarin and Schibler, 1994). Then, we enriched for NLIs by degrading linear RNAs using the decapping enzyme, RppH, and the 5’-to-3’ exoribonuclease, XRN-1 (**Fig. 1B**). Next, to capture the RNAP II position of individual NLIs, we ligated a UMI (unique molecular identifier)-containing adaptor to the 3’ ends of the isolated RNAs. Then, to capture the BP of the NLI, the molecular signature of this species, we incubated the ligated RNAs with primer and reverse transcriptase, which stops cDNA synthesis at the 2’-5’ linkage, one nucleotide downstream of the BP adenosine. Afterwards, we performed cDNA circularization and PCR amplification. Because ∼90% of BPs reside within the last 50 nts of an intron, and splicing can occur as soon as an intron is transcribed (Reimer et al., 2021; Sousa-Luís et al., 2021), we reasoned that NLI tail-derived fragments are short enough to be assayed by paired-end, short-read sequencing, which allowed for greater depth than longer-read sequencing methodologies. Further, despite our reliance on short reads, this strategy reported on co-transcriptional lariat formation events in which RNAP II is located at varying distances from the 3’ SS (see below). Thus, a CoLa-seq read corresponds to the tail of a given NLI and reveals both the BP location at its 5’ end and the position of RNAP II at its 3’ end (**Fig. 1A, B**). In addition to NLIs, CoLa-seq captures the tails of ELIs, because an ELI remains associated with chromatin at least until the spliceosome releases nascent mRNA, and an ELI tail also has a branch at the 5’ end and a free 3’ hydroxyl at the 3’ end (**Fig. 1A**). Like the 5’ end of a NLI read, the 5’ end of an ELI read reveals the BP location, but its 3’ end reveals a 3’ SS, in the same molecule, thereby coupling BP usage to 3’ SS usage (**Fig. 1A, B**).

In a first application of CoLa-seq, we prepared two biological replicate libraries from human K562 cells (**Fig. S1A**). After deduplicating uniquely mapped CoLa-seq reads, we combined the reads for downstream analyses. Verifying that CoLa-seq reads were enriched for lariat RNA species, the 5’ ends of CoLa-seq reads peaked 1 nt downstream of annotated BPs (**Fig. 1C**; Pineda and Bradley, 2018). Further, the 3’ ends of CoLa-seq reads peaked at the last nucleotide of annotated 3’ SSs (**Fig. 1C**), indicating the enrichment of ELIs among CoLa-seq reads. Importantly, the enrichment at both peaks decreased drastically in cells treated with either the splicing inhibitor pladienolide B or the transcription elongation inhibitor flavopiridol (**Fig. S1B, C**), indicating that these reads are dependent on splicing and transcription.

### CoLa-seq efficiently defines BP usage genome-wide

For CoLa-seq reads whose 5’ ends are located within 100 nts upstream of a 3’ SS in genes expressed in K562 cells (see Methods), only 24% of their 5’ ends are mapped to an annotated BP (**Fig. S2A**). Although reads not mapping to annotated BPs may reflect non-lariat RNA species, many CoLa-seq reads may map to unannotated BPs, given that BP annotations are far from complete (Briese et al., 2019; Mercer et al., 2015; Pineda and Bradley, 2018; Taggart et al., 2017). Thus, to comprehensively map BPs and to sensitively identify reads as deriving from lariat RNAs (NLIs or ELIs), we developed a computational pipeline to systematically identify BPs using CoLa-seq data.

First, we identified reads with a 5’ end within 100 nts upstream of an annotated 3’ SS and classified them as putative NLI reads if they traversed a 3’ SS or as putative ELI reads if they ended at a 3’ SS. Remarkably, through this simple classification, 5’ ends of these two classes of putative lariat RNA-derived reads already resemble annotated BPs in terms of their sequence context (YUNAY) and distance to the 3’ SS (**Fig. S2B**)(Gao et al., 2008; Mercer et al., 2015; Pineda and Bradley, 2018), especially for the 5’ ends of putative ELI reads. These observations suggest that a substantial fraction of putative NLI reads derived from authentic NLIs and that most putative ELI reads derived from authentic ELIs.

To further refine BP mapping, we built a classifier that integrates CoLa-seq data (read coverage of 5’ ends) along with previously defined BP motif and position information (Pineda and Bradley, 2018) to compute a composite BP likelihood for each nucleotide (**Fig. 1D**; see Methods). We then evaluated the BP calling accuracy of the composite BP-likelihood score using a set of gold-standard BPs observed both in a previous large-scale study (Pineda and Bradley, 2018) and our own analysis of K562 cells by lariat-seq (**Table S1**), a direct but inefficient method for identifying BPs (Mercer et al., 2015). Notably, our classifier was able to identify BPs with high accuracy using information from CoLa-seq data alone (AUC=0.96), and the accuracy of our classifier further improved by integrating BP motif and positional information (AUC=0.98; **Fig. 1D**). Using a score threshold that corresponds to a 1% false positive rate (99% sensitivity), our classifier recalled 69% of the gold-standard BPs with CoLa-seq data alone, and 80% of the gold-standard BPs when combining CoLa-seq data with BP motif and positional information (**Fig. 1D**).

With a 1% false positive rate, our classifier identified 165,282 high-quality BPs (**Fig. 1E**; **Fig. S2C, D**; **Table S2**), including 750 U12-type BPs (**Fig. S2E**) and BPs associated with recursive splicing (e.g., **Fig. S2F**). These high-quality BPs showed expected characteristics (**Fig. S2D**); indeed, 14,060 BPs were also observed by lariat-seq (**Fig. S2G**). The large number of BPs identified using our classifier is particularly remarkable considering that our classifier identified more BPs than the most recent large-scale study (**Fig. 1E**), which analyzed 1,000 times as many reads pooled across 17,164 RNA-seq libraries from 65 studies of diverse tissues and cell-types (Pineda and Bradley, 2018). Importantly, the CoLa-seq BPs overlap with previously, experimentally identified BPs to a similar extent as this large-scale study (**Fig. S2G**), indicating that the CoLa-seq BP annotations are of similar high quality. Of the CoLa-seq BPs, 58% (88,881) were novel (**Fig. S2D**) in comparison to BPs previously identified by four large scale studies (Briese et al., 2019; Mercer et al., 2015; Pineda and Bradley, 2018; Taggart et al., 2017), underscoring that BP annotations remain unsaturated. Of these novel BPs, we confirmed 5,733 by lariat-seq (**Table S1**). Previously annotated BPs that were not identified by CoLa-seq are enriched in genes lowly or not expressed in K562 cells (**Fig. S2H**). Together, these observations highlight the utility of Cola-seq in mapping BPs in a cell-type specific manner.

Consistent with previous findings (Pineda and Bradley, 2018), many constitutive 3’ SSs (32%) were associated with more than one BP (**Fig. S2I**). Intriguingly, while a subset of these 3’ SSs used two or more BPs at relatively similar frequencies, most 3’ SSs used a single dominant BP (**Fig. 1F**). Although our classifier may underestimate BP usage, if we more broadly define BP usage based simply on the 5’ end coverage of putative ELI reads (**Fig. S2B**), then we again observed that many 3’ SSs used 3 or more BPs at similar frequencies but more than 50% of the 3’ SSs used predominantly a single BP over 50% of the time (**Fig. S2J**). Thus, in a single cell type a single BP dominates in the majority of introns.

### Excised lariat introns reveal coupling between BP and 3’ SS usage

We first used our BP annotation to filter for ELI reads. As a result, we obtained filtered ELI reads corresponding to 75.2% of putative ELI reads, a more than two-fold increase compared to the number of ELI reads filtered using the largest previously annotated BP set (**Fig. S2A**). As expected, ELI reads derived primarily from protein-coding genes and showed expected BP characteristics (**Fig. 2A**). Additionally, the number of ELI reads for each intron showed strong reproducibility between biological replicates (Pearson’s *r* = 0.98; **Fig. 2B**). In total, we detected 2,574,753 ELI reads associated with 57,309 3’ SSs (≥ 3 ELI reads) (**Fig. 2C**), more than 20 unique ELI reads in 23,272 introns (**Fig. 2C**), and ELI reads in every intron of the major isoform for 1,432 genes. To the best of our knowledge, these reads represent the first genome-wide dataset of ELIs in humans.

**Figure 2.**
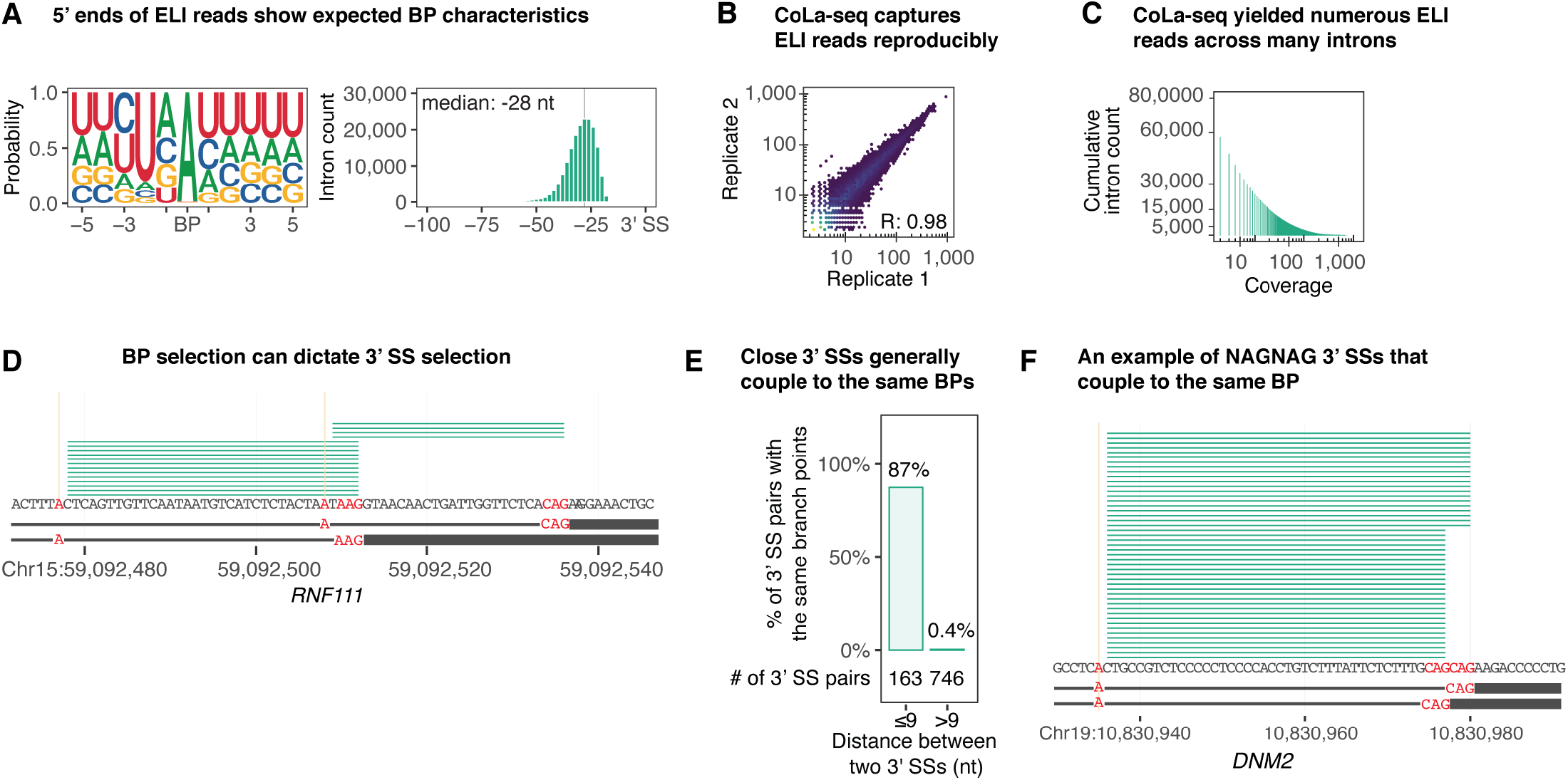
CoLa-seq captures excised lariat introns, revealing BP-3’ SS coupling. **A.** The seqlogo plot (left panel) illustrates the BP motif derived from ELI reads, which matches the YUNAY consensus (Gao et al., 2008). The histogram (right panel) illustrates the distribution of BP-3’ SS distances of ELI reads. Median distances are similar to a reported median distance of 31 nts (Pineda and Bradley, 2018). **B.** The scatter plot compares the number of ELI reads (≥3) associated with a given 3’ SS from two biological replicates. Pearson’s correlation coefficient (R) is shown. **C.** The cumulative bar plot shows cumulative intron counts as a function of the threshold of ELI read coverage. **D.** ELI reads are visualized for intron 12 of *RNF111* and indicate that two isoforms distinguished by alternative 3’ SSs derive from alternative BP usage. **E.** The bar plot illustrates the percentage (labeled on top of each bar) of alternative 3’ SS pairs that use the same set of BPs when the 3’ SSs are within 9 nts of or more than 9 nts away from one another. The number of 3’ SS pairs is shown at the bottom of each bar. **F.** ELI reads are visualized for intron 20 of *DNM2*, which utilizes two adjacent NAGNAG 3’ splice sites but one BP.

Since ELI reads capture both the BP and the 3’ SS of the same molecule, we used ELI reads to examine the functional coupling, or lack thereof, between BP selection and 3’ SS selection. ELI reads revealed cases of strict coupling between BP choice and 3’ SS choice. For example, for two alternative 3’ SSs in the 12th intron of *RNF111*, the upstream 3’ SS always coupled to an upstream BP, whereas the downstream 3’ SS always coupled to a downstream BP (**Fig. 2D**); in this case, the downstream 3’ SS is likely too far (59 nts) from the upstream BP to allow for efficient coupling, and the upstream 3’ SS is too close (3 nts) to the downstream BP to allow for their coupling. Although recognition of the 3’ SS by U2AF can impact BP selection prior to lariat formation, in this example BP selection and lariat formation must ultimately determine 3’ SS selection. On the other hand, when the distance between the BP and the 3’ SS is not restrictive, ELI reads revealed that alternative 3’ SSs that are near to one another are often chosen independently of the BP. Specifically, when pairs of competing 3’ SSs were within 9 nts, 87% of these pairs used the same set of BPs (**Fig. 2E, F**). Thus, by capturing both BP and 3’ SS information in a coupled manner, ELI reads directly reveal whether or not lariat formation restricts 3’ SS selection.

### Nascent lariat intermediates reveal in-order, out-of-order, and concurrent splicing

Next, we used our BP annotations to filter for NLI reads. As a result, we obtained filtered NLI reads corresponding to 38.5% of putative NLI reads, a nearly 2-fold increase compared to the number of NLI reads filtered using the largest previously annotated BP set (**Fig. S2A**). Similar to ELI reads (**Fig. 2A-C**), NLI reads derived primarily from protein-coding genes and showed expected BP characteristics (**Fig. 3A**), and their numbers showed strong reproducibility between biological replicates (Pearson’s *r* = 0.97; **Fig. 3B**). Importantly, despite size-bias in the capture of ELI cDNAs (**Fig. S2K, L**), 69% of the NLI-associated BPs overlapped with ELI-associated BPs (**Fig. S2M**), and NLI and ELI read numbers positively correlated (Pearson’s *r* = 0.32), consistent with the functional conversion of NLI species to ELI species. Aside from host gene expression differences (**Fig. S2H**), we found no major differences between introns that yielded CoLa-seq reads and introns that did not (**Fig. S2N**). In total, we detected 2,201,036 NLI reads associated with 73,562 3’ SSs (≥ 3 ELI reads) (**Fig. 3C**), more than 20 unique NLI reads in 25,476 introns (**Fig. 3C**), and NLI reads in every intron of the major isoform for 2,941 genes. To our knowledge, these NLI reads represent the first major dataset of NLIs in any organism. Importantly, the depth and breadth of these datasets allowed us to study co-transcriptional splicing at the level of individual introns, which has been difficult with lower-depth, long-read sequencing methods.

**Figure 3.**
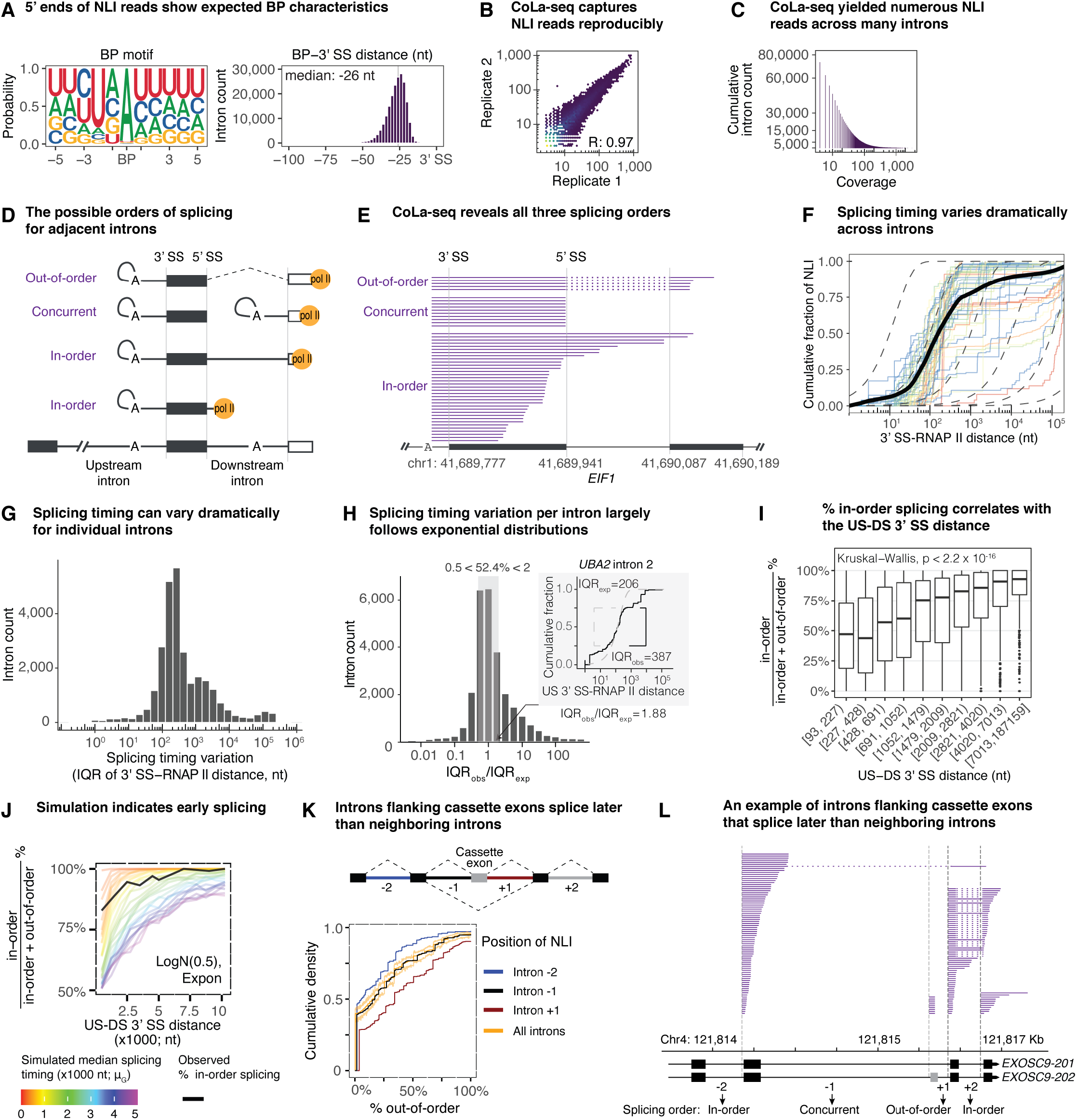
Nascent lariat intermediates capture in-order, out-of-order, and concurrent splicing, revealing dramatic variation in splicing timing. **A.** The seqlogo plot (left panel) illustrates the BP motif derived from NLI reads, which matches the YUNAY consensus (Gao et al., 2008). The histogram (right panel) illustrates the distribution of BP-3’ SS distances of NLI reads. **B.** The scatter plot compares the number of NLI reads (≥ 3) associated with a given 3’ SS from two biological replicates. Pearson’s correlation coefficient (R) is shown. **C.** The cumulative bar plot shows cumulative intron counts as a function of the threshold of NLI read coverage. **D.** Schematic of the possible orders of co-transcriptional splicing for adjacent introns, with the upstream intron at the intermediate stage of splicing and the downstream intron at the pre-mRNA, mRNA, or intermediate stage, representing in-order, out-of-order, and concurrent splicing, respectively. The dotted line indicates completed splicing of the downstream intron in the case of out-of-order splicing. **E.** An example of an intron, intron 1 of *EIF1,* that yields NLI reads corresponding to all three possible orders of splicing. **F.** A cumulative plot (solid black line) shows the distribution of 3’ SS-RNAP II distances from all in-order and out-of-order NLI reads. The colored lines show cumulative distributions of 3’ SS-RNAP II distances for fifty randomly sampled introns with >30 NLI reads. For reference, each dotted line is a cumulative distribution for a theoretical exponential distribution with the splicing half-life set to a major tick mark. **G.** The histogram shows the distribution of splicing timing variation per intron. For each intron, its splicing timing variation is defined as the interquartile range (IQR) of its 3’ SS-RNAP II distance distribution from its in-order and out-of-order NLI reads. **H.** The histogram shows the distribution of the ratio of the observed splicing variation (interquartile range, IQR_obs_) for an intron to the theoretical splicing variation (IQR_exp_) for the intron if the splicing half-life (median 3’ SS-RNAP II distance) were the same but splicing probability (i.e., 3’ SS-RNAP II distance distribution) decayed as an exponential function. The shaded area marks introns (52.4% of total) with a ratio of IQR_obs_/IQR_exp_ between 0.5 and 2. The inset shows the distribution of 3’ SS-RNAP II distance for in-order and out-of-order NLI reads observed in intron 2 of *UBA2*, highlighting the IQR_obs_ (solid black line) and the IQR_exp_ (dashed gray line). **I.** The box plot visualizes the correlation between % in-order splicing and the distance between the upstream 3’ SS and the downstream 3’ SS (US-DS 3’ SS distance). The % in-order splicing is calculated as (in-order NLI reads) / (in-order NLI reads + out-of-order NLI reads) x 100. Only size-corrected in-order and out-of-order NLI reads are used (see text). Introns were divided into 10 groups, with the same number of introns, based on the US-DS 3’ SS distance. **J.** Simulations indicate early splicing. Observed data (black) for % in-order splicing as a function of the US-DS 3’ SS distance is most consistent with simulations (colored) wherein half of splicing occurs when RNAP II is within 1000 nt downstream of the 3’ SS (see Methods for detail; see also **Fig. S3K-M**). Briefly, % in-order splicing was simulated under various scenarios where the splicing rates of individual introns in adjacent intron pairs are modeled as exponential distributions, expon(1/μ_1_) and expon(1/μ_2_), where μ_1_ and μ_2_ are drawn from a log-normal distribution, logN(μ_G_, σ^2^) with σ^2^ being 0.5; μ_G_ can be interpreted as the median splicing half-life across introns genome-wide in terms of 3’ SS to RNAP II distance. Simulated % in-order splicing for values of μ_G_ between 0 and 5kb are shown as colored lines. **K.** Introns flanking cassette exons splice later than their neighboring constitutive introns. The top panel illustrates the general gene structure of cassette exons. The cumulative plot in the bottom panel illustrates the % out-of-order splicing for the indicated introns. The vast majority of cassette exons inspected were predominantly included, such that NLIs associated with the BP of intron +1 derived from excision of intron +1, rather than excision of intron −1, intron +1, and the intervening exon. **L.** An example illustrating the later splicing of introns flanking a cassette exon. NLI reads from introns from *EXOSC9* are visualized. The predominant order of splicing is indicated for each intron. The cassette exon is labeled in gray. As in **K**, the cassette exon was predominantly included, such that the NLIs associated with the BP of intron +1 derived from excision of intron +1.

Investigating NLI reads, we first interrogated the order of splicing of adjacent intron pairs by defining the splicing status of a downstream intron relative to an upstream intron in which a NLI branch was detected. We observed three types of NLI reads, reflecting three classes of splicing order: (i) in-order splicing, (ii) out-of-order splicing, and, unexpectedly, (iii) concurrent splicing (**Fig. 3D, E**; **Fig. S3A**). Importantly, the relative proportion of reads supporting each class of splicing order at each intron pair is highly reproducible between replicates (**Fig. S3B**). In-order splicing represents the earliest splicing timing for an upstream intron and is indicated by NLI reads that reflect either an incompletely transcribed or unspliced downstream intron (**Fig. 3D, E**). By contrast, out-of-order splicing represents the latest splicing timing for a given upstream intron and is indicated by NLI reads that reflect a transcribed and spliced downstream intron (**Fig. 3D, E**); such out-of-order NLIs can also reflect further transcription and splicing of introns even farther downstream. In our dataset, we observed in-order splicing in 93% of introns and out-of-order splicing in 46% of introns (**Fig. S3C**). Notably, although CoLa-seq is biased toward shorter reads and consequently early splicing events (**Fig. S3D, E**), we observed that 7% of introns spliced predominantly out-of-order.

Strikingly, we also observed concurrent splicing, in which two adjacent introns splice at the same time. To our knowledge, concurrent splicing has only been observed *in vitro* and for a synthetic substrate (Christofori et al., 1987). For a given nascent intron, concurrent splicing represents an intermediate timing of splicing and is indicated by NLI reads that end at the last nucleotide of the intervening exon (**Fig. S3G**), reflecting cleavage of the downstream 5’ SS. Thus, we infer that these NLI reads indicate that the downstream intron is transcribed and, like the upstream intron, at the intermediate stage of splicing (**Fig. 3D, E**); indeed, 64% of the corresponding downstream introns also yielded NLIs (≥5 reads). In our dataset, we observed concurrent splicing in 90% of introns (**Fig. S3C**). In the cell, a concurrent splicing NLI will precede an out-of-order NLI, but CoLa-seq captures out-of-order splicing in a manner dependent on RNAP II position and the length of the downstream NLI tail, whereas concurrent splicing reports on splicing order independent of RNAP II position and the length of the downstream NLI tail, so concurrent splicing has the potential to report on the latest splicing timings.

### The timing of co-transcriptional splicing varies dramatically across introns

By analyzing NLI-associated RNAP II positions as well as the order of splicing, we found that splicing timing varies dramatically. Specifically, considering that the 3’ SS is required for spliceosome assembly for most introns (Shao et al., 2014), and the 3’ end of in-order and out-of-order NLI reads report on the position of RNAP II when splicing occurs (**Fig. 3D**), we calculated splicing timing by measuring the genomic distance from the 3’ SS to the position of RNAP II (3’ SS-RNAP II distance) for each NLI. Importantly, out-of-order splicing can only occur when RNAP II has transcribed beyond the downstream intron; nevertheless, splicing of the downstream intron and sometimes further introns (**Fig. S3J)** can lead to an NLI with a short tail, so through short-read sequencing, out-of-order reads can report on long 3’ SS-RNAP II distances, representing late splicing. By aggregating both in-order and out-of-order NLI reads across introns, we found that splicing timing varies over five orders of magnitude, from less than 3 nts to 1,115,021 nts (**Fig. 3F**). The median timing for all splicing events was 160 nts from the 3’ SS (**Fig. 3F**; a minimum estimate given short-read length bias), consistent with a recent study in murine cells (Reimer et al., 2021). Notably, in this dataset, 10% of splicing events occurred within 21 nts (see below), and 10% of events occurred beyond 7,470 nts. Similarly, by concurrent splicing, in which the distance from the upstream 3’ SS to the downstream 3’ SS represents a minimum splicing time for the upstream intron (**Fig. 3D**), we observed splicing events when RNAP II has already transcribed the downstream intron, with 10% of splicing events occurring beyond 4.9 kb and the latest event occurring beyond 179 kb (**Fig. S3H**).

The great depth of CoLa-seq also allowed us to assess splicing timing variation at the level of individual introns, considering 17,448 introns with ≥10 in-order or out-of-order NLI reads. We observed that splicing timing for an intron can vary substantially (**Fig. 3G**; **Fig. S3J**), with a median splicing variation (interquartile range, IQR) per intron of 258 nts but with a splicing variation (IQR) larger than 10,000 nts for 6.3% of introns. We note that under simple first-order kinetics with exponential decay for splicing, large splicing variation (e.g., >10kb) is expected for NLIs with long median 3’ SS-RNAP II distances, as we observed for some introns (**Fig. S3I, J**). To examine whether the variation in splicing timing within an intron is consistent with first-order exponential kinetics, we compared the observed splicing variation (IQR_obs_) of each intron to the splicing variation (IQR_exp_) expected for an exponential model with the observed median 3’ SS-RNAP II distance. Importantly, the majority of introns showed splicing variation (IQR_obs_) that is within 2-fold of the expected variation (IQR_exp_; **Fig. 3H**), indicating that simple first-order kinetics can capture the splicing variation of a typical intron. Interestingly, a small fraction of introns (11%) showed splicing timing variation (IQR_obs_) 10 times greater than the expected splicing variation (IQR_exp_), which suggests heterogeneity in splicing mechanisms and kinetics or in transcriptional dynamics; conversely, an even smaller fraction (4.2%) showed splicing variation 10 times narrower than expected, which might reflect deterministic splicing, a tight window of opportunity for splicing, or RNAP II pausing. These variations in splicing have important implications for splicing regulation.

As an alternative, proxy measure of splicing timing, we calculated the distribution of reads between the in-order and out-of-order splicing classes across all detected introns. To eliminate the impact of read size bias toward in-order splicing reads, which can end upstream of the downstream 5’ SS, unlike out-of-order splicing reads, we calculated the proportion of in-order splicing using only in-order and out-of-order NLI reads that extend beyond the downstream 5’ SSs and thus have similar length distributions (**Fig. S3D, E, F**). Notably, the proportion of in-order splicing strongly correlates with the distance between the upstream and downstream 3’ splice sites (US-DS 3’ SS distance; **Fig. 3I**). When the US-DS 3’ SS distance is longer than 2 kb, the median proportion of in-order splicing reaches over 75%. This observation is remarkable as it implies that the large majority of introns splice before RNAP II has transcribed 2 kb downstream of the intron. To investigate how this correlation can inform the timing of splicing, we simulated splicing timings, expressed as 3’ SS-RNAP II distances, and examined how our observed data fit various simulated scenarios. Briefly, we assumed that splicing half-lives (median splicing timings) from introns genome-wide follow a lognormal distribution with some unknown median half-life (**Fig. S3K, L, M**; see Methods). We then simulated the proportion of in-order splicing for adjacent intron pairs by drawing the splicing timing for each intron of the pairs from exponential distributions with different splicing half-lives. As expected, the percentage of in-order splicing increases as the US-DS 3’ SS distance increases for all simulated median splicing half-lives (**Fig. 3J**). For a median US-DS 3’ SS distance of 1.5 kb and a simulated median splicing half-life of ∼5 kb, the simulated percentage of in-order splicing approaches 50%, because with such a long half-life the transcription time for the US-DS 3’ SS distance becomes insignificant and the probability of splicing in-order versus out-of-order becomes equal. However, again with a median US-DS 3’ SS distance of ∼1.5 kb, we observed a higher percentage of in-order splicing (∼70%), implying that the median splicing half-life must be smaller than 5 kb. Indeed, in all simulation scenarios considered, we found that the median splicing half-life must be smaller than 1 kb for the simulated level of in-order splicing to match the high levels of in-order splicing empirically observed by CoLa-seq (**Fig. 3J**; **Fig. S3M**), consistent with the minimum median splicing time of 160 nts derived from 3’ SS-RNAP II distance distributions alone (**Fig. 3F**). Together, our analyses indicate that although the timing of splicing for individual introns can vary by orders of magnitude, co-transcriptional splicing occurs generally before an average-sized downstream intron is fully transcribed.

To corroborate our usage of splicing order as a proxy of splicing timing, we compared the order of splicing for various intron classes. Previous studies have suggested that higher levels of U12, relative to U2, intron-containing pre-mRNAs in cells could be due to either inefficient splicing or slow splicing (Niemelä and Frilander, 2015; Patel et al., 2002; Wachutka et al., 2019). We found that U12 introns yield lower levels of in-order splicing than U2 introns (**Fig. S3N**), providing supporting evidence that U12 introns generally splice later than U2 introns (Wachutka et al., 2019).

Previous findings also suggest that splicing timing of alternatively spliced introns is generally slower than that of constitutively spliced introns (Herzel and Neugebauer, 2015; Pai et al., 2017; Pandya-Jones and Black, 2009). To test for slower splicing, we examined the order of splicing for introns flanking cassette exons and their neighboring introns (**Fig. 3K,** top cartoon panel). Indeed, intron pairs upstream of the cassette exons (−2 and −1 introns) spliced by out-of-order splicing to a lower degree than all intron pairs (predominantly constitutive introns), supporting later splicing of introns immediately upstream of cassette exons (−1 introns), relative to constitutive introns (**Fig. 3K, L**). Further, intron pairs downstream of cassette exons (+1 and + 2 introns) spliced by out-order splicing to a higher degree than all intron pairs, supporting later splicing of introns downstream of cassette exons (+1 introns), relative constitutive introns. Curiously, intron pairs flanking cassette exons (−1 and +1 introns) spliced by out-of-order splicing to a degree similar to all, predominantly constitutive intron pairs (**Fig. 3K**), implying that introns flanking cassette exons splice later to a similar degree. These results are consistent with recent findings that examined the relative order of exon ligation for adjacent introns (Drexler et al., 2020; Kim et al., 2017). Together, the later splicing of U12 introns and cassette-flanking introns corroborate our use of splicing order as a proxy for splicing timing. Further, the later splicing of cassette-flanking introns supports a role for splicing timing variation in regulating alternative splicing.

### Intronic elements, gene architecture, and genomic context predict the order of co-transcriptional splicing

The substantial variation in splicing timing across introns suggested that specific genetic sequences influence splicing timing. To probe for such a genetic basis, we first investigated sequence features that correlate with an order of splicing. We found that introns with high levels of in-order splicing tend to have (i) strong splicing signals, such as a strong PPT reflected by high U content in the BP-3’ SS region; (ii) longer BP-3’ SS distances, exons, downstream introns, and, unexpectedly, upstream introns; and (iii) lower regional GC content (**Fig. S4A**). Because the percentages of in-order splicing and concurrent splicing are inversely correlated (Pearson’s *r* = −0.9), concurrent splicing showed correlations with features that were generally opposite of that for in-order splicing (**Fig. S4A**). Interestingly, while out-of-order splicing showed correlations that parallel concurrent splicing for some features, such as downstream intron length, for other features, such as downstream BP-3’ SS distance, the correlations were opposite (**Fig. S4A**). These observations imply that specific sequence features influence splicing timing. However, we note that many of the correlated features correlate among themselves (e.g., **Fig. S5G**), which complicates the identification of key features that impact splicing timing.

To explore the impact of sequence features on the order of splicing independent of such feature-feature correlations and in a quantitative manner, we trained computational models that predict the percentage of a splicing order, using a machine learning algorithm, XGBoost (Chen and Guestrin, 2016). Briefly, XGBoost uses decision tree ensembles to identify key features, such as sequence elements, that predict a given target variable, such as the percentage of a splicing order. After pre-screening 235 features (**Table S3**; Methods), we selected 22 features for modeling that include specific splicing elements, intron/exon architecture, sequence content, and secondary structure potential. We normalized each class of splicing order relative to the sum of all three classes, and trained models on 23,324 introns for % in-order splicing, % concurrent splicing, and % out-of-order splicing, separately. These models outperformed linear regression-based models by 1.3-to 2.2-fold in terms of predictive ability (**Fig. S4B**). Remarkably, our XGBoost-based models accounted for 40%±1% (std), 34%±1%, and 15%±1% of the overall variance for in-order splicing, concurrent splicing, and out-of-order splicing, respectively.

For each trained model, we calculated the importance of each feature to each prediction using SHAP (SHapley Additive exPlanation; Lundberg and Lee, 2017). For each prediction of a percent order of splicing for an intron pair, SHAP determines the overall importance of each feature by varying the values of a single feature and calculating the change in the model output. Additionally, SHAP breaks down the overall effect of each feature into its main effect, which is independent of other features, and its interaction effect, which is dependent on other feature values. In our models, interaction effects between features were minimal. Importantly, multiple features contribute to the prediction for each intron pair (e.g., **Fig. 4A, B**; **Fig. S4C, D**). Three general classes of elements emerged as predictive for the order of splicing: (i) canonical intronic splicing elements, (ii) the sizes of introns and exons, and (iii) GC content (**Fig. 4A**; **Fig. S4C**). Most of these features are consistent with a role in accelerating or delaying splicing of either the upstream or downstream intron.

**Figure 4.**
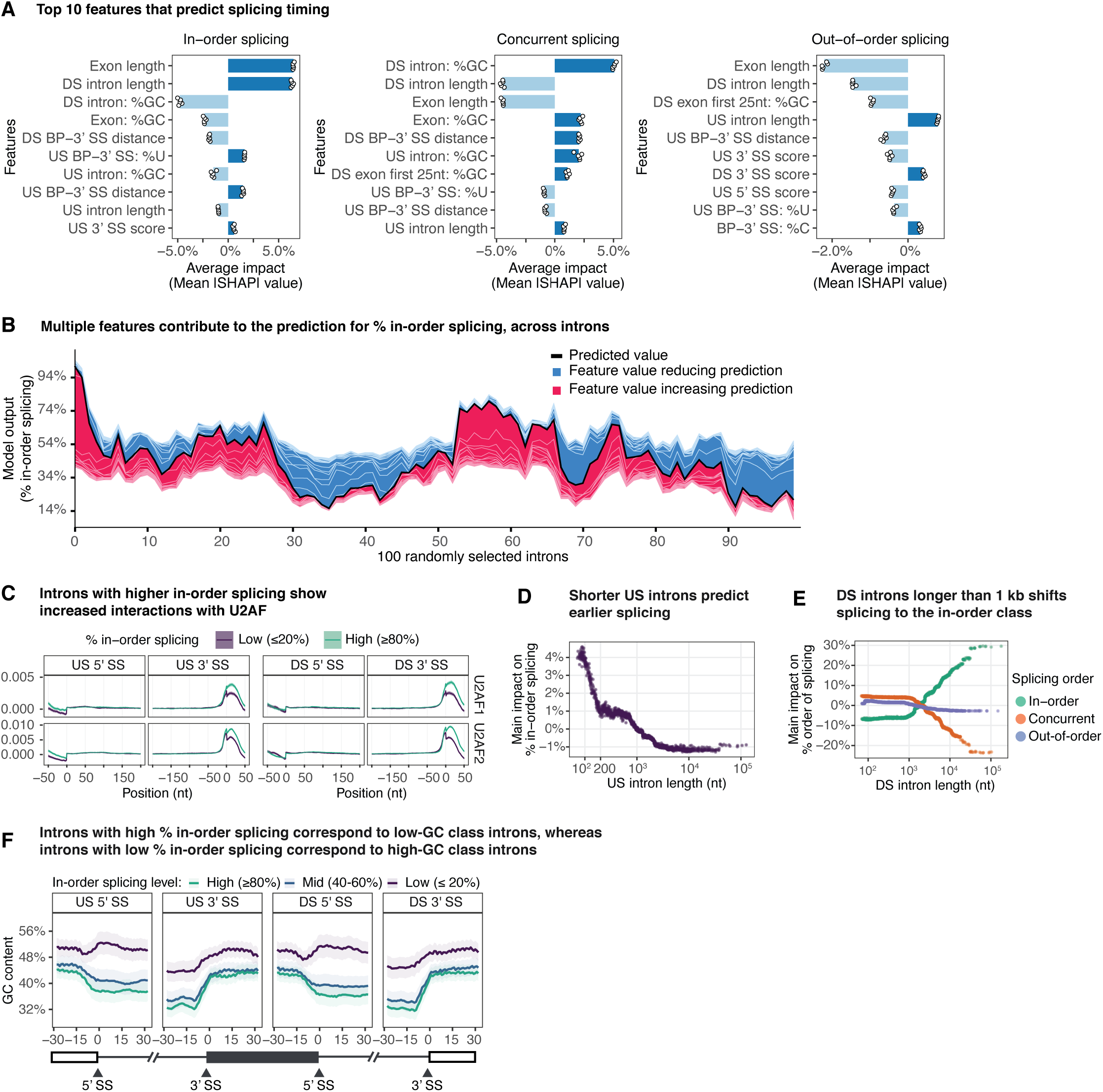
Intronic elements, local gene architecture, and genomic context predicts the order of co-transcriptional splicing. **A.** The average impact (average of absolute SHAP values for each intron) of the top 10 features is visualized for XGBoost models that predict the levels of in-order splicing, concurrent splicing, or out-of-order splicing. The average impact values from 5-fold cross validations are plotted as points and the averages of these values are shown as bars. The sign of the impact was determined by Spearman’s correlation. “US”, upstream intron harboring the NLI; “exon”, exon downstream of NLI; “DS”, intron downstream of NLI. SHAP value is calculated as the impact on the percentage of a splicing order. **B.** The summary force plot visualizes the feature contributions to the prediction of % in-order splicing for 100 randomly sampled introns. The black line plots the final prediction value. Features that increase the predictions from the base value (the % in-order splicing averaged over all introns, which is 44%) are colored in red; features that reduce the predictions from the base value are colored in blue. **C.** Line plots illustrate averaged eCLIP signals (Van Nostrand et al., 2020) of a given splicing factor in the 5’ SS region (50 nts upstream and 150 nts downstream) and the 3’ SS region (150 nts upstream and 50 nts downstream) for introns with either a high or low percentage of in-order splicing. Binding is shown for both the upstream, NLI intron, and the downstream intron. The 95% confidence intervals are shown by the shaded areas. **D.** The scatter plot illustrates the main impact of upstream (US) intron length on the prediction of % in-order splicing, revealing that when shorter than 200 nts, upstream intron length has a strong positive impact. The main impact is measured using SHAP, reveals impact independent of other features, and indicates the change in percentage predicted for in-order splicing at indicated values of upstream intron length. **E.** The scatter plot illustrates the main impact of downstream (DS) intron length on the prediction of % in-order splicing, concurrent splicing, or out-of-order splicing. The main impact values are as described in legend 4D. **F.** Introns with different levels of in-order splicing exhibit different patterns and percentages of GC content across splice sites of the upstream and downstream introns (+/-30 nts). The 95% confidence intervals are shown by the shaded areas.

Supporting the physiological significance of NLI levels observed by CoLa-seq as well as the validity of the models, the BP-3’ SS distance of both the upstream and downstream intron are important in the models, accounting, for example, for 11% of the predictive power of the in-order splicing model, as determined by the sum of the mean absolute SHAP value for each feature (**Fig. 4A**; **Fig. S4C**). Specifically, a longer downstream BP-3’ SS distance predicts higher levels of concurrent splicing but lower levels of out-of-order splicing (**Fig. S5A**), indicating higher levels of lariat intermediate for the downstream intron, as expected for a slower rate of conversion from lariat intermediate to mRNA when the BP-3’ SS distance is longer (Meyer et al., 2011). Similarly, a weaker downstream 3’ SS predicts higher levels of concurrent splicing and lower levels of out-of-order splicing, as expected (**Fig. S5A**).

As anticipated, the strength of intronic splicing elements generally contributed to predicting the percentage of an order of splicing; these elements together accounted, for example, for 13% of the predictive power of the in-order splicing model. Such intronic elements include features related to the 5’ SS and the 3’ SS (**Fig. 4A**; **Fig. S4C**), but, interestingly, the most important feature is U content in the upstream BP-3’ SS region, with higher U content predicting an increase in % in-order splicing and a decrease in % concurrent and % out-of-order splicing (**Fig. 4A**). This observation is notable as U content in this region positively correlates with PPT strength (Pearson’s *r* = 0.34). Similarly, the BP-3’ SS distance of the upstream intron positively correlates with PPT strength (Pearson’s *r* = 0.3), and an increase in length similarly predicts an increase in % in-order splicing (**Fig. S5B**). Using k-mer analysis and published eCLIP-seq data (Van Nostrand et al., 2020), we further observed that introns with higher levels of in-order splicing are enriched for U-rich hexamers (**Fig. S5C**) and exhibited increased interactions with the U2AF heterodimer (**Fig. 4C**), suggesting that a strong PPT and correspondingly strong U2AF binding promote in-order and thus early splicing. Of note, U content outperforms PPT strength in the model, implying that U content captures additional genetic features. Indeed, U content anti-correlates with RNA secondary structure potential (Pearson’s *r* = −0.27), which can interfere with 3’ SS recognition (Meyer et al., 2011 and reference cited therein). Interestingly, we observed a greater importance of the BP-3’ SS region than the 5’ SS region on the model output (**Fig. S4C**). This observation suggests that the later transcription of the 3’ end of the intron, relative to the 5’ end, imposes a greater constraint on 3’ SS recognition for early splicing of the upstream intron.

Importantly, we found that the lengths of the upstream intron, the exon, and the downstream intron all contribute to the model predictions. Exon length correlates with concurrent splicing read size and read size is influenced by size bias (**Fig. S3D, E**), and when we correct for size bias, exon length no longer correlates with % in-order splicing (**Fig. S3F, S5D**), so we did not consider exon length further. Upstream and downstream intron lengths accounted for 3% and 21%, respectively, of the predictive power of the in-order splicing model, for example (**Fig. 4A, D, E**). Whereas the simple correlation analysis (above) indicated unintuitively that a shorter upstream intron correlated with lower levels of early in-order splicing (**Fig. S4A**; Sousa-Luís et al., 2021), the model revealed the opposite – that shorter upstream introns predict higher levels of in-order splicing, especially when the upstream intron length is shorter than 200 nts (**Fig. 4D**), as would be expected (see Discussion).

Further, the model revealed that longer downstream introns (>1 kb) predict increasingly higher levels of in-order splicing and lower levels of both concurrent and out-of-order splicing (**Fig. 4E**). This importance of downstream intron length is not due to its impact on splicing of the downstream intron, because the importance of upstream intron length in predicting upstream intron splicing remains constant beyond ∼2 kb (**Fig. 4D**). Consistent with the early median splicing timing implied by our simulations (**Fig. 3J**; **Fig. S3K-M**), these findings imply that for downstream introns longer than 2 kb in-order splicing increases with downstream intron length because the transcription of these downstream introns delays their splicing. Supporting this notion, as downstream intron lengths extend beyond 5 kb, downstream intron length becomes the dominant feature predicting in-order splicing (**Fig. S5E**). Importantly, these findings highlight the co-transcriptional nature of splicing and the importance of transcription time on the outcomes of co-transcriptional splicing (see Discussion).

In addition to the lengths, the GC content of the upstream intron, the exon, and the downstream intron feature prominently in the models, accounting for 12% of the predictive power of the in-order splicing model, for example. In particular, high GC content in all three regions predict low levels of in-order splicing (**Fig. 4A**; **Fig. S5F**). Since the GC content of all three regions strongly correlate (**Fig. S5G)**, their importance in the models suggests that regional GC content bias influences the timing of splicing. Interestingly, such regional GC content bias has been shown to separate introns into two general classes: a higher GC class with shorter introns and similar GC content between exons and introns, and a lower GC class with longer introns and lower GC content in the flanking introns than in the exons (Amit et al., 2012). Consistent with a functional significance of this classification, we found that intron pairs with low levels of in-order splicing associate with higher, uniform GC content between exons and introns, whereas intron pairs with high levels of in-order splicing associate with lower, differential GC content between exons and introns (**Fig. 4F**). Further, consistent with previous findings that regional GC content bias often extends well beyond an intron or even a gene (Amit et al., 2012; Costantini and Musto, 2017; Costantini et al., 2006), we found evidence that the impact of regional GC content bias is evident on the gene level. Compared to introns from different genes, introns from the same genes tend to have similar GC content (p-value < 4.3 x 10^-167^; F-test) and similar splicing timings, as measured by % in-order splicing (p-value < 1.2 x 10^-24^; chi-square test). Given that high GC content increases local secondary structure potential and weakens the strength of splice sites (**Fig. S5H**; Amit et al., 2012), GC content likely impacts splicing timing by affecting the recognition and accessibility of splicing signals across the introns of a gene; indeed, U2AF binding correlates between adjacent introns (**Fig. 4C**). Taken together, our findings demonstrate that the broad genomic context of an intron is a critical determinant of splicing timing.

### 3’ splice site features, gene architecture, and GC content predict the timing of in-order splicing

Given that 93% of introns yielded NLI reads indicative of early, in-order splicing (**Fig. S3C**), we examined the timing of early lariat formation in terms of the 3’ SS-RNAP II distance distribution of in-order NLIs. The median 3’ SS-RNAP II distance of in-order NLIs from each intron is reproducible between replicates (Pearson’s *r* = 0.68). As anticipated (**Fig 3D**, **E**), the 3’ SS-RNAP II distances vary by over hundreds of nucleotides for individual introns and importantly between introns (**Fig. 5A, B, C**; **Fig. S6A, B**), implying a genetic basis for in-order splicing timing. Notably, some NLI 3’ ends are in the region immediately downstream of the 3’ SS, indicative of very early lariat formation (**Fig. 5A, B, C**; **Fig. S6A**).

**Figure 5.**
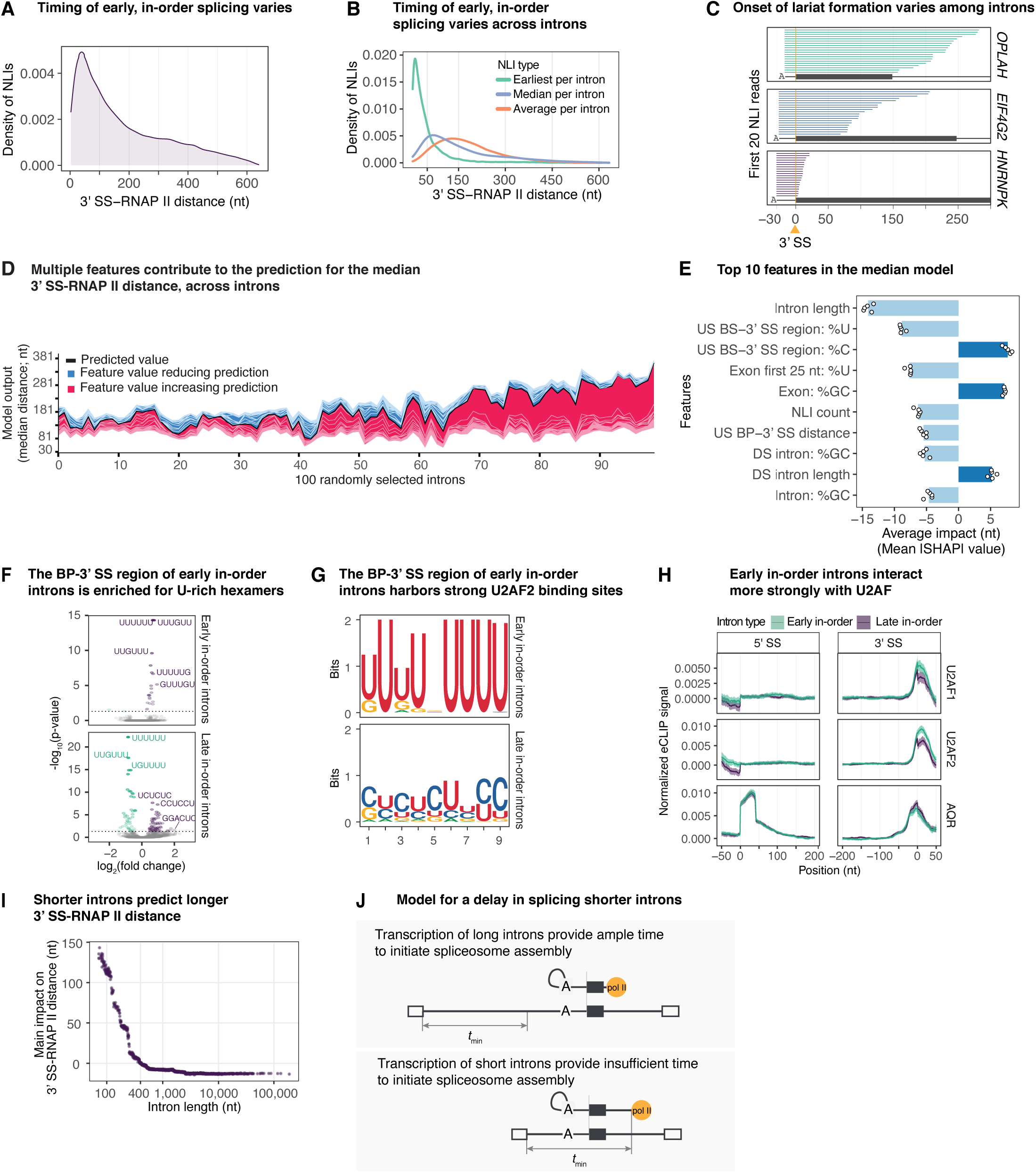
Specific features predict early in-order splicing. **A.** The density plot illustrates the varied, 3’ SS-RNAP II distance distribution of all in-order NLI reads. **B.** The density plot illustrates the distance distribution of the earliest, median, and average 3’ SS-RNAP II distances from each intron. **C.** The first 20 in-order NLI reads are visualized for intron 20 of *OPLAH* (of 56 in-order NLI reads), intron 16 of *EIF4G2* (of 85 in-order NLI reads), and intron 16 of *HNRNPK* (of 440 in-order NLI reads). **D.** The summary force plot visualizes the feature contributions to the prediction of the median 3’ SS-RNAP II distance for 100 randomly sampled introns. The black line plots the final prediction value. Features that increase the predictions from the base value (the average of the median 3’ SS-RNAP II distance across all introns, which is 131 nts) are colored in red; features that reduce the predictions from the base value are colored in blue. **E.** The average impact of the top 10 features for the median 3’ SS-RNAP II distance model is shown. The average impact of a given feature on 3’ SS-RNAP II distance, in units of nucleotides, represents the average of the absolute SHAP values for the indicated feature for each intron. Otherwise, details are as in Fig. 4A. **F.** U-rich hexamers are enriched in the BP-3’ SS region of early in-order introns and depleted in the same region of late in-order introns. Significantly-enriched hexamers are in purple, and significantly-depleted hexamers are in green (Benjamini-Hochberg corrected p-value <0.05). P-values (-log_10_(p-value)) are plotted as a function of the hexamer fold enrichment (log_2_(fold change)). Early in-order introns have ≥70% of 3’ SS-RNAP II distances ≤50 nts; late in-order introns have ≥70% of 3’ SS-RNAP II distances ≥200 nts. **G.** Seqlogos illustrate motifs enriched in the BP-3’ SS region of early or late splicing introns. The motif enriched in early in-order introns corresponds to a strong U2AF2 binding site, whereas the motif enriched in late in-order introns corresponds to a weak U2AF2 binding site (see Methods). **H.** Line plots illustrate averaged eCLIP signals (Van Nostrand et al., 2020) of splicing factors for early or late in-order introns. The eCLIP signal for Aquarius (AQR) is shown as a negative control. Otherwise, details are as in Fig. 4C. **I.** The main impact of intron length on in-order NLI 3’ SS-RNAP II distance (nt) is plotted as a function of 3’ SS-RNAP II distance. The main impact corresponds to the intron length SHAP value, independent of interactions with other features, and measured in nucleotides. Values are derived from the median 3’ SS-RNAP II distance model. **J.** The schematic illustrates that whereas transcription of long introns yields ample time to initiate spliceosome assembly and undergo splicing when RNAP II is just downstream of the 3’ SS, thus yielding nascent lariat intermediates with short 3’ SS-RNAP II distances, transcription of short introns yields insufficient time to initiate spliceosome assembly, and therefore short introns undergo splicing when RNAP II is further downstream of a 3’ SS, thus yielding nascent lariat intermediates with longer 3’ SS-RNAP II distance. The value *t*_min_ indicates the minimum time required for spliceosome assembly and is measured in nucleotides.

To investigate the impact of sequence features on lariat formation timing, we trained two computational models on 14,437 introns, again using XGBoost (Chen and Guestrin, 2016). In the “earliest” model, we used as the target variable the shortest 3’ SS-RNAP II distance for in-order NLIs of a given intron (**Fig. 5B;** median of earliest lengths, 18 nts), because such a length reflects the requirements for each intron to undergo lariat formation at the earliest detected timing; in the “median” model, we used as the target variable the median 3’ SS-RNAP II distance for in-order NLIs of a given intron (**Fig. 5B;** median of median lengths, 107 nts), because such a length reflects the requirements for the population of NLIs from an intron to undergo in-order lariat formation. After screening 235 features (**Table S3;** Methods), we selected 37 features for modeling, including the number of NLI reads per intron to account for sampling bias. The resulting earliest and median models accounted for 31%±2% and 28%±1% of the variance in 3’ SS-RNAP II distance, respectively. To quantify the contribution of each feature to the models, we again used SHAP (Lundberg and Lee, 2017). Both models reveal that multiple features contribute to the prediction of splicing timing for each intron (**Fig. 5D**) and highlight the importance of three general features – the 3’ SS region, intron length, and GC content (**Fig. 5E**; **Fig. S6C-E**); importantly, in the median model, relative to the earliest model, the 3’ SS features decrease in importance, whereas downstream, exonic features generally increase in importance (**Fig. S6C**), validating that median 3’ SS-RNAP II distances represent later splicing events.

Consistent with an early splicing timing for in-order NLI events, in both models the features flanking the 3’ SS (16 in total) dominated in feature importance (**Fig. S6D, E**). In the case of the median model, the 3’ SS-flanking features accounted for 51% of the predictive power (**Fig. 5E**; **Fig. S6E**). In particular, just as high U and low C content in the BP-3’ SS region predict high levels of in-order splicing and thus early splicing (**Fig. 4A**; **Fig. S4C**), high U and low C content in the same region predict shorter 3’ SS-RNAP II distances and thus earlier in-order splicing (**Fig. 5E**; **Fig. S6D-G**), suggesting that early in-order splicing is promoted by optimal binding of the U2AF heterodimer (Kang et al., 2020). To test this idea, we examined sequence motifs and U2AF1/U2AF2 binding in two classes of introns distinguished by their 3’ SS-RNAP II distances: “early in-order” introns (≥70% of 3’ SS-RNAP II distances ≤ 50 nts, n=1,576), and “late in-order” introns (≥70% of 3’ SS-RNAP II distances ≥ 200 nts, n=2,639). Indeed, the BP-3’ SS region of early in-order introns is enriched for U-rich hexamers and a strong U2AF2 consensus site, whereas the same region of late in-order introns is depleted of U-rich hexamers and is instead enriched for C/U-rich hexamers and a weak U2AF2 consensus site (**Fig. 5F, G**) (Shao et al., 2014). Although high U content and low C content in the region immediately downstream of the 3’ SS also predict short 3’ SS-RNAP II distances (**Fig. 5E**; **Fig. S6E**), this region in early in-order introns is enriched for distinct U-rich hexamers, which are depleted in late in-order introns, and also enriched for a modest U/A-rich motif (**Fig. S6H, I**), highlighting the possibility that factors beyond U2AF bind to the 3’ SS region to impact lariat formation timing. Significantly, by analyzing published eCLIP-seq data (Van Nostrand et al., 2020), we confirmed that early in-order introns, relative to late in-order introns, showed increased interactions between the 3’ end of an intron and the U2AF heterodimer (**Fig. 5H**), but not a control splicing factor (e.g., Aquarius; **Fig. 5H)**. These observations indicated that optimal U2AF binding promotes early lariat formation.

As with the order of splicing models, intron length emerged as an important feature in both 3’ SS-RNAP II distance models; indeed, intron length is the single most important feature in both models, accounting for ∼17% of the predictive power of the median model, for example (**Fig. 5E**; **Fig. S6E**). (Note: downstream intron length also emerged as a contributing feature but for technical reasons; see Methods.) Unexpectedly, whereas in the splicing order models shorter introns (<1 kb) predict higher levels of in-order splicing and thus earlier splicing overall (**Fig. 4A**), in the 3’ SS-RNAP II distance models shorter introns (<400 nts) predict longer 3’ SS-RNAP II distances and thus relatively later in-order splicing (**Fig. 5I**; **Fig. S6D, E**). Indeed, in the median model, for introns shorter than 300 nts, intron length outranks all other features (**Fig. S6J**); for the 325 introns that are all shorter than 200 nts, intron length accounts for over 50% of the predictive power of the model, and for the shortest introns, the median model predicts increases in 3’ SS-RNAP II distance approaching up to an additional 150 nts (**Fig. 5I**). Consistent with this analysis, shorter introns correlate with longer median 3’ SS-RNAP II distances (**Fig. S6K**). These observations indicate that short introns (<400 nts) initiate splicing when RNAP II is further downstream from the 3’ SS than longer introns (>400 nts) do, suggesting that splicing of these short introns (<400 nts) is delayed by a minimum elapsed time required for splicing after transcription of the 5’ SS, a time that may reflect constraints on the speed of spliceosome assembly (**Fig. 5J**; see Discussion).

The remaining top features correspond to the GC content of the NLI intron, exon, and downstream intron, which account, for example, for 19% of the predictive power of the median model (**Fig. 5E**; **Fig. S6E**). As with the order of splicing models (**Fig. 4A**), lower GC content in the exon predicts shorter 3’ SS-RNAP II distances and thus earlier in-order splicing (**Fig. 5E**; **Fig. S6E, L, M**), again suggesting that low secondary structure potential in this region promotes early splicing. However, in contrast to the order of splicing models (**Fig. 4A**), higher GC content in both NLI and downstream introns predicts shorter 3’ SS-RNAP II distances and thus earlier in-order splicing (**Fig. 5E**; **Fig. S6E, L**), which may reflect an outsized impact of GC content on transcription and thereby splicing timing when splicing is fast (see Discussion).

### Early co-transcriptional lariat formation implicates widespread splicing independent of exon definition

Because the exon definition pathway is thought to predominate in humans (Berget, 1995; Hollander et al., 2016), we expected that in the modeling of 3’ SS-RNAP II distance that the strength of the downstream 5’ SS and its position relative to the 3’ SS (i.e., exon length) would contribute to predicting 3’ SS-RNAP II distance. Indeed, a stronger downstream 5’ SS and a shorter exon predict earlier in-order splicing in the median model (**Fig. S6E**); however, these features together accounted only modestly (5%) for the predictive power of the median model (**Fig. S6E**), indicating that exon definition may only modestly contribute to the timing of in-order splicing when RNAP II is within hundreds of nts downstream of the 3’ SS. Additionally, exon length does not correlate with in-order splicing (**Fig. S5D**).

To test further for a role for exon definition, we asked whether in-order splicing timing depends on transcription of the downstream 5’ SS by examining the 3’ end locations of in-order NLI reads relative to the downstream 5’ SSs. We observed a broad distribution of 3’ end locations extending both downstream and upstream of 5’ SSs (**Fig. 6A)**; both classes of 3’ end locations were reduced by transcription or splicing inhibition (**Fig. S7A**), indicating that both require ongoing transcription and splicing. The same distribution can be observed for individual introns, such as at intron 10 of *MYG* (**Fig. 6B**). When considering only in-order splicing reads, some introns splice primarily upstream or downstream of the 5’ SS (**Fig. S7B**), but even when considering all orders of splicing, only 10% of assayed introns splice exclusively downstream of the downstream 5’ SS (**Fig. S7C**), although 50% of introns splice upstream of the 5’ splice site ≤20% of the time. Notably, compared to introns upstream of constitutive exons, introns upstream of cassette exons tend to splice when RNAP II is downstream of the downstream 5’ SS (**Fig. S7D**), consistent with evidence that cassette exons splice later (**Fig. 3K, L**; Drexler et al., 2020; Kim et al., 2017) and the expectation that cassette exons are more dependent on exon definition mechanisms (De Conti et al., 2013). Although CoLa-seq does not enable quantitation of the absolute fraction of splicing events that occur upstream of a downstream 5’ SS, these observations imply that the vast majority of introns can splice, at least to some extent, independent of a downstream 5’ SS and consequently independent of exon definition that relies on U1 snRNP binding to the downstream 5’ SS.

**Figure 6.**
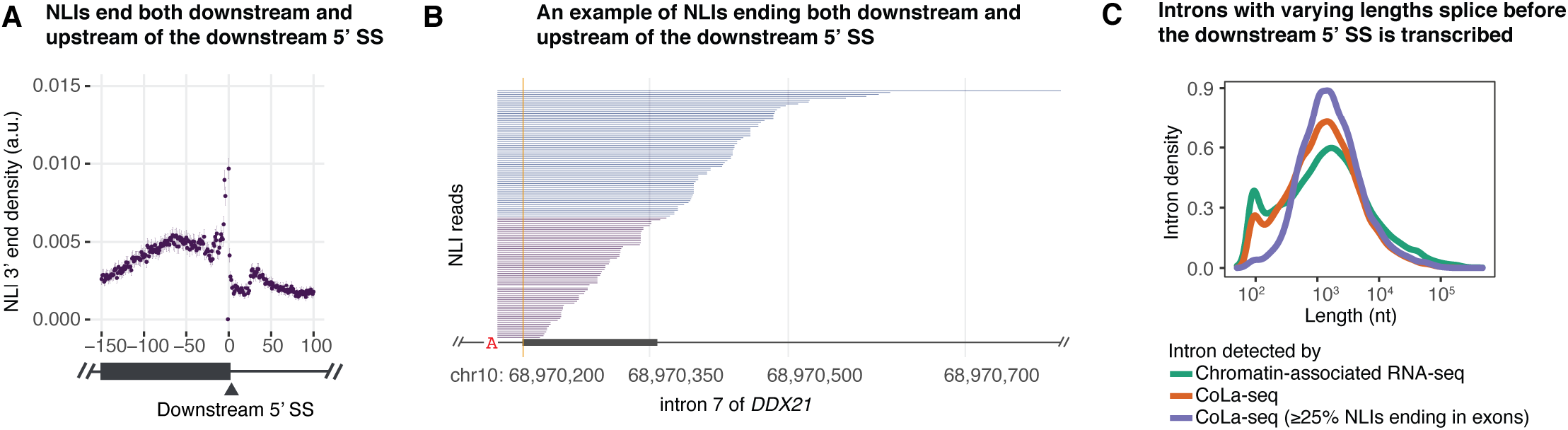
Early co-transcriptional lariat formation indicates widespread exon definition-independent splicing. **A.** The scatter plot illustrates the 3’ ends of NLIs around the downstream 5’ SS. The density of in-order NLI 3’ ends is plotted as a function of nucleotide position, relative to the 5’ SS; error bars reflect the 95% confidence interval. **B.** In-order NLI reads derived from intron 10 of *MYB* serve as an example for 3’ ends of NLIs that are located both downstream of the downstream 5’ SS (NLI tails colored blue) and upstream of the downstream 5’ SS (NLI tails colored purple). **C.** The density of introns, in particular early introns, is plotted as a function of intron length. Introns for which ≥25% of NLIs terminate upstream of the downstream 5’ SS are plotted in blue; for comparison, all introns detected by CoLa-seq (≥5 reads) are plotted in brown, and all introns detected by RNA-seq of chromatin-associated RNA are plotted in green.

Because intron definition is thought to be favored for short introns and exon definition is thought to be favored for longer introns (De Conti et al., 2013), we were surprised that our 3’ SS-RNAP II distance models predict that longer introns favored earlier in-order splicing (**Fig. 5I**). Consequently, we assessed the timing of splicing relative to the downstream 5’ SS as a function of intron length. Remarkably, introns spanning a wide range of lengths yielded at least 25% of NLI species with 3’ ends upstream of the downstream 5’ SS, including introns greater than 10 kb long (**Fig. 6C**; **Fig. S7E**). These observations indicate that longer introns do not necessitate U1-dependent exon definition, contrary to expectations based on the classic exon definition model and supporting observations (Berget, 1995; Hollander et al., 2016); early in-order splicing of long introns likely reflects the many additional features that contribute to the timing of co-transcriptional lariat formation (**Fig. 5D, E**; **Fig. S6D, E**).

### Evidence that early lariat formation driven by U2AF2 recognition of the PPT is AG/U2AF1-independent

Previous biochemical studies have found that some introns splice in an AG- and U2AF1-independent manner and that such introns generally have strong PPTs and interact more strongly with U2AF2 (Guth et al., 1999; Wu et al., 1999). Notably, we found that high U content in the BP-3’ SS region associates with and predicts shorter 3’ SS-RNAP II distance for NLIs and thus earlier splicing (**Figs. 5E, S6D-F, S8A**). Moreover, early in-order introns (≥70% of 3’ SS-RNAP II distances ≤50 nts) bind U2AF2 stronger than late in-order introns (≥70% of 3’ SS-RNAP II distances ≥200 nts; **Fig. 5F-H**). Therefore, we investigated whether strong recognition of the PPT by U2AF2 in early in-order introns renders lariat formation AG-independent and therefore U2AF1-independent.

To test for AG-independence, we asked whether any early in-order splicing events occurred before the AG has even emerged from the RNAP II exit channel. Indeed, for the majority of the early in-order class of introns, the 3’ end of the shortest 3’ SS-RNAP II distance is located less than 15 nts downstream of the 3’ SS (**Fig. 7A**), and thus the AG was still inside of RNAP II when splicing occurred, because RNAP II sequesters the last 15 nts of nascent RNA (Bernecky et al., 2017; Martinez-Rucobo et al., 2015). Although only a portion of the BP-3’ SS region of these ultra-early NLIs would be available for U2AF2 binding at the time of splicing, these emerged and exposed regions are long enough to accommodate U2AF2 binding (**Fig. S8B**) and have high U content, U-rich hexamers, and a strong U2AF2 binding motif (**Fig. 7B-D**); UUUUAG hexamers were de-enriched, ruling out upstream, cryptic AGs as the basis for ultra-early splicing (**Fig. 7C**). Together, these observations support the hypothesis that early lariat formation is dependent on strong U2AF2 binding and independent of the 3’ splice site AG and, by inference, U2AF1 binding (**Fig. 7E**).

**Figure 7.**
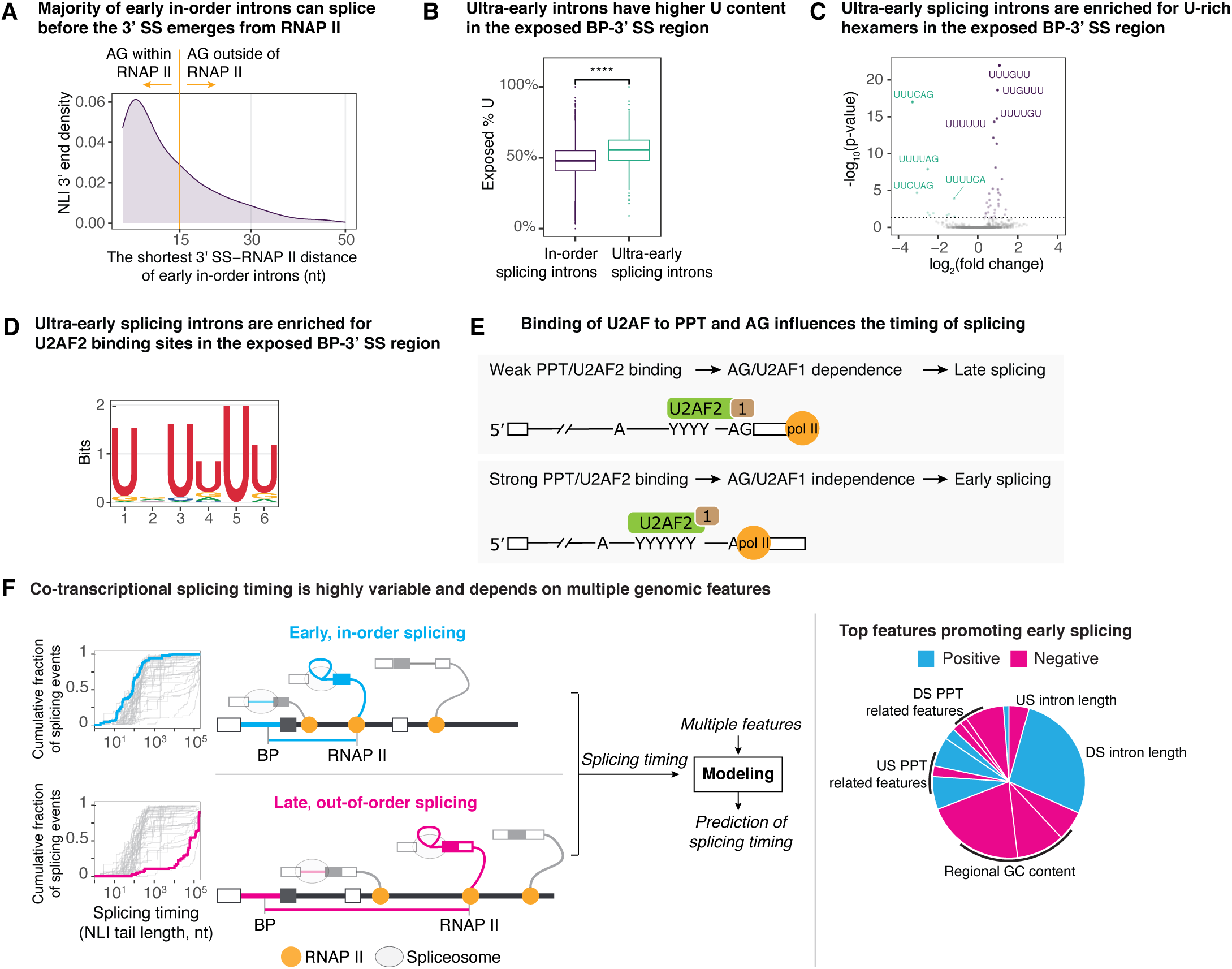
Early lariat formation is AG/U2AF1-independent. **A.** The density of NLI 3’ ends is plotted as a function of 3’ SS-RNAP II distance, in nucleotides, for the shortest 3’ SS-RNAP II distance of the early in-order introns. **B.** The box plot illustrates that the exposed BP-3’ SS region of ultra-early introns has higher U content than the BP-3’ SS regions of all in-order splicing introns. For ultra-early introns, the exposed region is defined as the region extruded from RNAP II. **C.** Hexamer analysis identified U-rich hexamers enriched in the exposed BP-3’ SS regions of ultra-early introns, relative to the BP-3’ SS regions of all in-order splicing introns, indicating that the exposed regions of ultra-early introns harbor strong U2AF2 binding sites. Significantly-enriched hexamers are in purple, and significantly-depleted hexamers are in green (Benjamini-Hochberg corrected p-value <0.05). P-values (-log_10_(p-value)) are plotted as a function of the hexamer fold enrichment (log_2_(fold change)). **D.** The U2AF2 binding sites were identified by motif similarity search after *de novo* analysis for motif enrichment in ultra-early introns, relative to all in-order splicing introns, indicating that U2AF2 binding sites are enriched in ultra-early splicing introns, relative to all in-order splicing introns. **E.** Model illustrating the interplay between splicing timing and AG/U2AF1-dependence. A weaker PPT necessitates U2AF1 binding to the 3’ splice site AG for efficient U2AF2 recruitment, thereby necessitating later splicing. By contrast, a strong PPT enables strong binding by U2AF2 and independence of U2AF1 binding to the 3’ SS AG, thereby enabling ultra-early splicing. “1” indicates U2AF1. **F.** The timing of co-transcriptional splicing is highly variable across introns; examples of early and late splicing are highlighted in blue and pink, respectively. The variable timings of co-transcriptional lariat formation were used to train computational models to identify features that contribute to the prediction of the model. A large number of genetic features, including key features, were found to contribute to splicing timing.

## Discussion

The spliceosome excises the majority of introns co-transcriptionally (Herzel et al., 2017), and the timing of splicing relative to transcription can impact splicing outcomes (Saldi et al., 2016), yet our understanding of co-transcriptional splicing timing has been limited. In this work, we have developed CoLa-seq to define the timing of lariat formation relative to transcription on a genome-wide scale (**Fig. 1**). We observed that the timing of lariat formation varies by five orders of magnitude across introns, that lariat formation within an intron can occur at different times, and that the splicing timing of regulated introns is later than for constitutive introns (**Fig. 3**). Further, machine learning-based modeling revealed that multiple features contribute combinatorially to predictions of the timing of lariat formation, with intron length, GC content, and the PPT playing major roles (**Figs. 4, 5**) and with strong binding of the PPT by U2AF2 predicting early, in-order splicing (**Figs. 4C, 5F-H, 7B-D**).

Unexpectedly, we observed widespread lariat formation before transcription of a downstream 5’ SS (**Fig. 6**) and cases of lariat formation even before the 3’ SS has emerged from RNAP II (**Fig. 7**), both implying that exon definition is not obligatory, even with long introns, and that instead multiple mechanisms initiate splicing in humans. By enriching for lariat RNA species, CoLa-seq also enabled annotation of BPs with unprecedented efficiency and yield (**Fig. 1**) and revealed coupling of BP and 3’ SS choice (**Fig. 2**). Overall, our analysis of splicing by CoLa-seq has revealed that the timing of lariat formation varies widely, can be fast, and depends particularly on several genomic features (**Fig. 7F**), all of which have important implications for the regulation of splicing.

The validity of splicing timing as defined by CoLa-seq is favored and supported by our procedures and observations. Although splicing in principle might have the potential to occur during sample preparation, thereby biasing splicing to shorter times, we included EDTA at the start of cell lysis to quench splicing, given that EDTA efficiently inhibits splicing catalysis (e.g., Fica et al., 2013; Sontheimer et al., 1997). Indeed, we found that splicing inhibition during cell growth alone drastically reduced the levels of lariat RNA reads (**Fig. S1B**), ruling out pervasive *de novo* spliceosome assembly *in vitro*. Further, the impact of NLI intron length and downstream intron length on splicing timing (**Figs. 4D, 4E, 5I**) directly implicates co-transcriptional splicing, consistent with the dependence of lariat RNA reads on transcription **(Fig. S1B**). Although NLIs could in principle reflect stalled intermediates, the vast majority of NLIs correlated with cognate ELIs (**Fig. S2L, M**), consistent with the conversion of NLIs to ELIs, and introns yielding NLIs did not differ substantially from those introns that did not (**Fig. S2N**), and introns that did not yield NLIs were distinguished simply by poor expression in K562 cells (**Fig. S2H**). Our estimates of splicing timing can be impacted not only by differing rates of NLI formation but also by differing rates of exon ligation (**Fig. S5A**). In principle, our estimates might also be impacted by other factors that could influence the levels of NLIs, such as differing rates of spliceosome disassembly or RNA decay, but these processes cannot easily account for many of the features predictive of splicing timing (e.g., PPT). An impact of decay on our estimates is also unlikely given that we observed coherent 3’ ends of CoLa-seq reads at 3’ splice sites (**Fig. 1C**), representing ELIs, and at 5’ splice sites (**Fig. S3G),** representing concurrent splicing intermediates. Furthermore, CoLa-seq yielded observations of splicing timing that align with previous studies; for example, CoLa-seq indicated that splicing is delayed in (i) introns flanking cassette exons, relative to constitutive introns (**Fig. 3K, L**), and (ii) U12 introns, relative to U2 introns (**Fig. S3N**; cf. Herzel and Neugebauer, 2015; Pai et al., 2017; Pandya-Jones and Black, 2009; Wachutka et al., 2019).

Though an essential reactant in the splicing reaction, the BP is insufficiently annotated in humans, largely due to the lack of an efficient method for BP mapping. Using CoLa-seq, we mapped the largest BP dataset to-date (165,282 BPs), with the majority of BPs novel (**Fig. 1E**; **Fig. S2D, G**). We demonstrate that CoLa-seq is an efficient method for branch point mapping on a genome-wide scale, paving the way for future studies on the regulation and perturbation of BP usage, such as the impact of somatic cancer mutations on BP and 3’ SS usage (Darman et al., 2015) (cf. **Fig. 2D-F**).

By focusing on un-spliced and spliced nascent transcripts and using long-read sequencing, recent genome-wide studies have been limited by depth and consequently have resorted to analyzing global splicing timing by aggregating reads across different introns. Further, estimates of splicing timing that rely on linear transcripts can be complicated by the contamination of chromatin-associated RNA with abundant mature mRNA; indeed, previous reports of splicing timing differ significantly. In this work, we took advantage of NLIs, because they (i) cannot derive from mature mRNAs, (ii) do allow for enzymatic enrichment, given their unique lariat structure, (iii) indicate splicing is in progress, (iv) reveal concurrent splicing, (v) offer the potential to independently report on the kinetics of the two chemical steps of splicing, and (vi) uncouple lariat formation, which reflects most regulation, from exon ligation. Importantly, using short-read sequencing, we were also able to examine the timing of co-transcriptional lariat formation on an individual intron level. We found that while co-transcriptional splicing occurs generally before an average-sized downstream intron (∼1.5 kb) is fully transcribed, splicing timing varies over a broad range of times across introns (**Fig. 3F**) and the range of splicing timing varies for individual introns (**Fig. 3G**). At one extreme, ultra-early lariat formation appeared to occur when RNAP II was only 3 nts downstream of the 3’ SS, indicating that splicing is surprisingly fast or that RNAP II is paused near the 3’ SS (e.g., at nucleosomes positioned at exons; Tilgner et al., 2009). At the other extreme, the latest splicing events detected by CoLa-seq occurred when RNAP II was more than a million nts downstream of a 3’ SS, implying splicing times of more than 10 hours. The wide range of timings accessible to splicing provides an opportunity for splicing regulation. For instance, changes in splicing timing could alter the levels of out-of-order splicing, which we observe widely, and thus change the availability of RNA regulatory elements (e.g., Nasim et al., 1990; Takahara et al., 2002).

Our work raises important questions concerning the mechanisms of splice site recognition in humans. Consistent with evidence for exon definition in humans (Berget, 1995; Drexler et al., 2020; Hollander et al., 2016; Sousa-Luís et al., 2021), many introns undergo lariat formation after transcription of a downstream 5’ SS (**Fig. 6A**). However, many introns also undergo lariat formation before transcription of a downstream 5’ SS (**Fig. 6A**), inconsistent with U1-dependent exon definition. Although future studies are required to determine what fraction of splicing is exon-definition independent, our observations are consistent with recent genomic studies also reporting splicing before transcription of a downstream 5’ SS in mammalian cells (Reimer et al., 2021; Sousa-Luís et al., 2021). Strikingly, we observed lariat formation in the exon not only for short introns but also for long introns (**Fig. 6C**). This result is unexpected because inefficient splice site recognition across a long intron has been thought to necessitate exon definition (Berget, 1995; Hollander et al., 2016). However, while an important feature, intron length is just one of multiple predictive features in our models (**Fig. 5D, E**; **Fig. S6D, E**). We envision several mechanisms by which long introns could splice early. First, in a variation of exon definition, factors that bind to exonic splicing enhancers downstream of the intron could aid 3’ SS recognition in lieu of U1 snRNP binding to the downstream 5’ SS (Wu and Maniatis, 1993; Ule and Blencowe, 2019). Second, upstream exon tethering mediated by U1 snRNP bound to transcribing RNAP II could effectively mitigate any penalty for long introns and obviate the need for U1 snRNP bound to the downstream 5’ SS (Das et al., 2007; Leader et al., 2021; Nojima et al., 2018; Zhang et al., 2021). Notably, though, the contribution of intron length to predicting splicing timing implies that exon tethering is insufficient to overcome entirely a penalty for longer introns (**Fig. 4D**). Lastly, because these early lariat formation events are characterized by a strong PPT and strong U2AF binding (**Fig. 5F-H**), and binding of U2AF2 to a strong PPT can induce an active conformation of U2AF2 independent of U2AF1 (Warnasooriya et al., 2020), we suggest that at some introns U2AF2 binds to a PPT in a manner also independent of either a downstream or upstream 5’ SS – that is, by neither canonical exon nor intron definition. Indeed, U2AF2 along with co-factors can bind in extract to a substrate that includes a BP, PPT, and 3’ splice site AG but lacks any 5’ SS (e.g., Mackereth et al., 2011; Maul-Newby et al., 2021), and the association of U2AF2 with RNAP II (David et al., 2011) could allow rapid and efficient recognition of the PPT. Reconciling evidence for early splicing and exon definition will be an important goal for the field in the future.

The depth of our CoLa-seq dataset enabled modeling of splicing timing for individual introns (**Figs. 4, 5**). Importantly, we found that a large number of genetic features contribute combinatorially to predictions of splicing timing models (e.g., **Fig. 4B, 5D**). Nevertheless, we uncovered three key features – U2AF binding, intron length, and local GC content. In both the splicing order and 3’ SS-RNAP II distance models, we found that a strong PPT predicts early splicing and that U2AF binds strongly to introns that splice early (**Figs. 4, 5**). This result suggests an explanation for the positive correlation between U2AF binding and exon inclusion (Shao et al., 2014) – that early splicing, mediated by U2AF, favors in-order splicing and thus exon inclusion. Strikingly, in introns with strong PPTs, we detected ultra-early lariat formation events that occurred before the 3’ SS has emerged from RNAP II (**Fig. 7A**), consistent with previous biochemical studies indicating that PPT strength correlates with the efficiency of spliceosome assembly (Mackereth et al., 2011) and that introns with strong PPTs splice independently of the 3’ splice site AG and U2AF1 (Guth et al., 1999; Wu et al., 1999). Ultra-early splicing introns tend to have longer BP-3’ SS distances that provide enough exposed region for U2AF2 binding (**Figs. 7B-D, S8B**); indeed, long BP-3’ SS distances correlate with strong U2AF2 binding near the BP (Briese et al., 2019). Together, these findings imply that U2AF2 can drive fast splicing and dictate splicing outcomes, rationalizing why this factor serves as a key target for regulatory factors.

Consistent with fundamental polymer dynamics (Toan et al., 2006) and previous biochemical findings that shorter introns facilitate splice site recognition (e.g., Fox-Walsh et al., 2005), we found that shorter upstream introns predict higher levels of in-order splicing (**Fig. 4D**). Notably, this observation is the opposite of what we observe from a simple correlation analysis (**Fig S4A**), but such correlations fail to regress out the effects of associated features, such as GC content, which anti-correlates with intron length, highlighting the value of our modeling. Although shorter introns generally predict higher levels of in-order splicing, relative to concurrent and out-of-order splicing, a finer analysis of the splicing timing of in-order splicing in terms of 3’ SS-RNAP II distance revealed unexpectedly that for short introns (≤400 nts), shorter intron lengths predict increasingly later timings of the earliest lariat formation (**Fig. 5I**), suggesting that a minimum time of lariat formation is longer than the transcription time of these short introns (≤400 nts; **Fig. 5J**). This delay is unlikely to reflect a minimum time for recognition of the 3’ end of the intron, because 3’ SS recognition can occur quite early (**Figs. 5B, 5C, 7A**). Instead, the delay more likely reflects a minimum time required for 5’ SS recognition by U1 snRNP or a subsequent step in spliceosome assembly.

Longer downstream introns predict higher levels of in-order splicing (**Fig. 4E**), implying that a longer time for transcribing the downstream intron delays splicing of the downstream intron, relative to the upstream intron. Notably, our simulations (**Fig. 3J, S3K-M**) indicate that an average intron splices on a timescale that is shorter than the timescale for transcribing an average, downstream intron. In the context of cassette exons, in-order splicing occurs on a timescale that precludes the possibility of exon skipping, whereas concurrent splicing and out-of-order splicing, requiring transcription of the downstream intron, occur on a timescale that permits exon inclusion or exon skipping. Thus, the transcription time of the downstream intron represents a compelling target for regulating exon skipping, consistent with the general idea that transcription serves as a key target for splicing regulation (Saldi et al., 2016).

High GC content in the NLI intron, the exon, and the downstream intron predicts low levels of in-order splicing (**Fig. 4A, S4C, S5F**). We also found that introns within the same genes share more similar splicing profiles than introns from different genes, consistent with previous findings that GC content bias extends well beyond an intron and can encompass an entire gene (**Fig. S5G**; Amit et al., 2012; Costantini and Musto, 2017; Costantini et al., 2006). Thus, the correlation of splicing timing of nearby introns, also observed by others (Drexler et al., 2020; Reimer et al., 2021), may not reflect coordination of splicing between adjacent introns but rather reflect the impact of regional GC content bias on splicing timing across multiple introns within a gene. Indeed, we discovered that U2AF binding, favored in GC-poor introns, correlates between adjacent introns (**Fig. 4C**). High GC content likely impacts splicing timing in multiple ways, such as weakening splice site strength (Amit et al., 2012), stabilizing local RNA secondary structure (**Fig. S5H**), accelerating or decelerating transcription (Saldi et al., 2021; Turowski et al., 2020; Veloso et al., 2014; Zamft et al., 2012), and associating with different regions of the nucleus, given the impact of GC content on the conformation of the genome (Lemaire et al., 2019; van Steensel and Belmont, 2017). Consistent with an impact on RNA structure and splice site recognition, GC-rich cassette exons are skipped after knockdown of the RNA helicases DDX5 and DDX17, which promote U1 snRNP binding (Lemaire et al., 2019). Further, given that pulse-chase experiments have indicated that the net effect of high GC content is to slow transcription (Jonkers et al., 2014; Veloso et al., 2014), high GC content might lead to short 3’ SS-RNAP II distance for early, in-order splicing, where the length (**Fig. 5I**) and likely transcription time of the NLI intron matters. Together, although future studies will be required to discern the primary impacts of GC content, our findings imply that the larger genomic context of an intron is important in modulating splicing timing.

## Acknowledgement

We thank D. Bishop and P. Connell laboratories for their help with human tissue culture, the Genomics Facility at University of Chicago for Illumina sequencing, and the Research Computing Center at the University of Chicago for providing computing resources. We thank the members of the J.P.S., Y.I.L., and A.J.R. laboratories for their helpful discussions, especially K. Nielsen. We thank J. Valcárcel, D. Bentley, and J. Pritchard for their comments on the manuscript. A.K. was supported by a postdoctoral fellowship from the American Heart Association. J.M.H. is supported by a predoctoral fellowship from American Heart Association (20PRE35180107). This work was funded by grants from the NIH (R01HL148719 to A.J.R., R01GM130738 to Y.I.L., R01GM062264 to J.P.S., and R01HG011067 to J.P.S. and Y.I.L.).

## Author contributions

Conceptualization, Y.Z. and J.P.S.; methodology, Y.Z., B.J.F., A.K., J.M.H., Y.I.L., and J.P.S.; experiments, Y.Z., A.K., Y.H.; data analysis, statistics, interpretation, and visualization, Y.Z., H.Z., B.J.F., Y.I.L., and J.P.S.; funding acquisition, Y.I.L., A.J.R., and J.P.S.; writing – original draft, Y.Z. and J.P.S.; writing – review and editing, Y.Z., B.J.F., Y.I.L., and J.P.S. with input from all authors.

## Methods

### Chromatin-associated RNA isolation

Chromatin RNA was isolated from K562 cells similar to previously described methods (Pandya-Jones and Black, 2009; Wuarin and Schibler, 1994; Werner and Ruthenburg, 2015), with modifications noted below. All the steps were performed with pre-chilled, ice-cold, freshly prepared RNase-free buffers using RNase-free equipment on ice or at 4 °C. Roughly 5% of subcellular fractions were saved for quality control (western and RNA fragment size analysis). Fifty million K562 cells were spun down at 100 x g for 3 minutes and washed twice with 15 mL of ice-cold 1X PBS (phosphate-buffered saline) and once with 1 mL of ice-cold 1X PBS. Washed cells were then gently resuspended in 1mL of 0.1% (v/v) Triton X-100-containing buffer A (10 mM HEPES (pH 7.5), 10 mM KCl, 10% glycerol, 0.116 g/mL sucrose, 1 mM DTT, 1x protease inhibitor, 2 mM EDTA) and lysed on ice for 9 min. After incubation, the lysed cells were centrifuged at 800 x g for 5 minutes at 4 °C. The supernatant corresponding to the cytoplasmic fraction was carefully removed. The remaining nuclei pellet was gently resuspended in 250 µL of NRB buffer (20 mM HEPES (pH 7.5), 75 mM NaCl, 50% glycerol, 0.5 mM EDTA, 1 mM DTT, 1x protease inhibitors). Then, 250 µL of NUN buffer (50 mM HEPES (pH 7.5), 300 mM NaCl, 0.2 mM EDTA, 1 M urea, 1 mM DTT, 1% NP-40) was slowly added and mixed by inversion. The resulting nuclei suspension was incubated on ice for 5 minutes and centrifuged at 800 x g for 5 minutes at 4 °C. The supernatant corresponding to the nucleoplasmic fraction was carefully removed. The chromatin pellet was washed once with 500 µL of premixed NRB/NUN buffer (1:1) and centrifuged at 800 x g for 5 minutes at 4 °C. The supernatant was carefully removed. The washed chromatin pellet was gently resuspended in 200 µL of NRB buffer. RNA from whole cells, the cytoplasmic fraction, the nucleoplasmic fraction, and the chromatin pellet were extracted by Trizol (Invitrogen) followed by a second extraction with phenol:chloroform:isoamyl alcohol (25:24:1) and isopropanol precipitation. The isolated RNA was furthered treated with Turbo DNase (Invitrogen) to remove residual genomic DNA, followed by extraction with phenol:chloroform:isoamyl alcohol (25:24:1).

### Confirmation of fractionation

The efficiency of fractionation was tested by western blotting of the cytosolic fraction, the nucleoplasmic fraction, and the chromatin fraction with appropriate antibodies. The antibodies used were GAPDH (Santa Cruz Biotechnology), RNA Pol II (Ser-5 phosphorylated, MBL International), and Histone H3 (Abcam).

### Enrichment for lariat RNA species

For each CoLa-seq library, chromatin-associated RNA from 200 million cells was used. To enrich for lariat RNA species (nascent lariat intermediates and excised lariat introns) from the chromatin-associated RNA, ≤200 nts RNAs were removed using SPRI paramagnetic beads (Beckman Coulter) with a beads to sample ratio of 1:1.2. Next, ribosomal RNA was depleted using oligo-based Ribo-Zero Magnetic Gold Kit (Illumina) according to the manufacturer’s manual. During the optimization of CoLa-seq, we noticed abundant ncRNA reads in our libraries, similar to previous observations in mNET-seq (Mayer et al., 2015). To overcome this issue, biotinylated anti-sense oligos were designed to specifically target and deplete ncRNAs that were found to be abundant in our initial libraries (**Table S4**). Briefly, for every μg of rRNA-depleted RNA, a depletion reaction was assembled, containing 82.5 pmol of total depletion oligo mix (**Table S4**), 2X SSC, and 5 mM EDTA. The depletion reaction was put through an oligo annealing cycle (5 min at 75 °C, cool down to 22°C at 0.1 °C/sec), after which ncRNA bound biotinylated depletion oligos were pulled down using Dynabeads™ MyOne™ Streptavidin C1 beads (Invitrogen), according to the manufacturer’s manual. The RNA in the supernatant was purified using the RNA Clean and Concentrate kit (Zymo Research) and excess anti-sense oligos were removed by treatment with Turbo DNase (Invitrogen). Finally, according to the manufacturer’s instructions, purified RNA was treated with RppH (NEB) to decap linear RNA and then was degraded using Terminator 5’-3’ exonuclease (Lucigen). The resulting RNA was put through a size selection step to remove ≤200 nts RNA using SPRI paramagnetic beads (Beckman Coulter) with a beads to sample ratio of 1:1.2. The final RNA quantity is about 0.5-1% of the starting material.

### CoLa-seq library prep

Once the chromatin RNA was enriched for lariat species, ∼500 ng of RNA was used for CoLa-seq library prep. The 3’ ends of RNA were ligated to a pre-adenylated 3’ end adapter (**Table S5**), which includes a 6 nt UMI (unique molecule identifier) at the 5’ end to allow correction for PCR duplication and RT mis-priming. The ligation reaction contained 17.5% PEG-8000 (w/v), 1X T4 RNA ligase buffer (NEB), 1 µM of the pre-adenylated 3’ end adaptor, and 10 units/µL T4 RNA Ligase 2 (truncated K227Q, NEB). The ligation reaction was incubated for 3 hours at 22 °C. After the reaction excess adapters were removed by size selection using SPRI paramagnetic beads (Beckman Coulter) with a beads to sample ratio of 1.6:1. Then, the adapter-ligated RNA was used for cDNA synthesis using 0.25 µM of RT primer (**Table S5**) and 10 units/µL of Superscript III (Invitrogen) at 50 °C for 1 hour, according to the manufacturer’s manual. Afterwards, to degrade excessive RT primer, 3.5 µL of ExoSAP-IT was added to the cDNA synthesis reaction and incubated for another 30 min at 37 °C. The reaction was stopped by adding 10 µL of 0.5M EDTA and RNA was degraded by adding 10 µL of 1 N NaOH. The resulting cDNA was cleaned up using Oligo Clean & Concentrator columns (Zymo Research) and was circularized with Circligase (Lucigen), according to the manufacturer’s manual. Final library amplification was performed on the circularized cDNA using Phusion polymerase (Thermo Fisher Scientific) with primers that add sequencing adaptor sequences and specific sequencing indices, which are located in the reverse primers (**Table S5**). The number of PCR cycles was between 15 and 18 cycles. The amplified product was run on a 6% TBE gel and DNA fragments between 175 and 750 bps were gel purified and 3 nM of the library was sequenced in a 100 bp paired-end mode on an Illumina HiSeq 4000 (Genomics Facility at University of Chicago). Reads were preprocessed and analyzed as described below.

### Sequencing of chromatin-associated RNA

For each library, 10 μg chromatin-associated RNAs were depleted of ribosomal RNA depletion using Ribo-Zero Magnetic Gold Kit (human/mouse/rat; Illumina), according to the manufacturer’s instruction. Abundant ncRNAs were depleted as described in the section of enrichment for lariat RNA species (see above). Then, 100 ng rRNA/ncRNA-depleted RNA was used to prepare an RNA-seq library using NEBNext Ultra™ Directional RNA Library Prep Kit for Illumina, according to the manufacturer’s instruction. The final library was sequenced in a 100 bp paired-end mode on an Illumina HiSeq 4000 (Genomics Facility at University of Chicago). Nascent RNA-seq data were pre-processed as follows: (1) adaptors were trimmed using Cutadapt (Martin, 2011), with default parameters except discarding reads shorter than 15 nts, (2) remaining reads were aligned to the human reference genome (hg38) with GENCODE transcriptome annotation (v26) using the STAR software package (Dobin et al., 2013) with the following settings: --outSAMattributes NH HI AS nM NM MD jM jI XS, --outFilterMultimapNmax 50, --outFilterMismatchNmax 15, -- outSJfilterDistToOtherSJmin 10 0 5 10 --alignIntronMin 15, --outSAMtype BAM SortedByCoordinate, --twopassMode Basic, (3) aligned reads were further filtered for unique mapping and proper alignment using samtools (Li et al., 2009). The resulting reads were used for downstream analyses. Separately, after the adapter removal step, reads were used for gene expression quantitation using Kallisto with default settings (Bray et al., 2016).

### Lariat sequencing

Lariat-seq was used to identify BPs via inverted reads that traverse through the branch in the lariat RNAs and performed similarly to previously described methods (Mayerle et al., 2017; Mercer et al., 2015) with modifications. Briefly, chromatin-associated RNA was first isolated as above to capture both lariat intermediates and excised lariat introns. Then, 10 μg chromatin-associated RNAs were depleted of ribosomal RNA depletion using Ribo-Zero Magnetic Gold Kit (human/mouse/rat; Illumina), according to the manufacturer’s instruction. To further enrich lariat RNA species, RNase R (Lucigen) digestion was performed to degrade linear RNA, according to the manufacturer’s instruction; treatment also trims tails of lariat RNA species. The resulting RNA was used to prepare an RNA-seq library using NEBNext Ultra™ Directional RNA Library Prep Kit from Illumina, according to the manufacturer’s instruction. The final library was sequenced in a 100 bp paired-end mode on an Illumina Hiseq 4000 (Genomics Facility at University of Chicago). The resulting reads were used to identify branch points according to previously described methods (Mercer et al., 2015; Pineda and Bradley, 2018).

### Splicing inhibition and transcription inhibition

K562 cells at 90% confluency were treated for 2 hours with one of the following reagents (1) pladienolide B (Santa Cruz Biotechnology) in DMSO to a final concentration of 10 μM or (2) flavopiridol (Sigma) in DMSO to a final concentration of 1 μM. A control treatment was also performed with an equal volume of DMSO. Splicing inhibition was confirmed by RT-PCR to detect previously observed splicing changes (Nojima et al., 2015); primers used are listed in **Table S6**. Chromatin fractionation and CoLa-Seq libraries were prepared from the DMSO or drug treated cells as described above.

### Data pre-processing of CoLa-seq data

CoLa-seq data were processed as follows: (1) 6 nt UMIs were trimmed from reads and attached to read names to maintain their association with reads using fastp (Chen et al., 2018a), (2) adaptors were trimmed using Cutadapt. Reads shorter than 15 nts were discarded, (3) the remaining reads were aligned to the human reference genome (hg38) with GENCODE transcriptome annotation (v26) using the STAR software (Dobin et al., 2013) with the following settings: --outSAMattributes NH HI AS nM NM MD jM jI XS --outFilterMultimapNmax 50 --outFilterMismatchNmax 15 -- outSJfilterDistToOtherSJmin 10 0 5 10 --alignIntronMin 15 --outFilterScoreMinOverLread 0 -- outFilterMatchNminOverLread 0 --outFilterMatchNmin 15 --outSAMtype BAM SortedByCoordinate -- twopassMode Basic, (4) aligned read pairs were further filtered for unique mapping using samtools, (5) to remove RT mis-primed reads, raw reads were separately mapped without UMI trimming. If the UMI portion of a read was mapped to the genome without any mismatch, it is considered as a mis-primed read and removed from the reads that were mapped with UMI trimming. These RT mis-primed reads were removed using samtools and pysam, (6) PCR duplicates were removed using UMI-tools (Smith et al., 2017), (7) reads overlapping with ncRNAs (snRNAs, snoRNAs, miRNAs, rRNAs) were removed using samtools based on the GENCODE reference (v26). The remaining reads were used for downstream analyses.

### Analysis of CoLa-seq read ends around branch points, 5’ splice sites, or 3’ splice sites

The 5’ ends of reads were intersected with 21-nt regions centered on previously annotated BPs (Pineda and Bradley, 2018), whereas read 3’ ends were intersected with 21-nt regions centered on either constitutive 5’ SSs or constitutive 3’ SSs (see below). To minimize expression differences, reads numbers across each region for each intron were normalized such that the total read count ranged from 0 to 1. Then, regions were aligned with their respective feature (BP, 5’ SS, or 3’ SS) in the center, and the mean, normalized signal was calculated at each position.

### Sequence logo of branch point motif

Using Bioconductor tools and ggseqlogo package in R (Wagih, 2017; Huber et al., 2015), the sequence logo was generated as follows: (1) for each branch point, an 11 nt long sequence (with the branch point in the middle) was extracted, (2) the resulting sequence set was used to calculate the position weight matrix and then to plot the sequence logo for the branch point motif.

### Branch point calling algorithm

Putative ELI and NLI fragments were identified as sequenced fragments for which their 5’ ends mapped to within 100 nts of an annotated 3’SS, and putative ELI fragments were further defined as sequenced fragments for which their 3’ end maps to an annotated 3’ splice site (Gencode v29), and putative NLI fragments were further defined as extending without gaps past an annotated 3’ splice site. To finally filter for authentic ELI and NLI fragments, we aimed to require that their 5’ ends align with a BP. Because published databases of known branch points (Pineda and Bradley, 2018) are relatively incomplete, we built a probabilistic classifier to expand the set of branch point positions, from which NLI reads are anchored. Specifically, each position in the 5 to 100 nt region upstream of annotated 3’ splice sites is assigned a probability that it is a branch point based on multiple features and a naive Bayes assumption of independence. The probability distribution of features over the non-branch points is assumed uniform and therefore a constant that can be ignored at all positions.

Therefore, at each position, *X*, we can assign an unscaled posterior probability that the position is a BP:

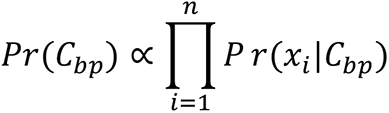

where *Pr*(*C_bp_*) is the posterior probability of classification as a branch point C*_bp_* (as opposed to non-branch point *C_n_*) at position **X**, and *x_i_* is the value of feature *i* at position *X*. The features considered are the pentamer motif (−3 to +1 relative to position *X*), the distance to the annotated 3’ SS, and the relative coverage of CoLa-seq 5’ read ends. The probability distribution of pentamers in *C_bp_* was trained on branch points empirically identified previously (Pineda and Bradley, 2018). Specifically, all U2-recognized, high-quality BPs were used to generate a position weight matrix (weighted by the read count supporting each branch point in Pineda and Bradley, 2018) to assign branch point probabilities for each pentamer. Similarly, the empirical distance from a U2-recognized, high-quality BP to the annotated 3’ SS (across all BPs) was used to create a smoothed probability distribution using the density function in R with bw=3 bandwidth adjustment. Feature training and classification were repeated separately for introns annotated as U12 introns (Olthof et al., 2019). To determine a branch point probability from relative coverage of CoLa-seq 5’ ends (P*r*(*x_i_*|*C_bp_*)) where *x*_*i*_ is the relative coverage of CoLa-seq 5’ ends at known branch points, *C_bp_*, we did not consider the empirical probability of relative coverage from known branch points, but, rather, we devised an alternate probability function as follows: CoLa-seq reads from all experiments were combined and filtered for fragments that end at (putative ELI) or cross (putative NLI) an annotated 3’ SS. For each 3’ SS region with at least 10 ELI and/or putative NLI fragments, read coverage of the 5’ position of fragments was normalized to the total coverage in the region after adding a 0.1 pseudo-count to all positions. Then, a smoothed probability distribution was created with the density function in R with a bandwidth setting of bw=0.5. Smoothing was introduced as a way to emulate the natural imprecision of reverse transcription termination, as we have observed that though reverse transcriptase usually stops one nucleotide downstream of the BP, reverse transcriptase sometimes terminates one nucleotide further downstream or one nucleotide upstream (i.e., at the branch point; **Fig. S2A**). Following the evidence that putative ELI and putative NLI CoLa-seq reads most often terminate at the base downstream of the branch point (**Fig. S2A**), the probability distribution was shifted upstream by one base such that the center of smoothed peaks corresponds to more likely branch points. Finally, to classify bases as belonging to a branch point region, we defined a threshold

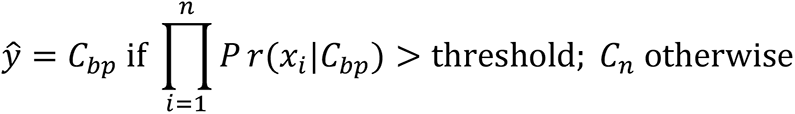

Or equivalently:

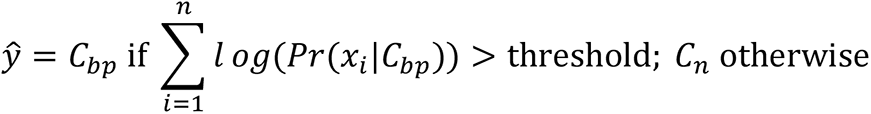

The threshold was chosen based on ROC analysis using a validation set of branch point positions identified in both Pineda and Bradley, 2018, as well as lariat-seq from K562 (this study; see above; **Table S1**), as the true response. We chose a threshold that yields a false positive rate of 0.01 in the ROC analysis. Finally, after applying this classifier to all 114,540 3’ SS regions with >10 ELI/LI reads, we merged branch points with one or zero bases in between each other into a single most likely branch point. In total, we identified 164,532 BPs amongst 109,256 U2-type 3’ SSs and 750 BPs amongst 583 U12-type 3’ SSs.

### Identification of nascent lariat intermediates and excised lariat introns

Exons from the GENCODE transcriptome annotation (v26) that we also identified in chromatin-associated RNA-seq and total RNA-seq were used for identifying NLIs and ELIs. Exons whose ends can be used as alternative 3’ splice sites were removed to avoid ambiguity. To identify reads derived from NLIs, the following filtering steps were performed using a custom snakemake (Köster and Rahmann, 2018) pipeline wrapped around bedtools (Quinlan and Hall, 2010), samtools, and Picard tools (2019): (1) reads were defined as putative NLI reads, as noted above, if they overlapped with an annotated 3’ SS and their 5’ ends were within 100 nts upstream of 3’ splice sites; and (2) using branch points annotated above, putative NLI reads were designated as genuine NLI reads if their 5’ ends corresponded to the BP or BP+1 position. To identify reads derived from ELIs, the following filtering steps were performed using a custom snakemake pipeline wrapped around bedtools, samtools, and Picard tools: (1) reads were defined as putative ELI reads, as noted above, if their 3’ ends corresponded to the last nt of an intron, and their 5’ ends were within 100 nts upstream of 3’ splice sites; and (2) using branch points annotated above, putative ELI reads were designated as genuine ELI reads if their 5’ ends corresponded to the BP or BP+1 position.

### Identification of constitutive and alternative splice sites

Using Regtools (Feng et al., 2018) and LeafCutter (Li et al., 2018), 5’ splice sites and 3’ splice sites were extracted from chromatin-associated RNA-seq data (this study), nuclear RNA-seq data from ENCODE, as well as total RNA-seq data from ENCODE. Splice sites were considered as constitutive sites when their usage percentages were higher than 95%.

### Quantification of branch point usage

Branch point usage was quantified as described in (Mercer et al., 2015). Briefly, ELI read (or NLI read) counts of a branch point were divided by ELI read (or NLI read) counts of all branch points associated with a 3’ splice site.

### Quantification of usage of coupled branch point and 3’ splice sites

Alternative 3’ splice sites using the same 5’ splice sites were paired together, and then the nucleotide distances between the alternative 3’ splice sites were calculated and used to sort these pairs into different groups. If both 3’ splice sites within a pair use the same set of branch points, the coupling index of this pair was set to 1, whereas if they used different ones, the coupling index was set to 0. The coupling fraction of each group was then calculated by taking the average of the coupling index for all 3’ splice site pairs in the group. NAGNAG sites were extracted from GENCODE transcriptome annotation (v26) and supplemented with NAGNAG sites from Bradley et al., 2012.

### Classification of in-order, concurrent, and out-of-order splicing

To further classify NLI reads into classes of in-order, concurrent, and out-of-order splicing, NLI reads that ended at the last nt of an annotated exon were classified as concurrent splicing. NLI reads that were split reads (N in the CIGAR string) were considered as out-of-order splicing. NLI reads that did not belong to the other two classes and were shorter than 650 nts (upper insert size limit of the library) were considered as in-order splicing; note that for intron pairs in which the downstream intron is short, <600 nts, the likelihood of concurrent splicing and out-of-order splicing increases, and consequently the likelihood of capturing in-order NLIs with longer tails decreases.

### Splicing timing simulation

The level of in-order splicing of an intron relative to its downstream intron strongly correlates with the distance from its 3’ SS to the 3’ SS of the downstream intron (US-DS 3’ SS distance; **Fig. 3I**). Importantly, the median level of in-order splicing reaches over 75% when the US-DS 3’ SS distance exceeds 2000 nts. We thus reasoned that the timing of splicing must be generally faster than the transcription time of a 2000-nt intron. To derive an estimate of the median timing of splicing for all introns, we set out to simulate the median proportion of in-order splicing for adjacent introns as a function of the US-DS 3’ SS distance (***ds***, in nts). To this end, we sampled splicing timing for an intron (*1*) and its downstream intron (*2*) from distributions with a range of possible median splicing timings, ranging from 0 to 5000 nt. More specifically, for a specific overall median splicing timing *t* (i.e., the timing of lariat formation), we draw a median splicing timing *µ_1_* and *µ_2_* for each intron *1* and *2* from a lognormal distribution logN(*µ_G_*, *σ^2^*) with several possible variance parameters of *σ^2^* that yield a reasonable distribution of splicing timing (**Fig. S3L**). Next, for each median splicing timing *t* and distance ***ds***, we drew 1,000 samples each from T_1_ ∼ expon(*1/ µ_1_*) and T_2_ ∼ expon(1/µ_2_), where T_1_ and T_2_ represent individual splicing events, resulting in 1,000 pairs of simulated splicing timings (t_1,1_, …, t_1,1000_) and (t_2,1_, …, t_2,1000_). To calculate the number of in-order splicing events, we counted the number of intron pairs such that t_1,i_ < t_2,i_ + ***ds*** + t_s_, as this formula indicates that the timing of splicing of intron *1* is faster than the timing of splicing b plus ***ds***, which corresponds the delay in splicing of intron *2* imposed by the time to transcribe the exon and the downstream intron, and plus t_s_, which is fixed to 50 nts to roughly estimate the time of the second step of the downstream intron as it must have occurred to observe an out-of-order NLI. To calculate the number of out-of-order splicing events, we counted the number of intron pairs such that t_1,i_ > t_2,i_ + ***ds*** + t_s_. To calculate the proportion of in-order splicing, we divided the number of in-order splicing events by the sum of the in-order and out-of-order splicing events. Of note, the conclusion from our simulations did not change qualitatively when using reasonable variance parameters or different values (0 to 200 nts) for the timing of the second step of splicing.

To compare the empirical proportion of in-order splicing, we binned intron pairs according to the US-DS 3’ SS distance in 750 nt bins, from 500 nts to 10k nts. We then plotted the median proportion of in-order splicing of all intron pairs in each bin with at least 3 informative NLI reads. We then performed simulations as outlined above by using ***ds*** as the midpoint of each bin and by varying the overall median splicing timing. When we plotted the simulated data along with the empirical data for different variance parameters or different choices of splicing timing distributions; we found that the median splicing timing that best matched the empirical data was well under 1000 nts for most simulations, and under 1500 nts for all simulations with varying parameters.

To reduce the impact of read size bias on the empirical and simulated data, we calculated the proportion of in-order splicing using only in-order and out-of-order NLI reads that overlap with the downstream 5’ SSs, which resulted in reads with similar length distributions in the empirical data (**Fig. S3E, F**). Moreover, in the simulations, we restricted the calculation of the proportion of in-order splicing to intron pairs (x_a,i,_ x_b,i_) such that x_a,i_ or x_b,i_ are both less than 500 nts, given similar length constraints on the empirical CoLa-seq data.

### Sequence feature generation

For exons and their neighboring introns, sequence features such as length and nucleotide compositions (%A, %G, %C, %U, and %GC) were extracted and calculated using Bioconductor tools within R. For 5’ and 3’ splice sites, their scores were calculated using MaxEntscan algorithm (Yeo and Burge, 2004), and a 50-nt window surrounding each site was used to calculate nucleotide compositions and RNA secondary structure potentials (using viennaRNA (Lorenz et al., 2011)). The polypyrimidine tract (PPT) score is calculated using the script from Taggart et al., 2017). Binding energy was calculated using viennaRNA for U1 snRNA and the 5’ SS, U6 snRNA and the 5’ SS, and U2 snRNA and the BP. For branch points annotated by CoLa-seq, the following features were generated: branch point position, the percent usage of a branch point relative to alternative branch points, BP-3’ SS distance, and nucleotide composition and RNA secondary structure around branch points and in regions between the BP and 3’ SS. NLI read counts were also included in the 3’ SS-RNAP II distance models, because introns with greater numbers of NLI reads are more likely to sample earlier splicing events.

### XGBoost models to identify features that predict the order of splicing and the timing of in-order splicing

For the splicing order models, 23,324 introns pairs containing at least 10 CoLa-seq reads were used, and % in-order splicing, % concurrent splicing, and % out-of-order splicing were used as target variables. For the NLI 3’ SS-RNAP II distance models, 14,437 introns containing at least 10 in-order NLI reads were used and the minimum 3’ SS-RNAP II distance and the median 3’ SS-RNAP II distance were used as target variables. For each model, introns were randomly split into a training set and a test set in a 7:3 ratio. In the training set, when two features were more than 80% correlated (Pearson correlation), one of them was randomly removed. Afterwards, 22 features were used for the splicing-order models (**Table S3**) and 37 features were selected for the 3’ SS-RNAP II distance models. XGBoost (Chen and Guestrin, 2016) was used to train each model. The basic parameters for all models were as follows: objective=reg:squarederror, booster=gbtree, predictor=gpu_predictor, tree_method=gpu_hist. The hyperparameters for models were tuned using Hyperopt (Bergstra et al., 2013). The resulting hyper-parameters for the splicing-order models were as follows: max_depth=5, learning_rate=0.01, subsample=0.6, colsample_bynode=0.6, reg_alpha=0.5, reg_lambda=0.8, random_state=123, min_child_weight=60, n_estimators=600. The resulting hyper-parameters for the 3’ SS-RNAP II distance models were based on the default settings with the following changes: max_depth=5, learning_rate=0.01, subsample=0.6, reg_alpha=0.5, reg_lambda=0.8, random_state=123, min_child_weight = 200, n_estimators=550. The test set was used for model performance evaluation (metric: R-squared). Furthermore, 5-fold cross-validation was performed on the training set to derive the final averaged model performance as well as the variance of the performance (metric: R-squared). To evaluate the contribution of each feature within each model, SHAP values were computed and visualized using the SHAP package in python (Lundberg and Lee, 2017; Lundberg et al., 2018). In the 3’ SS-RNAP II models, downstream intron length is modeled but not considered further, because shorter downstream intron length correlates with shorter in-order NLI tails; this correlation emerges simply because when the downstream intron is short, in-order NLI tails would position RNAP II downstream of the downstream 5’ SS and thus directly compete with concurrent splicing and out-of-order splicing and deplete longer NLI tails.

### GC content calculation across splice sites for introns with different levels of in-order splicing

Intron pairs with different levels of in-order splicing were divided to three groups: low (≤20% in-order splicing), mid (40-60% in-order splicing), and high (≥80% in-order splicing). To calculate GC content around splice sites for each intron pair, 50 nts flanking each splice site were extracted. Sequences corresponding to splice site signals were not included in GC content calculation; specifically, for 5’ SSs, the last 3 nts in the exons and first 6 nts in the introns were removed, and for 3’ SSs, the last 3 nts in the introns and first 2 nts in the exons were removed. GC content was calculated at each nucleotide position with a 10 nt sliding window. Within each group, the average GC content and 95% confidence intervals were calculated at each position and plotted across splice sites of both upstream and downstream introns.

### Analysis of eCLIP-seq data in K562 cells from ENCODE

Bam files of input and IP for each RBP were downloaded from the ENCODE project website (https://www.encodeproject.org/; Van Nostrand et al., 2016). For each tested RBP (**Fig. 4C, 5H**), eCLIP-signals were processed using RBP-Maps (Yee et al., 2019) with default settings for introns with low in-order splicing (≤20%), introns with high in-order splicing (≥80%), early in-order introns (≥70% of 3’ SS-RNAP II distances ≤50 nts), and late in-order introns (≥70% of 3’ SS-RNAP II distances ≥200 nts). The resulting signals were further processed in R for visualization.

### De novo motif identification and motif enrichment analysis

To identify *de novo* motifs in specific regions of early splicing and late splicing introns, meme from MEME SUITE (Bailey et al., 2009; Tanaka et al., 2014) was used with a Markov model generated from the same regions of all introns containing at least 10 NLI reads. The identified motifs were further processed in R using universalmotif (Tremblay, 2019) and ggseqlogo for making sequence logos. The identified motifs were also fed into tomtom from MEME SUITE to search for enriched known motifs of RNA binding protein against the human CISBP-RNA database (Ray et al., 2013).

### Hexamer analysis

To identify enriched and depleted hexamers in specific regions of early splicing and late splicing introns, k-mer analysis was performed using transite package in R (Krismer et al., 2020).

### Quantification, statistical analysis, and plot generation

All quantification and statistical analyses were done in R and python. Analysis details can be found in figure legends and the result sections. All plots were prepared using data.table (Dowle and Srinivasan, 2019), ggpubr (Kassambara, 2020), cowplot (Wilke, 2019), patchwork (Pedersen, 2019), ggrepel (Slowikowski, 2020), or tidyverse tools (Wickham et al., 2019) in R, except for SHAP summary plots, which were made using SHAP and matplotlib (Hunter, 2007) in python.

## Supplementary files

**Table S1**. Branch points identified by lariat-seq

**Table S2.** Branch points identified by CoLa-seq

**Table S3.** Sequence features considered for modeling

**Table S4.** ncRNA depletion oligos

**Table S5.** Primer sequences used in library preparation

**Table S6.** Primer sequences used for PlaB validation experiments

## Supplementary figures

**Figure S1.**
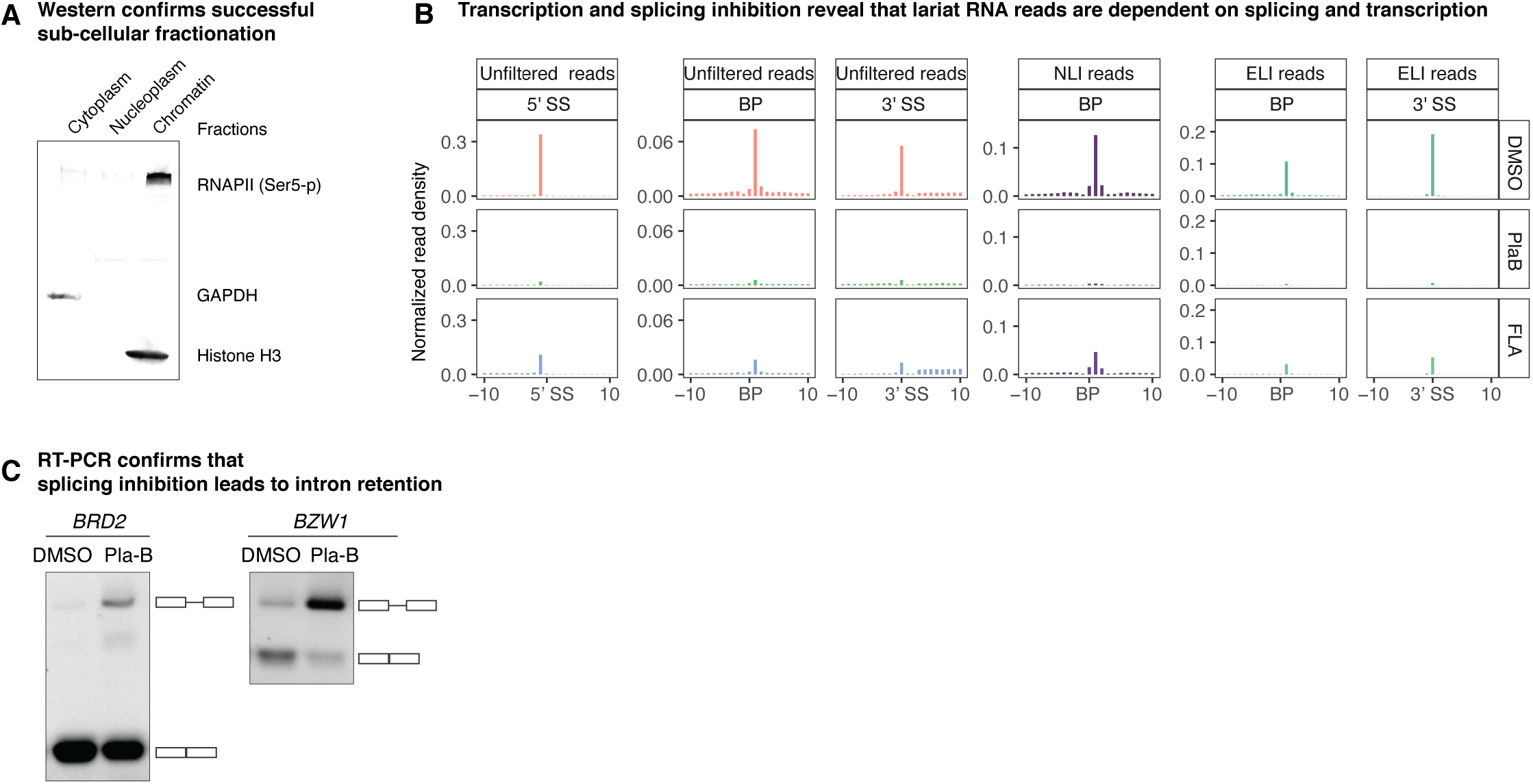
CoLa-seq reads are dependent on splicing and transcription; related to Figure 1. **A.** Western blot analysis of protein markers for each subcellular fraction. GAPDH was probed as a cytoplasmic marker. Histone H3 and RNAP II, phosphorylated at serine 5, are probed as chromatin markers. **B.** Bar plots illustrate i) the 5’ ends of CoLa-seq reads in regions around BPs, representing the lariat structure of either a NLI or an ELI and ii) the 3’ ends of CoLa-seq reads in regions around 5’SSs or 3’ SSs, representing a cleaved 5’ SS of a 5’ exon splicing intermediate or a cleaved 3’ SS of an ELI, respectively. CoLa-seq was performed on cells treated with DMSO (as a control), the splicing inhibitor pladienolide B (PlaB), or the transcription inhibitor flavopiridol (FLA). “Unfiltered”, uniquely mapped, deduplicated reads, before filtering for ELI or NLI reads; “NLI”, filtered, NLI reads; “ELI”, filtered, ELI reads. **C.** RT-PCR analysis of splicing efficiency for intron 4 of *BRD2* and intron 3 of *BZW1* from cells treated with DMSO or PlaB.

**Figure S2.**
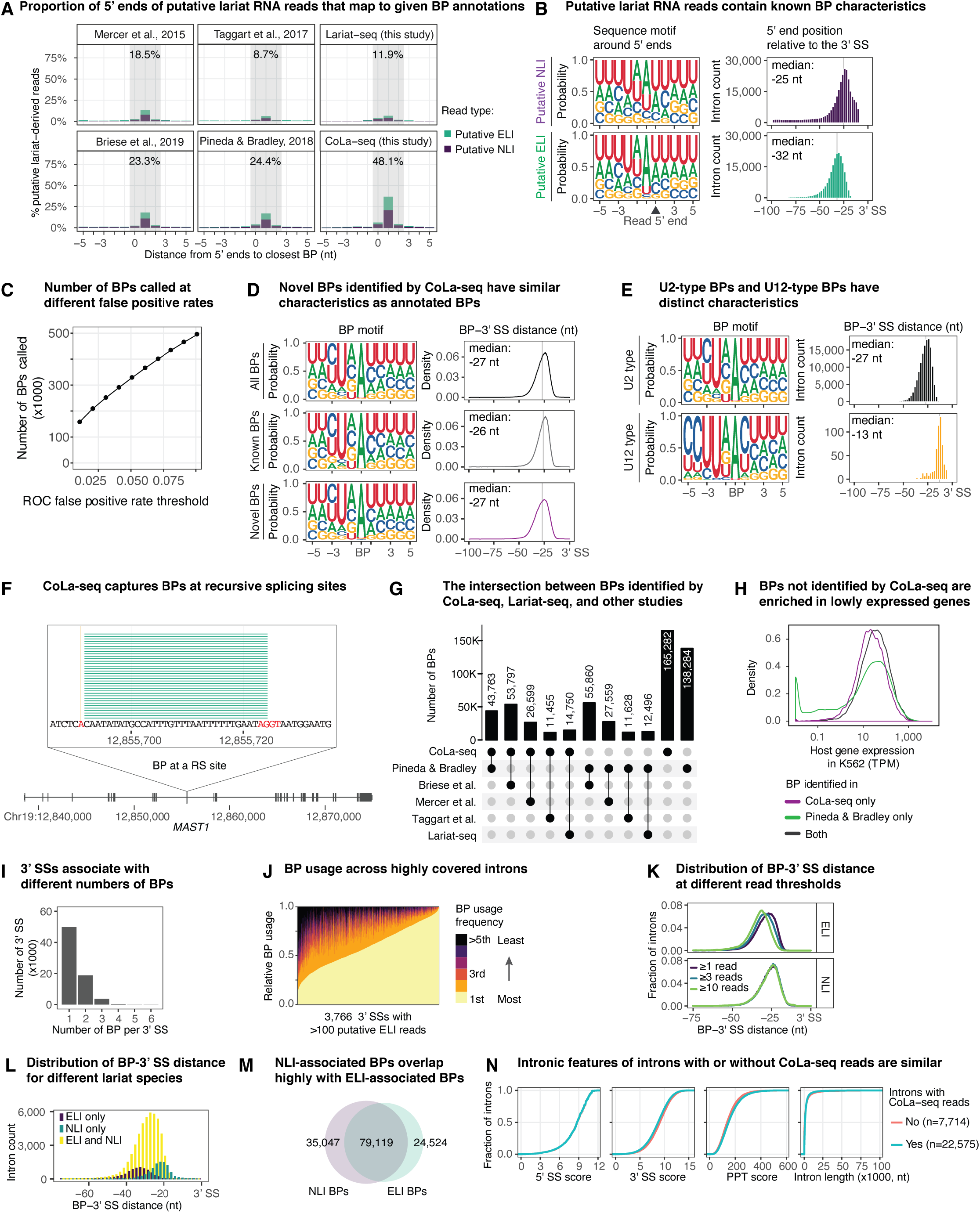
CoLa-seq maps human BPs to an unprecedented depth; related to Figure 1 and Figure 2. **A.** The fraction of 5’ ends of putative lariat RNA reads that terminate at or around a BP (+/- 5 nts), as determined by various BP annotation sources. Putative lariat RNA reads are uniquely mapped, deduplicated, reads whose 5’ ends are located within 100 nt upstream of a 3’ SS and whose 3’ ends cross a 3’ SS (putative NLI reads) or terminate at a 3’ SS (putative ELI reads). The grey zone indicates the sum of the % putative lariat-derived reads, whose 5’ ends align with the BP or 1-2 nts downstream. BPs identified by lariat-seq were obtained in this study (see Methods; **Table S1**). BPs identified by CoLa-seq were obtained in this study (see Methods; Fig. 1D**; Table S2**). **B.** Seqlogos and histograms illustrate the sequence motifs of BPs and the positions of BPs, relative to a 3’ SS, for putative NLI and putative ELI reads, defined as in Fig. S2A. **C.** The scatter plot illustrates the number of BPs identified at different false positive rates, as determined by the ROC (receiver operating characteristic curve) plot in Fig. 1D (right panel). A false positive rate of 0.01 was used in subsequent analysis. **D.** Seqlogos and histograms illustrate the sequence motif of BPs and the positions of BPs, relative to a 3’ SS, for all CoLa-seq-identified BPs (all), those that were previously identified (known; Briese et al., 2019; Mercer et al., 2015; Pineda and Bradley, 2018; Taggart et al., 2017), and those newly identified by CoLa-seq in this work (novel). Density reflects the number of BPs at a given BP-3’ss distance. **E.** Seqlogos and histograms illustrate the motif of BPs and the positions of BPs, relative to a 3’ SS, for CoLa-seq-identified BPs in U2-type and U12-type introns. **F.** ELI reads from a recursive splice site in *MAST1* are visualized. The recursive splice site at the 3’ end of the reads is denoted by AGGT in red, with the AG corresponding to a 3’ splice site and the GT corresponding to a 5’ splice site; the branch point at the 5’ end of the reads is denoted by an A, also in red. **G.** The upset plot illustrates the number of BPs intersecting between various pairs of BP annotations. **H.** Host-gene expression in K562 cells is plotted for BPs identified by CoLa-seq in K562 cells versus BPs identified in a previous study from a variety of cells types (Pineda and Bradley, 2018). Host gene expression in K562 cells was obtained from ENCODE (https://www.encodeproject.org). TPM, transcript per million. **I.** The bar plot illustrates the number of 3’ splice sites that are associated with the indicated numbers of BPs. **J.** The stacked bar plot illustrates the relative BP usage for each of 3,766 3’ SS with >100 putative ELI reads. BP usage was assessed independent of CoLa-seq BP annotations by simply quantifying the 5’ ends of putative ELI reads, defined as terminating at an annotated 3’ SS. **K.** The detection of shorter BP-3’ SS distances by ELI reads is limited by low read numbers. Density plots illustrate BP-3’ SS distance distributions for BPs identified by different thresholds of ELI or NLI reads. **L.** Distribution of BP-3’ SS distance for different lariat species captured by CoLa-seq. Histograms illustrate BP-3’ SS distance distributions for BPs that were observed by both NLI and ELI reads (ELI and NLI), by far the largest class; only by ELI reads (ELI only); and only by NLI reads (NLI only). BPs that have at least 10 reads are used. Note that short BP-3’ SS distances for ELI reads are underrepresented, relative to other ELI reads and NLI reads, likely because of the lower-bound, molecular weight cutoff used during the preparation of CoLa-seq libraries. Long BP-3’ SS distances for ELI reads are overrepresented, relative to other ELI reads and NLI reads, likely because these reads are short (but above the lower-bound, molecular weight cutoff) and favored in cDNA synthesis and/or sequencing. **M.** The Venn diagram illustrates the robust overlap between NLI-associated BPs and ELI-associated BPs. **N.** Cumulative plots illustrate that introns yielding CoLa-seq reads are similar to introns without CoLa-seq reads, in terms of the distribution of 5’ SS scores, 3’ SS scores, PPT scores, and intron lengths. A subtle increase in PPT score, a subtle decrease in 3’ SS score, and a subtle decrease in intron length for introns yielding CoLa-seq reads is consistent with a role for the PPT in early NLI formation (see text), a role for the 3’ SS in converting NLI to mRNA, and a role for shorter introns in promoting early splicing, all of which likely favor NLI capture by CoLa-seq. For simplicity, only constitutive introns were analyzed.

**Figure S3.**
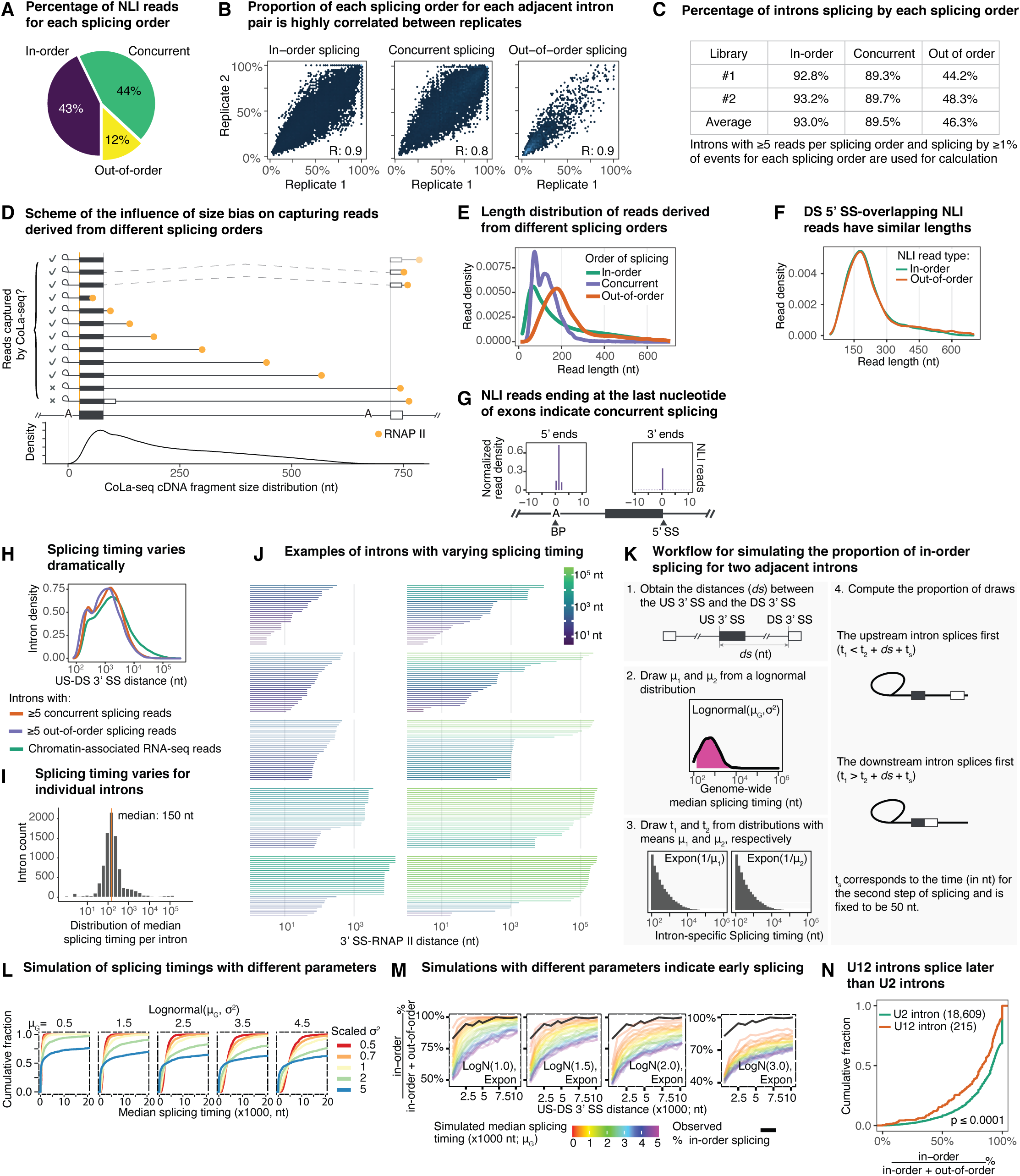
Nascent lariat intermediates reveal prevalent in-order, out-of-order, and concurrent splicing; related to Figure 3. **A.** The pie chart illustrates the percentage of all NLI reads corresponding to each order of splicing, as observed in the CoLa-seq library. Note that although concurrent NLI species could in principle reflect a back splicing intermediate, the exons and flanking introns of these species are generally too short to support back splicing, and the importance of features at the 3’ end of the downstream intron to concurrent NLI levels identified by modeling (Fig. 5) is inconsistent with back splicing but consistent with concurrent splicing. **B.** Scatter plots illustrate that the percentage of events of each splicing order for each pair of adjacent introns is highly reproducible between two biological replicates. **C.** Percentage of introns splicing by each order of splicing, in two biological replicates. **D.** The impact of length selection and length bias in the capture of NLIs by CoLa-seq. The final CoLa-seq libraries were isolated by gel purification with an upper insert size cutoff of ≤650 nts (top ten instances). Consequently, in-order splicing and out-of-order NLIs with longer tails, representing later splicing events, are not captured (bottom two instances). Further, due to biases in Illumina sequencing disfavoring longer molecules, NLIs with shorter tails are more highly represented as implied by the density plot of CoLa-seq read lengths (bottom density plot). Concurrent splicing and out-of-order splicing NLIs can be captured if the cDNA is small and thus reflect very late splicing events when the downstream intron is long (top three instances), despite the length restriction and bias, as reflected by the genomic position of RNAP II during concurrent splicing (faded RNAP II) and out-of-order splicing. **E.** The density plot illustrates the size distribution of read lengths belonging to the different classes of splicing order. **F.** The density plots illustrate that filtering NLI reads for only those extending beyond the downstream exon eliminates the difference in length distribution between in-order and out-of-order NLI reads, as illustrated in panel **E**, thereby enabling a direct quantitative comparison between the read numbers for in-order splicing versus out-of-order splicing. **G.** A meta-analysis of the 3’ ends of all NLI reads aligned to the downstream 5’ SS reveals a prominent peak at the last nucleotide of exons, indicative of concurrent splicing. For reference, the 5’ ends of concurrent NLIs are also separately aligned to annotated BPs. Bar plots illustrate the normalized read density of 5’ ends around BPs or of 3’ ends around the downstream 5’ SS. **H.** The density plot illustrates the distribution of US-DS 3’ SS distances for introns that are detected by chromatin-associated RNA-seq, introns having ≥ 5 concurrent splicing reads, or introns having ≥5 out-of-order splicing reads. For the introns yielding out-of-order splicing reads, the US-DS 3’ SS distances vary from 93 nts to 1,115,021 nts, and for introns yielding concurrent splicing reads, the US-DS 3’ SS distances vary from 93 nts to 178,779 nts, underscoring dramatic variation in splicing timing. **I.** The histogram shows the distribution of median 3’ SS-RNAP II distance per intron. Assuming first-order kinetics, the median 3’ SS-RNAP II distance corresponds to the splicing half-life. The vertical yellow line indicates the median of the distribution. **J.** Examples illustrate the in-order and out-of-order NLI reads for 15 randomly selected introns (30 reads were sampled from each intron for visualization purposes). The example in the lower right shows an intron yielding a few in-order splicing reads; more out-of-order splicing reads, resulting from earlier splicing of the downstream intron; and even more out-of-order splicing reads resulting from earlier splicing of the first two downstream introns. **K.** The workflow of % in-order splicing simulation. (1) For each intron pair for which we empirically measured % in-order splicing, we calculated the distance from the US 3’ SS to DS 3’ SS (*ds*) in nts. The % in-order splicing expected was simulated under various scenarios as follows. Assuming the splicing rates of individual introns in intron pairs vary and are independent from each other. (2) We randomly drew different splicing timings for each intron (μ_1_ and μ_2_) from a lognormal distribution: logN(μ_G_, σ^2^), where μ_G_ represents the median splicing half-life of introns genome-wide, and μ_1_ and μ_2_ represent the median splicing timings of individual introns in intron pairs; σ^2^ reflects the variance and consequently the shape of the distribution. (3) We then repeatedly drew individual splicing timings (the time of forming a lariat intermediate) for each intron (t_1_ and t_2_) using the splicing rates determined in (2), assuming that splicing timing of individual introns can be modeled as an exponential distribution. (4) Finally, we assessed the proportion of draws wherein t_1_ and t_2_ would indicate that splicing occurred in-order versus out-of-order. **L.** The plots visualize the cumulative density of aggregated median splicing timings across introns following different lognormal distributions for step 2 in the simulation (**Fig. S3K**). The variable μ_G_ is the location parameter and can be interpreted as the genome-wide median of median splicing timings for individual introns. Scaled sigma squared (σ^2^) is the shape parameter and is a measure of how varied splicing rates are across different introns. Because our simulations are sensitive to the assumed value of σ^2^ (Fig. 3J**; Fig. S3M**), we plot the genome-wide distribution of splicing times for different values of σ^2^, in addition to varying μ_G_. **M.** The results of simulations under different scenarios with different assumed values of μ_G_ (colored lines) as well as of σ^2^, as indicated in each panel as logN(σ^2^). Simulations assumed that splicing timing for individual introns follows an exponential distribution. The observed % in-order splicing as a function of US-DS 3’ SS distance is shown in black. In all simulated scenarios examined, the median, genome-side splicing rate, μ_G_, that best matches the data indicates that splicing generally happens when RNAP II is within 1000 nts downstream of the 3’ SS. **N.** The cumulative plot illustrates that U2 introns have higher levels of in-order splicing, implying earlier splicing than U12 introns.

**Figure S4.**
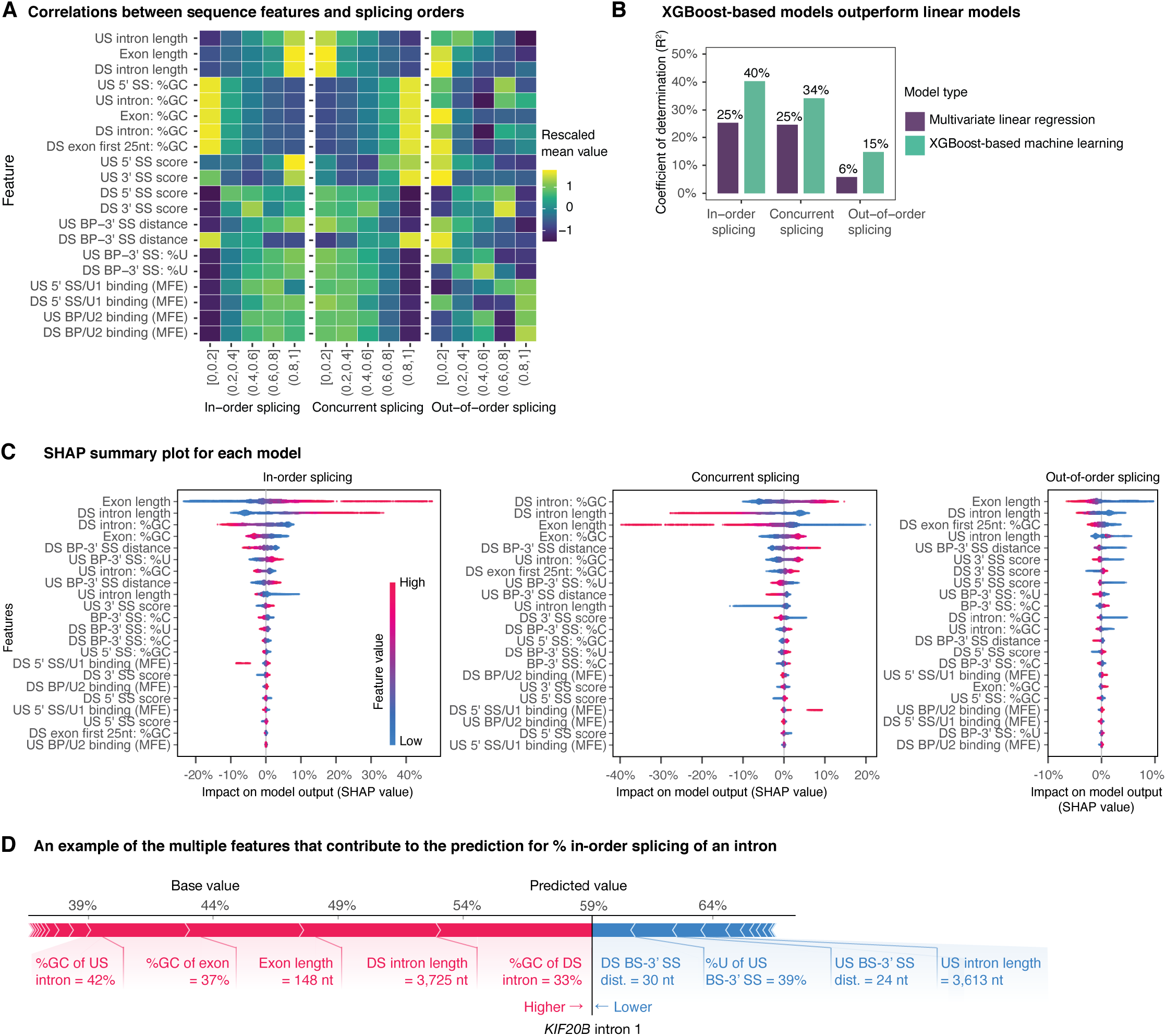
Intronic elements, gene architecture, and genomic context predict the order of co-transcriptional splicing; related to Figure 4. **A.** The heatmap illustrates the relationship between the indicated features and the percentage of in-order splicing, concurrent splicing, or out-of-order splicing. The fraction of an order of splicing for each intron is sorted into one of five bins, as indicated. Feature values are normalized within each feature. **B.** Bar plots illustrate that the model performance of each model, measured by the coefficient of determination (R^2^). **C.** SHAP summary plots illustrate the impact of each feature on the prediction of the percentage of in-order splicing, concurrent splicing, or out-of-order splicing, for each intron pair detected. Features are ordered from high to low by the sum of the absolute SHAP values of each feature for each intron pair. **D.** The force plot visualizes the feature contribution to the prediction of % in-order splicing for one randomly sampled intron as an example. The color schemes and the definition of “base value” are the same as in Fig. 4B.

**Figure S5.**
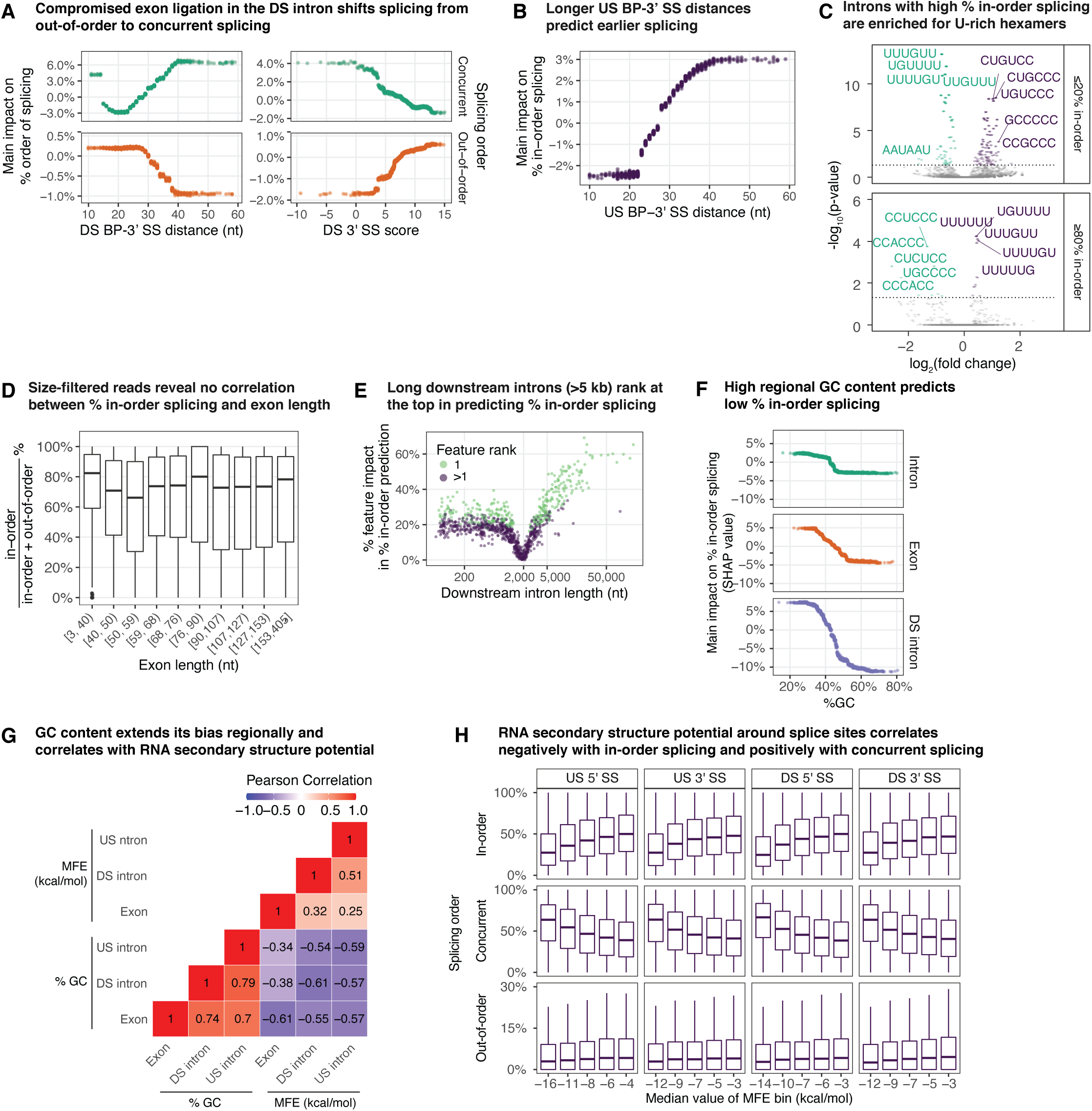
Features contribute to the prediction in the splicing-order models; related to Figure 4. **A.** Feature values in the downstream (DS) intron that delay the transition from splicing intermediates to splicing products shift the distribution of splicing events from out-of-order splicing to concurrent splicing, from species in which the downstream intron is spliced to species in which the downstream intron is at the intermediate stage of splicing. Scatter plots illustrate the impact (i.e., main SHAP value) of the downstream BP-3’ SS distance and the downstream 3’ SS strength on the prediction of the percentage of concurrent and out-of-order splicing. **B.** The scatter plot illustrates the main impact of the upstream BP-3’ SS distance on the prediction of the percentage of in-order splicing. Although a longer BP-3’ SS can delay exon ligation and accumulate lariat intermediate (e.g., see panel **A** and legend), the lariat intermediate state of the upstream intron is common to all splicing orders, so changes in the lifetime of this species would not be expected to favor one order over another. Consequently, the favorable impact of BP-3’ SS distance specifically for in-order splicing may reflect the greater likelihood of a strong PPT and thus strong U2AF binding, which predicts early, in-order splicing (Fig. 4A**, C**). **C.** U-rich hexamers are enriched in the BP-3’ SS region of introns with ≥80% in-order splicing and depleted in the same region of introns with ≤20 % in-order splicing. Significantly-enriched hexamers are in purple, and significantly-depleted hexamers are in green (Benjamini-Hochberg corrected p-value <0.05). P-values (-log_10_(p-value)) are plotted as a function of the hexamer fold enrichment (log_2_(fold change)). **D.** The box plot visualizes % in-order splicing as a function of exon length. The % in-order splicing is calculated as (in-order NLI reads) / (in-order NLI reads + out-of-order NLI reads) x 100. Only size-filtered in-order and out-of-order NLI reads are used (see **Fig. S3F**). Samples were divided to five groups with the same number of introns based on exon length. **E.** The contribution of downstream intron length to the prediction of % in-order splicing for each intron is plotted as a function of intron length. When downstream intron length is the top feature (“1”) in the prediction for an intron, the data point for the intron is labeled green; otherwise, when downstream intron length is not the top feature (“>1”) in the prediction for an intron, the data point for the intron is labeled dark purple. **F.** Scatter plots illustrate that higher regional GC contents predict lower levels of in-order splicing. The plots visualize the main impact SHAP value for GC content on the prediction of % in-order splicing, as a function of GC content in the upstream intron, the exon, and the downstream intron. **G.** The heatmap shows that the GC content of an intron, the downstream exon, and the downstream intron, in a single gene, are highly correlated and that these GC contents correlate with regional secondary structure potential, as calculated by minimum free energy (MFE; kcal/mol). Pearson correlations are indicated. When an intron is longer than 200 nts, its first 100 and last 100 nts are used, instead of the entire intron, for MFE calculation. **H.** Boxplots show that strong secondary potential around both the upstream and downstream splice sites correlate negatively with in-order and out-of-order splicing and positively with concurrent splicing. The percent in-order splicing, concurrent splicing, and out-of-order splicing is shown as a function of MFE at the upstream and downstream 5’ and 3’ SSs. Samples were divided to five groups with the same number of introns. The median value of each group is indicated on the x-axis.

**Figure S6.**
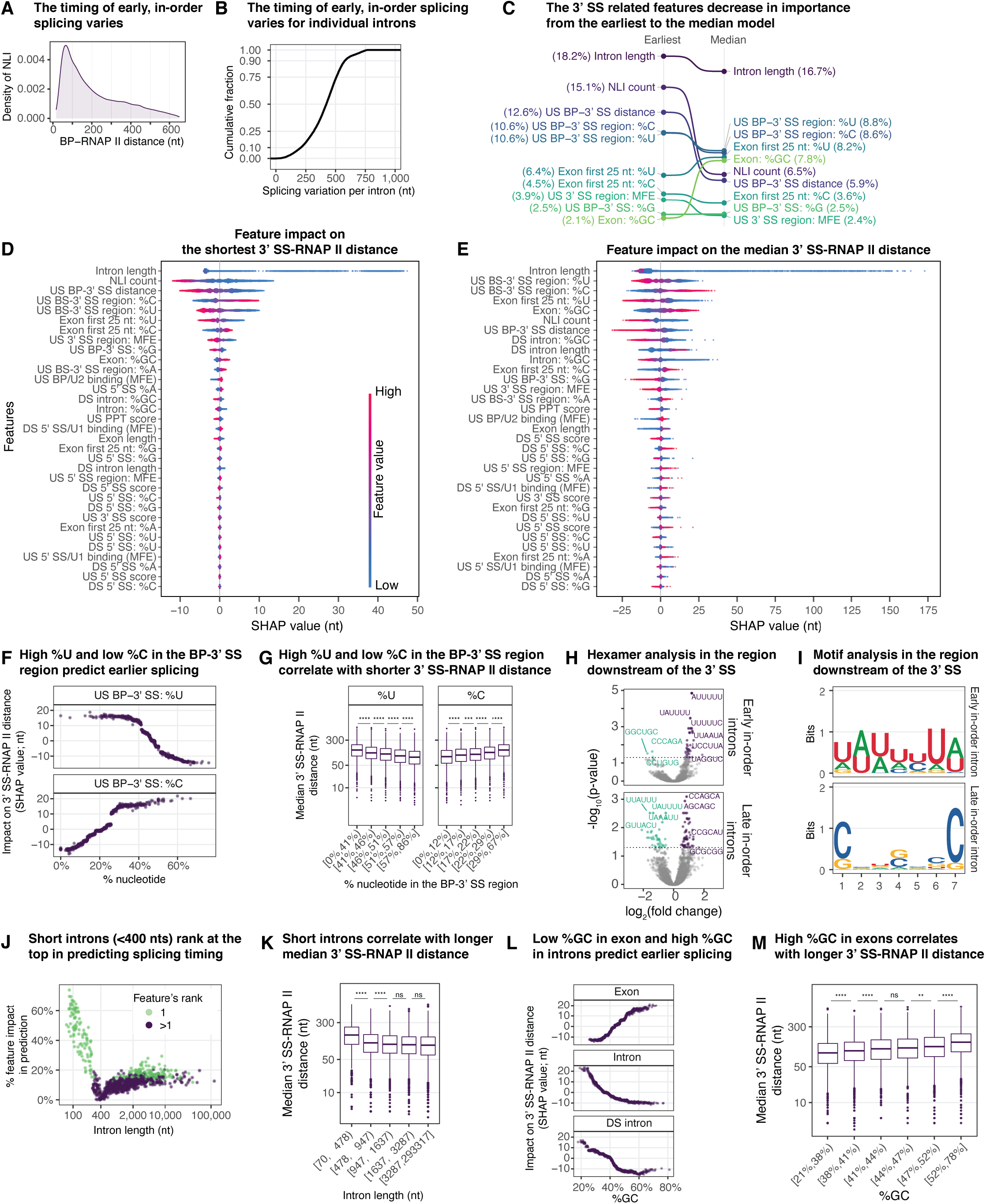
Specific features are associated with early in-order splicing; related to Figure 5. **A.** The density plot illustrates the BP-RNAP II distance distribution of all in-order NLI reads. **B.** The cumulative plot shows the distribution of splicing variation per intron. For each intron, its splicing variation is defined as the difference between its shortest and its longest 3’ SS-RNAP II distance from its in-order NLI reads. In total, 14,438 introns (≥10 in-order NLI reads) are used for plotting. **C.** Feature importance changes from the earliest model to the median model. The % contribution for a feature is shown for the earliest 3’SS-RNAP II distance model and the median 3’SS-RNAP II distance model. The connecting lines indicate whether a feature increases or decreases in its predictive value for the median model, relative to the earliest model **D.** The SHAP summary plot illustrates the impact of each feature for each intron, as determined by the earliest model. Features are ranked from high to low SHAP values by the sum of absolute SHAP values of each feature for each intron and plotted as a function of SHAP value, where the SHAP value indicates predicted impact on the shortest 3’ SS-RNAP II distance (nt). The magnitude of a feature is indicated by a heatmap, with red reflecting high values and blue reflecting low values. **E.** The SHAP summary plot illustrates the impact of each feature for each intron, as determined by the median model. Values are plotted as in panel **D**. **F.** Scatter plots illustrate the predicted impact of U or C content in the BP-3’ SS region, as determined by the median model. The impact is plotted as SHAP values, in terms of nucleotides, as a function of nucleotide composition. **G.** Box plots illustrate the relationship between U or C content in the BP-3’ SS region and median 3’ SS-RNAP II distance. Samples are split into five bins, with the same number of introns, based on either %U or %C in the BP-3’ SS region. ****, P ≤ 0.0001; ***, P ≤ 0.001. **H.** The scatter plot illustrates the results of a hexamer enrichment analysis in the 50-nt region downstream of the 3’ SS in early or late in-order introns. Significantly enriched hexamers are in purple, and significantly depleted hexamers are in green (Benjamini-Hochberg corrected p-value <0.05). P-values (-log_10_(p-value)) are plotted as a function of the hexamer fold enrichment (log_2_(fold change)). **I.** Seqlogos show the top motifs in the 50-nt region downstream of the 3’ SS for early or late in-order introns. **J.** The scatter plot illustrates that, with short introns, intron length outweighs other features in predicting splicing timing in terms of 3’ SS-RNAP II distance, as determined by the median model; note that the earliest model shows the same trend. The contribution of intron length to the prediction of splicing timing for each intron is plotted as a function of intron length. When intron length is the top feature (“1”) in the prediction for an intron, the data point for the intron is labeled green; otherwise, when intron length is not the top feature (“>1”) in the prediction for an intron, the data point for the intron is labeled dark purple. **K.** The box plot illustrates the relationship between intron length and median 3’ SS-RNAP II distance. Median 3’ SS-RNAP II distance is plotted as a function of intron length, binned as noted. ****, P ≤ 0.0001; ns, not significant. **L.** Scatter plots illustrate the impact, the SHAP value, of predicting 3’ SS-RNAP II distance (nt) as a function of GC content of the NLI intron, the downstream exon, and the downstream intron, as determined by the median model. **M.** The box plot illustrates median 3’ SS-RNAP II distance as a function of GC content, binned as indicated. ****, P ≤ 0.0001; **, P ≤ 0.01; ns, not significant.

**Figure S7.**
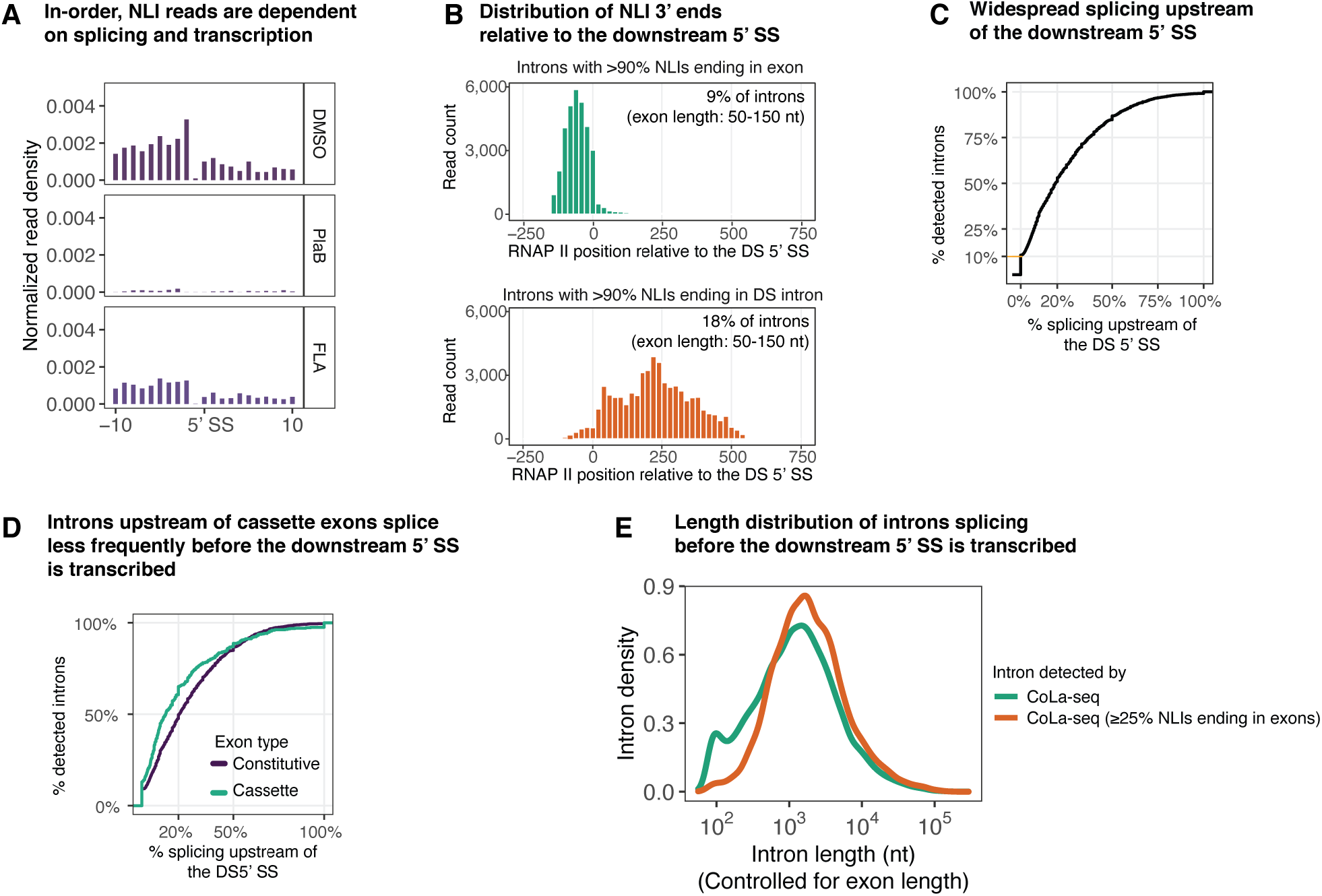
Early co-transcriptional lariat formation indicates widespread usage of intron definition; related to Figure 6. **A.** In-order, NLI reads captured by CoLa-seq depend on spliceosome assembly and transcription. Bar meta-plots illustrate the 3’ ends of in-order, NLI reads around the downstream 5’ SSs. CoLa-seq was performed on cells treated with DMSO, the splicing inhibitor pladienolide B (PlaB), or the transcription inhibitor flavopiridol (FLA). The data derive from the same experiment that yielded the data illustrated in **Fig. S1B.** **B.** Histograms illustrate the distribution of in-order NLI 3’ ends relative to the downstream 5’ SS for introns with ≥90% NLIs ending in either the exon (top) or the downstream intron (bottom). To control for potential bias of the exon length, only introns with exons ranging from 50 to 150 nts are used. **C.** The cumulative plot illustrates the percentage of introns, detected by CoLa-seq, as a function of the percentage of NLIs with 3’ ends upstream of the downstream 5’ splice site for each intron, implying independence of such splicing on the downstream 5’ splice site, if not also exon definition. The percentage of splicing that occurred upstream of the DS 5’ SS per intron is calculated as follows: (in-order NLI reads upstream of the DS 5’ SS)/(in-order NLI reads + concurrent NLI reads + out-of-order NLI reads) x 100. **D.** Introns upstream of cassette exons, relative to constitutive exons, splice less frequently before the downstream 5’ splice site is transcribed. The percentage of introns detected by CoLa-seq are plotted as a function of the cumulative percentage of the corresponding NLI 3’ ends terminating upstream of the downstream 5’ SS (i.e., independent of canonical exon definition). Green, introns upstream of constitutive exons; purple, introns upstream of cassette exons. The % splicing upstream of the DS 5’ SS is calculated the same way as in **C**. **E.** Distribution of intron lengths for introns splicing upstream of the downstream 5’ SS, controlled for exon length. Exon-definition-independent splicing events are observed in introns with wide-ranging lengths, including long introns. The density of introns is plotted as a function of intron length. Introns for which ≥25% of NLIs terminate upstream of the downstream 5’ splice site are plotted in orange; for comparison, all introns detected by CoLa-seq (≥5 reads) are plotted in green. Only introns associated with exons ranging from 50 and 150 nts are plotted to minimize the impact of length biases.

**Figure S8.**
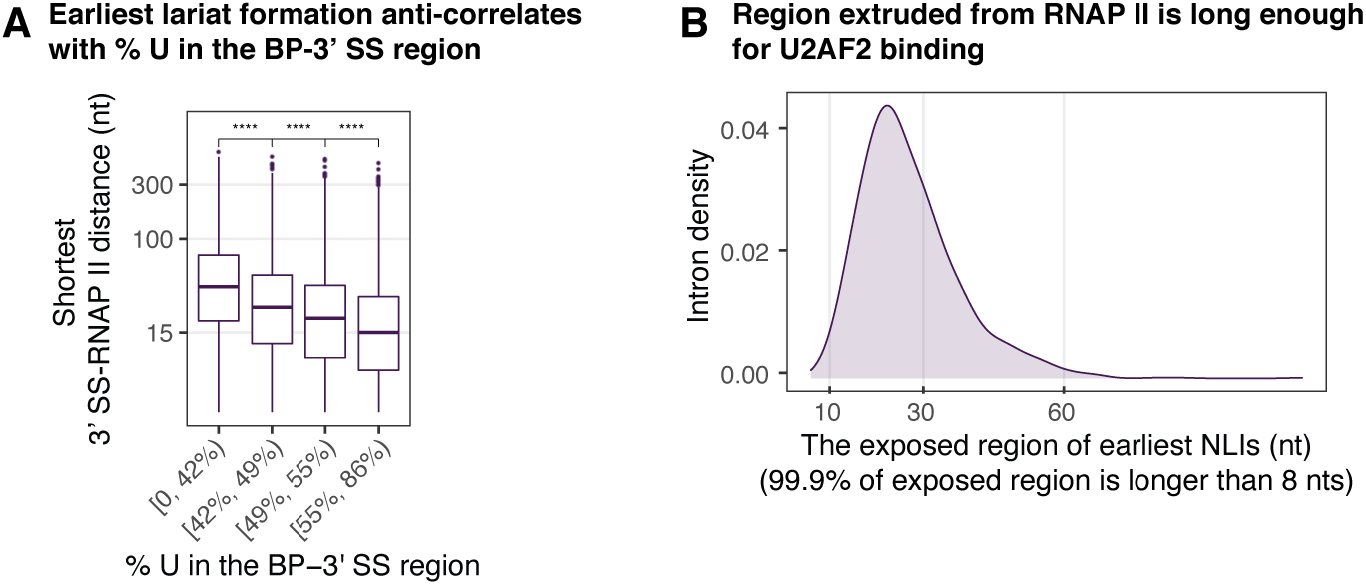
Related to Figure 7. **A.** The box plot displays the shortest 3’ SS-RNAP II distance of NLIs as a function of the percentage of U in the BP-3’ SS region, binned as indicated. ****, P ≤ 0.0001.**B.** An intron density plot is shown for the length of the exposed region downstream of the BP for the shortest NLI of each intron, expressed in nucleotides.

## References

Amit, M., Donyo, M., Hollander, D., Goren, A., Kim, E., Gelfman, S., Lev-Maor, G., Burstein, D., Schwartz, S., Postolsky, B., et al. (2012). Differential GC Content between Exons and Introns Establishes Distinct Strategies of Splice-Site Recognition. Cell Rep. 1, 543–556.

Bailey, T.L., Boden, M., Buske, F.A., Frith, M., Grant, C.E., Clementi, L., Ren, J., Li, W.W., and Noble, W.S. (2009). MEME Suite: tools for motif discovery and searching. Nucleic Acids Res. 37, W202– W208.

Berget, S.M. (1995). Exon Recognition in Vertebrate Splicing. J. Biol. Chem. 270, 2411–2414.

Bergstra, J., Yamins, D., and Cox, D.D. (2013). Making a science of model search: hyperparameter optimization in hundreds of dimensions for vision architectures. In Proceedings of the 30th International Conference on International Conference on Machine Learning - Volume 28, (Atlanta, GA, USA: JMLR.org), p. I-115–I–123.

Bernecky, C., Plitzko, J.M., and Cramer, P. (2017). Structure of a transcribing RNA polymerase II– DSIF complex reveals a multidentate DNA–RNA clamp. Nat. Struct. Mol. Biol. 24, 809–815.

Beyer, A.L., and Osheim, Y.N. (1988). Splice site selection, rate of splicing, and alternative splicing on nascent transcripts. Genes Dev. 2, 754–765.

Bradley, R.K., Merkin, J., Lambert, N.J., and Burge, C.B. (2012). Alternative Splicing of RNA Triplets Is Often Regulated and Accelerates Proteome Evolution. PLoS Biol. 10, e1001229.

Braunschweig, U., Gueroussov, S., Plocik, A.M., Graveley, B.R., and Blencowe, B.J. (2013). Dynamic Integration of Splicing within Gene Regulatory Pathways. Cell 152, 1252–1269.

Bray, N.L., Pimentel, H., Melsted, P., and Pachter, L. (2016). Near-optimal probabilistic RNA-seq quantification. Nat. Biotechnol. 34, 525–527.

Briese, M., Haberman, N., Sibley, C.R., Faraway, R., Elser, A.S., Chakrabarti, A.M., Wang, Z., König, J., Perera, D., Wickramasinghe, V.O., et al. (2019). A systems view of spliceosomal assembly and branchpoints with iCLIP. Nat. Struct. Mol. Biol. 26, 930–940.

Carrillo Oesterreich, F., Herzel, L., Straube, K., Hujer, K., Howard, J., and Neugebauer, K.M. (2016). Splicing of Nascent RNA Coincides with Intron Exit from RNA Polymerase II. Cell 165, 372–381.

Chen, M., and Manley, J.L. (2009). Mechanisms of alternative splicing regulation: insights from molecular and genomics approaches. Nat. Rev. Mol. Cell Biol. 10, 741–754.

Chen, T., and Guestrin, C. (2016). XGBoost: A Scalable Tree Boosting System. Proc. 22nd ACM SIGKDD Int. Conf. Knowl. Discov. Data Min. 785–794.

Chen, S., Zhou, Y., Chen, Y., and Gu, J. (2018a). fastp: an ultra-fast all-in-one FASTQ preprocessor. Bioinformatics 34, i884–i890.

Chen, W., Moore, J., Ozadam, H., Shulha, H.P., Rhind, N., Weng, Z., and Moore, M.J. (2018b). Transcriptome-wide Interrogation of the Functional Intronome by Spliceosome Profiling. Cell 173, 1031–1044.e13.

Christofori, G., Frendewey, D., and Keller, W. (1987). Two spliceosomes can form simultaneously and independently on synthetic double-intron messenger RNA precursors. EMBO J. 6, 1747–1755.

Costantini, M., and Musto, H. (2017). The Isochores as a Fundamental Level of Genome Structure and Organization: A General Overview. J. Mol. Evol. 84, 93–103.

Costantini, M., Clay, O., Auletta, F., and Bernardi, G. (2006). An isochore map of human chromosomes. Genome Res. 16, 536–541.

Darman, R.B., Seiler, M., Agrawal, A.A., Lim, K.H., Peng, S., Aird, D., Bailey, S.L., Bhavsar, E.B., Chan, B., Colla, S., et al. (2015). Cancer-Associated SF3B1 Hotspot Mutations Induce Cryptic 3′ Splice Site Selection through Use of a Different Branch Point. Cell Rep. 13, 1033–1045.

Das, R., Yu, J., Zhang, Z., Gygi, M.P., Krainer, A.R., Gygi, S.P., and Reed, R. (2007). SR proteins function in coupling RNAP II transcription to pre-mRNA splicing. Mol. Cell 26, 867–881.

De Conti, L., Baralle, M., and Buratti, E. (2013). Exon and intron definition in pre-mRNA splicing. Wiley Interdiscip. Rev. RNA 4, 49–60.

Dobin, A., Davis, C.A., Schlesinger, F., Drenkow, J., Zaleski, C., Jha, S., Batut, P., Chaisson, M., and Gingeras, T.R. (2013). STAR: ultrafast universal RNA-seq aligner. Bioinformatics 29, 15–21.

Dowle, M., and Srinivasan, A. (2019). data.table: Extension of ‘data.fram’.

Drexler, H.L., Choquet, K., and Churchman, L.S. (2020). Splicing Kinetics and Coordination Revealed by Direct Nascent RNA Sequencing through Nanopores. Mol. Cell 77, 985–998.e8.

Feng, Y.-Y., Ramu, A., Cotto, K.C., Skidmore, Z.L., Kunisaki, J., Conrad, D.F., Lin, Y., Chapman, W.C., Uppaluri, R., Govindan, R., et al. (2018). RegTools: Integrated analysis of genomic and transcriptomic data for discovery of splicing variants in cancer. BioRxiv 436634.

Fica, S.M., Tuttle, N., Novak, T., Li, N.-S., Lu, J., Koodathingal, P., Dai, Q., Staley, J.P., and Piccirilli, J.A. (2013). RNA catalyses nuclear pre-mRNA splicing. Nature 503, 229–234.

Fox-Walsh, K.L., Dou, Y., Lam, B.J., Hung, S., Baldi, P.F., and Hertel, K.J. (2005). The architecture of pre-mRNAs affects mechanisms of splice-site pairing. Proc. Natl. Acad. Sci. 102, 16176–16181.

Gao, K., Masuda, A., Matsuura, T., and Ohno, K. (2008). Human branch point consensus sequence is yUnAy. Nucleic Acids Res. 36, 2257–2267.

Guth, S., Martínez, C., Gaur, R.K., and Valcárcel, J. (1999). Evidence for Substrate-Specific Requirement of the Splicing Factor U2AF35 and for Its Function after Polypyrimidine Tract Recognition by U2AF65. Mol. Cell. Biol. 19, 8263–8271.

Herzel, L., and Neugebauer, K.M. (2015). Quantification of co-transcriptional splicing from RNA-Seq data. Methods San Diego Calif 85, 36–43.

Herzel, L., Ottoz, D.S.M., Alpert, T., and Neugebauer, K.M. (2017). Splicing and transcription touch base: co-transcriptional spliceosome assembly and function. Nat. Rev. Mol. Cell Biol. 5, 347.

Herzel, L., Straube, K., and Neugebauer, K.M. (2018). Long-read sequencing of nascent RNA reveals coupling among RNA processing events. Genome Res. 28, 1008–1019.

Hollander, D., Naftelberg, S., Lev-Maor, G., Kornblihtt, A.R., and Ast, G. (2016). How Are Short Exons Flanked by Long Introns Defined and Committed to Splicing? Trends Genet. 32, 596–606.

Huber, W., Carey, V.J., Gentleman, R., Anders, S., Carlson, M., Carvalho, B.S., Bravo, H.C., Davis, S., Gatto, L., Girke, T., et al. (2015). Orchestrating high-throughput genomic analysis with Bioconductor. Nat. Methods 12, 115–121.

Hunter, J.D. (2007). Matplotlib: A 2D Graphics Environment. Comput. Sci. Eng. 9, 90–95.

Jonkers, I., Kwak, H., and Lis, J.T. (2014). Genome-wide dynamics of Pol II elongation and its interplay with promoter proximal pausing, chromatin, and exons. ELife 3, e02407.

Kang, H.-S., Sánchez-Rico, C., Ebersberger, S., Sutandy, F.X.R., Busch, A., Welte, T., Stehle, R., Hipp, C., Schulz, L., Buchbender, A., et al. (2020). An autoinhibitory intramolecular interaction proof-reads RNA recognition by the essential splicing factor U2AF2. Proc. Natl. Acad. Sci. 117, 7140–7149.

Kassambara, A. (2020). ggpubr: “ggplot2” Based Publication Ready Plots.

Kastner, B., Will, C.L., Stark, H., and Lührmann, R. (2019). Structural Insights into Nuclear pre-mRNA Splicing in Higher Eukaryotes. Cold Spring Harb. Perspect. Biol. a032417.

Kessler, O., Jiang, Y., and Chasin, L.A. (1993). Order of intron removal during splicing of endogenous adenine phosphoribosyltransferase and dihydrofolate reductase pre-mRNA. Mol. Cell. Biol. 13, 6211– 6222.

Kim, S.W., Taggart, A.J., Heintzelman, C., Cygan, K.J., Hull, C.G., Wang, J., Shrestha, B., and Fairbrother, W.G. (2017). Widespread intra-dependencies in the removal of introns from human transcripts. Nucleic Acids Res. 45, 9503–9513.

Krismer, K., Bird, M.A., Varmeh, S., Handly, E.D., Gattinger, A., Bernwinkler, T., Anderson, D.A., Heinzel, A., Joughin, B.A., Kong, Y.W., et al. (2020). Transite: A Computational Motif-Based Analysis Platform That Identifies RNA-Binding Proteins Modulating Changes in Gene Expression. Cell Rep. 32, 108064.

Leader, Y., Lev Maor, G., Sorek, M., Shayevitch, R., Hussein, M., Hameiri, O., Tammer, L., Zonszain, J., Keydar, I., Hollander, D., et al. (2021). The upstream 5′ splice site remains associated to the transcription machinery during intron synthesis. Nat. Commun. 12, 4545.

Lemaire, S., Fontrodona, N., Aubé, F., Claude, J.-B., Polvèche, H., Modolo, L., Bourgeois, C.F., Mortreux, F., and Auboeuf, D. (2019). Characterizing the interplay between gene nucleotide composition bias and splicing. Genome Biol. 20, 259.

Li, H., Handsaker, B., Wysoker, A., Fennell, T., Ruan, J., Homer, N., Marth, G., Abecasis, G., Durbin, R., and Subgroup, 1000 Genome Project Data Processing (2009). The Sequence Alignment/Map format and SAMtools. Bioinformatics 25, 2078–2079.

Li, Y.I., Knowles, D.A., Humphrey, J., Barbeira, A.N., Dickinson, S.P., Im, H.K., and Pritchard, J.K. (2018). Annotation-free quantification of RNA splicing using LeafCutter. Nat. Genet. 50, 151–158.

Lorenz, R., Bernhart, S.H., Höner zu Siederdissen, C., Tafer, H., Flamm, C., Stadler, P.F., and Hofacker, I.L. (2011). ViennaRNA Package 2.0. Algorithms Mol. Biol. 6, 26.

Lundberg, S.M., and Lee, S.-I. (2017). A Unified Approach to Interpreting Model Predictions. In Advances in Neural Information Processing Systems 30, I. Guyon, U.V. Luxburg, S. Bengio, H. Wallach, R. Fergus, S. Vishwanathan, and R. Garnett, eds. (Curran Associates, Inc.), pp. 4765–4774.

Lundberg, S.M., Nair, B., Vavilala, M.S., Horibe, M., Eisses, M.J., Adams, T., Liston, D.E., Low, D.K.-W., Newman, S.-F., Kim, J., et al. (2018). Explainable machine-learning predictions for the prevention of hypoxaemia during surgery. Nat. Biomed. Eng. 2, 749–760.

Mackereth, C.D., Madl, T., Bonnal, S., Simon, B., Zanier, K., Gasch, A., Rybin, V., Valcárcel, J., and Sattler, M. (2011). Multi-domain conformational selection underlies pre-mRNA splicing regulation by U2AF. Nature 475, 408–411.

Manning, K.S., and Cooper, T.A. (2017). The roles of RNA processing in translating genotype to phenotype. Nat. Rev. Mol. Cell Biol. 18, 102–114.

Martin, M. (2011). Cutadapt removes adapter sequences from high-throughput sequencing reads. EMBnet.Journal 17, 10–12.

Martinez-Rucobo, F.W., Kohler, R., van de Waterbeemd, M., Heck, A.J.R., Hemann, M., Herzog, F., Stark, H., and Cramer, P. (2015). Molecular Basis of Transcription-Coupled Pre-mRNA Capping. Mol. Cell 58.

de la Mata, M., Alonso, C.R., Kadener, S., Fededa, J.P., Blaustein, M., Pelisch, F., Cramer, P., Bentley, D., and Kornblihtt, A.R. (2003). A Slow RNA Polymerase II Affects Alternative Splicing In Vivo. Mol. Cell 12, 525–532.

Maul-Newby, H.M., Amorello, A.N., Sharma, T., Kim, J.H., Modena, M.S., Prichard, B., and Jurica, M.S. (2021). A Model for DHX15 Mediated Disassembly of A-Complex Spliceosomes.

Mayer, A., di Iulio, J., Maleri, S., Eser, U., Vierstra, J., Reynolds, A., Sandstrom, R., Stamatoyannopoulos, J.A., and Churchman, L.S. (2015). Native elongating transcript sequencing reveals human transcriptional activity at nucleotide resolution. Cell 161, 541–554.

Mayerle, M., Raghavan, M., Ledoux, S., Price, A., Stepankiw, N., Hadjivassiliou, H., Moehle, E.A., Mendoza, S.D., Pleiss, J.A., Guthrie, C., et al. (2017). Structural toggle in the RNaseH domain of Prp8 helps balance splicing fidelity and catalytic efficiency. Proc. Natl. Acad. Sci. U. S. A. 542, 201701462–201701466.

Mercer, T.R., Clark, M.B., Andersen, S.B., Brunck, M.E., Haerty, W., Crawford, J., Taft, R.J., Nielsen, L.K., Dinger, M.E., and Mattick, J.S. (2015). Genome-wide discovery of human splicing branchpoints. Genome Res. 25, 290–303.

Meyer, M., Plass, M., Pérez-Valle, J., Eyras, E., and Vilardell, J. (2011). Deciphering 3′ss Selection in the Yeast Genome Reveals an RNA Thermosensor that Mediates Alternative Splicing. Mol. Cell 43, 1033–1039.

Nasim, F.H., Spears, P.A., Hoffmann, H.M., Kuo, H.C., and Grabowski, P.J. (1990). A Sequential splicing mechanism promotes selection of an optimal exon by repositioning a downstream 5’ splice site in preprotachykinin pre-mRNA. Genes Dev. 4, 1172–1184.

Niemelä, E.H., and Frilander, M.J. (2015). Regulation of gene expression through inefficient splicing of U12-type introns. RNA Biol. 11, 1325–1329.

Nojima, T., Gomes, T., Grosso, A.R.F., Kimura, H., Dye, M.J., Dhir, S., Carmo-Fonseca, M., and Proudfoot, N.J. (2015). Mammalian NET-Seq Reveals Genome-wide Nascent Transcription Coupled to RNA Processing. Cell 161, 526–540.

Nojima, T., Rebelo, K., Gomes, T., Grosso, A.R., Proudfoot, N.J., and Carmo-Fonseca, M. (2018). RNA Polymerase II Phosphorylated on CTD Serine 5 Interacts with the Spliceosome during Co-transcriptional Splicing. Mol. Cell 72, 369–379.e4.

Olthof, A.M., Hyatt, K.C., and Kanadia, R.N. (2019). Minor intron splicing revisited: identification of new minor intron-containing genes and tissue-dependent retention and alternative splicing of minor introns. BMC Genomics 20, 686.

Pai, A.A., Henriques, T., McCue, K., Burkholder, A., Adelman, K., and Burge, C.B. (2017). The kinetics of pre-mRNA splicing in the Drosophila genome and the influence of gene architecture. ELife 6, 1123.

Pandya-Jones, A., and Black, D.L. (2009). Co-transcriptional splicing of constitutive and alternative exons. RNA N. Y. N 15, 1896–1908.

Patel, A.A., McCarthy, M., and Steitz, J.A. (2002). The splicing of U12-type introns can be a rate-limiting step in gene expression. EMBO J. 21, 3804–3815.

Pedersen, T.L. (2019). patchwork: The Composer of Plots.

Pineda, J.M.B., and Bradley, R.K. (2018). Most human introns are recognized via multiple and tissue-specific branchpoints. Genes Dev. 32, 577–591.

Qin, D., Huang, L., Wlodaver, A., Andrade, J., and Staley, J.P. (2016). Sequencing of lariat termini in S. cerevisiae reveals 5’ splice sites, branch points, and novel splicing events. RNA N. Y. N 22, 237– 253.

Quinlan, A.R., and Hall, I.M. (2010). BEDTools: a flexible suite of utilities for comparing genomic features. Bioinformatics 26, 841–842.

Ray, D., Kazan, H., Cook, K.B., Weirauch, M.T., Najafabadi, H.S., Li, X., Gueroussov, S., Albu, M., Zheng, H., Yang, A., et al. (2013). A compendium of RNA-binding motifs for decoding gene regulation. Nature 499, 172–177.

Reimer, K.A., Mimoso, C.A., Adelman, K., and Neugebauer, K.M. (2021). Co-transcriptional splicing regulates 3’ end cleavage during mammalian erythropoiesis. Mol. Cell 81, 998–1012.e7.

Saldi, T., Cortazar, M.A., Sheridan, R.M., and Bentley, D.L. (2016). Coupling of RNA Polymerase II Transcription Elongation with Pre-mRNA Splicing. J. Mol. Biol. 428, 2623–2635.

Saldi, T., Riemondy, K., Erickson, B., and Bentley, D.L. (2021). Alternative RNA structures formed during transcription depend on elongation rate and modify RNA processing. Mol. Cell 81, 1789–1801.e5.

Shao, C., Yang, B., Wu, T., Huang, J., Tang, P., Zhou, Y., Zhou, J., Qiu, J., Jiang, L., Li, H., et al. (2014). Mechanisms for U2AF to define 3′ splice sites and regulate alternative splicing in the human genome. Nat. Struct. Mol. Biol. 21, 997–1005.

Slowikowski, K. (2020). ggrepel: Automatically Position Non-Overlapping Text Labels with “ggplot2.”

Smith, T.S., Heger, A., and Sudbery, I. (2017). UMI-tools: Modelling sequencing errors in Unique Molecular Identifiers to improve quantification accuracy. Genome Res. gr.209601.116.

Sontheimer, E.J., Sun, S., and Piccirilli, J.A. (1997). Metal ion catalysis during splicing of premessenger RNA. Nature 388, 801–805.

Sousa-Luís, R., Dujardin, G., Zukher, I., Kimura, H., Carmo-Fonseca, M., Weldon, C., Proudfoot, N.J., and Nojima, T. (2021). POINT technology illuminates the processing of polymerase-associated intact nascent transcripts. Mol. Cell 81, 1935–1950.e6.

van Steensel, B., and Belmont, A.S. (2017). Lamina-associated domains: links with chromosome architecture, heterochromatin and gene repression. Cell 169, 780–791.

Taggart, A.J., DeSimone, A.M., Shih, J.S., Filloux, M.E., and Fairbrother, W.G. (2012). Large-scale mapping of branchpoints in human pre-mRNA transcripts in vivo. Nat. Struct. Mol. Biol. 19, 719–721.

Taggart, A.J., Lin, C.-L., Shrestha, B., Heintzelman, C., Kim, S., and Fairbrother, W.G. (2017). Large-scale analysis of branchpoint usage across species and cell lines. Genome Res. 27, 639–649.

Takahara, K., Schwarze, U., Imamura, Y., Hoffman, G.G., Toriello, H., Smith, L.T., Byers, P.H., and Greenspan, D.S. (2002). Order of Intron Removal Influences Multiple Splice Outcomes, Including a Two-Exon Skip, in a COL5A1 Acceptor-Site Mutation That Results in Abnormal Pro-α1(V) N-Propeptides and Ehlers-Danlos Syndrome Type I. Am. J. Hum. Genet. 71, 451–465.

Tanaka, E., Bailey, T.L., and Keich, U. (2014). Improving MEME via a two-tiered significance analysis. Bioinformatics 30, 1965–1973.

Testa, S.M., Disney, M.D., Turner, D.H., and Kierzek, R. (1999). Thermodynamics of RNA−RNA Duplexes with 2- or 4-Thiouridines: Implications for Antisense Design and Targeting a Group I Intron. Biochemistry 38, 16655–16662.

Tilgner, H., Nikolaou, C., Althammer, S., Sammeth, M., Beato, M., Valcárcel, J., and Guigó, R. (2009). Nucleosome positioning as a determinant of exon recognition. Nat. Struct. Mol. Biol. 16, 996– 1001.

Tilgner, H., Knowles, D.G., Johnson, R., Davis, C.A., Chakrabortty, S., Djebali, S., Curado, J., Snyder, M., Gingeras, T.R., and Guigó, R. (2012). Deep sequencing of subcellular RNA fractions shows splicing to be predominantly co-transcriptional in the human genome but inefficient for lncRNAs. Genome Res. 22, 1616–1625.

Toan, N.M., Marenduzzo, D., Cook, P.R., and Micheletti, C. (2006). Depletion effects and loop formation in self-avoiding polymers. Phys. Rev. Lett. 97, 178302.

Tremblay, B.J.-M. (2019). universalmotif: Import, Modify, and Export Motifs with R.

Turowski, T.W., Petfalski, E., Goddard, B.D., French, S.L., Helwak, A., and Tollervey, D. (2020). Nascent Transcript Folding Plays a Major Role in Determining RNA Polymerase Elongation Rates. Mol. Cell.

Van Nostrand, E.L., Pratt, G.A., Shishkin, A.A., Gelboin-Burkhart, C., Fang, M.Y., Sundararaman, B., Blue, S.M., Nguyen, T.B., Surka, C., Elkins, K., et al. (2016). Robust transcriptome-wide discovery of RNA-binding protein binding sites with enhanced CLIP (eCLIP). Nat. Methods 13, 508–514.

Van Nostrand, E.L., Freese, P., Pratt, G.A., Wang, X., Wei, X., Xiao, R., Blue, S.M., Chen, J.-Y., Cody, N.A.L., Dominguez, D., et al. (2020). A large-scale binding and functional map of human RNA-binding proteins. Nature 583, 711–719.

Veloso, A., Kirkconnell, K.S., Magnuson, B., Biewen, B., Paulsen, M.T., Wilson, T.E., and Ljungman, M. (2014). Rate of elongation by RNA polymerase II is associated with specific gene features and epigenetic modifications. Genome Res. 24, 896–905.

Wachutka, L., Caizzi, L., Gagneur, J., and Cramer, P. (2019). Global donor and acceptor splicing site kinetics in human cells. ELife 8, e45056.

Wagih, O. (2017). ggseqlogo: a versatile R package for drawing sequence logos. Bioinformatics 33, 3645–3647.

Wan, Y., Anastasakis, D.G., Rodriguez, J., Palangat, M., Gudla, P., Zaki, G., Tandon, M., Pegoraro, G., Chow, C.C., Hafner, M., et al. (2021). Dynamic imaging of nascent RNA reveals general principles of transcription dynamics and stochastic splice site selection.

Warnasooriya, C., Feeney, C.F., Laird, K.M., Ermolenko, D.N., and Kielkopf, C.L. (2020). A splice site-sensing conformational switch in U2AF2 is modulated by U2AF1 and its recurrent myelodysplasia-associated mutation. Nucleic Acids Res. 48, 5695–5709.

Werner, M.S., and Ruthenburg, A.J. (2015). Nuclear Fractionation Reveals Thousands of Chromatin-Tethered Noncoding RNAs Adjacent to Active Genes. CellReports 12, 1089–1098.

Wickham, H., Averick, M., Bryan, J., Chang, W., McGowan, L.D., François, R., Grolemund, G., Hayes, A., Henry, L., Hester, J., et al. (2019). Welcome to the tidyverse. J. Open Source Softw. 4, 1686.

Wilke, C.O. (2019). cowplot: Streamlined Plot Theme and Plot Annotations for “ggplot2.”

Wu, J.Y., and Maniatis, T. (1993). Specific interactions between proteins implicated in splice site selection and regulated alternative splicing. Cell 75, 1061–1070.

Wu, S., Romfo, C.M., Nilsen, T.W., and Green, M.R. (1999). Functional recognition of the 3′ splice site AG by the splicing factor U2AF 35. Nature 402, 832–835.

Wuarin, J., and Schibler, U. (1994). Physical isolation of nascent RNA chains transcribed by RNA polymerase II: evidence for cotranscriptional splicing. Mol. Cell. Biol. 14, 7219–7225.

Yee, B.A., Pratt, G.A., Graveley, B.R., Nostrand, E.L.V., and Yeo, G.W. (2019). RBP-Maps enables robust generation of splicing regulatory maps. RNA 25, 193–204.

Yeo, G., and Burge, C.B. (2004). Maximum entropy modeling of short sequence motifs with applications to RNA splicing signals. J. Comput. Biol. J. Comput. Mol. Cell Biol. 11, 377–394.

Zamft, B., Bintu, L., Ishibashi, T., and Bustamante, C. (2012). Nascent RNA structure modulates the transcriptional dynamics of RNA polymerases. Proc. Natl. Acad. Sci. U. S. A. 109, 8948–8953.

Zhang, S., Aibara, S., Vos, S.M., Agafonov, D.E., Lührmann, R., and Cramer, P. (2019). Picard toolkit. Broad Inst. GitHub Repos.

